# Computational enzyme design by catalytic motif scaffolding

**DOI:** 10.1101/2024.08.02.606416

**Authors:** Markus Braun, Adrian Tripp, Morakot Chakatok, Sigrid Kaltenbrunner, Celina Fischer, David Stoll, Aleksandar Bijelic, Wael Elaily, Massimo G. Totaro, Melanie Moser, Shlomo Y. Hoch, Horst Lechner, Federico Rossi, Matteo Aleotti, Mélanie Hall, Gustav Oberdorfer

## Abstract

Enzymes are broadly used as biocatalysts in industry and medicine due to their coverage of vast areas of chemical space, their exquisite selectivity and efficiency as well as the mild reaction conditions at which they operate. Custom designed enzymes can produce tailor-made biocatalysts with potential applications extending beyond natural reactions. However, current design methods require testing of high numbers of designs and mostly produce de novo enzymes with low catalytic activities. As a result, they require costly experimental optimization and high-throughput screening to be industrially viable. Here we present rotamer inverted fragment finder–diffusion (Riff-Diff), a hybrid machine learning and atomistic modelling strategy for scaffolding catalytic arrays in de novo proteins. We highlight the general applicability of Riff-Diff by designing enzymes for two mechanistically distinct chemical transformations, the retro-aldol reaction and the Morita-Baylis-Hillman reaction. We show that in both cases it is possible to generate catalysts exhibiting activities rivalling those optimized by in-vitro evolution, along with exquisite stereoselectivity. High resolution structures of six of the designs revealed an angstrom level of active site design precision. The design strategy can, in principle, be applied to any catalytically competent amino acid constellation. These findings enable the practical applicability of de novo protein catalysts in synthesis and shed light on fundamental principles of protein design and enzyme catalysis.

## Introduction

Natural enzymes have emerged as an indispensable tool for chemists, offering unparalleled precision and selectivity in chemical transformation^1^. However, identifying natural enzymes with the desired activity can require extensive amounts of resources and screening capabilities. Very recently, computational protein design enabled the creation of biocatalysts designed for a variety of reactions^2–4^. Yet, an often underemphasized limitation of current approaches to enzyme design is the low initial catalytic rates of the designed biocatalysts. Even with recent advancements^5^ the current paradigm for generating efficient biocatalysts is to compensate for low initial efficiencies with high-throughput screening and directed evolution^6^. While this approach is valuable and robust, it is not suited to create novel, proficient catalysts for chemical reactions that are not found in nature or are exceedingly difficult to assess through high-throughput screening. For this reason, it is necessary to advance our ability to design efficient enzymes one-shot. This, however, requires addressing current methodological shortcomings in functional design and prediction of enzymatic activity. Overcoming these challenges will ultimately deepen our understanding of the fundamental principles of enzyme catalysis.

De novo enzyme design strategies are commonly based on the transition-state model of enzyme catalysis^7^. The model proposes that functional groups in the active site accelerate chemical reactions by stabilizing the reaction’s transition state over the ground state. In accordance with this model, minimal active sites, called theozymes, can be constructed by placing amino acid functional groups in a stabilizing geometry around a transition state model^8,9^. In the first computationally designed enzymes, these theozymes were introduced into cavities of natural proteins, which could support the desired geometry^10–12^. The thus-designed enzymes successfully catalysed chemical transformations, some of which were considered new to nature at the time, but with low catalytic rate accelerations and often required extensive screening^11–13^. Among the main limiting factors of the approach was the accuracy with which the catalytic side chains could be placed within the active site. Successful designs exhibited less than 1 Å side chain RMSD between the geometry in the theozyme and a high-resolution experimental structure. Inactive designs, however, showed unproductive conformations^14–16^. The recent transformative successes in accurate structure prediction methods and protein design tools warrant another attempt to solve this problem^17,18^.

Since the first successful attempts, critical analysis of designed enzymes and their evolved variants revealed additional aspects of the enzyme design problem. One was the construction of catalytically potent theozymes^15,19,20^. This was demonstrated by early mutational studies of designed retro-aldolases, for which multiple catalytic interactions were programmed to stabilize the reaction’s transition states^12^. Of the programmed interactions, only the catalytic lysine buried in the hydrophobic binding pocket significantly contributed to catalysis. Directed evolution of one of these designs even replaced the initial catalytic machinery with a completely new catalytic tetrad in the final variant. The geometry of this experimentally evolved tetrad was unlike any of the calculated theozymes, which further hinted at the low quality of initial theozymes^21^. Furthermore, molecular dynamics simulations of retro-aldolases at intermediate steps during directed evolution demonstrated the importance of preorganization in catalytic arrays. The conformational populations of catalytic amino acids were increasingly rigidified towards catalytically productive conformations^21^. Conversely, conformational distributions of active site residues in designed Kemp eliminases were shown to discriminate between active and inactive designs^22–24^. In addition, backbone-to-side-chain compatibility, as well as atomic density inside an active site pocket, was found to be a predictor of activity for a class of recombined de novo-designed xylanases^25^.

In this study, we investigated whether efficient biocatalysts can be created *one-shot* by incorporating three distinct catalytic centers in de novo protein backbones. We developed a method called Riff-Diff (**R**otamer **i**nverted **f**ragment **f**inder–**Diff**usion) that focuses on the precise placement of backbone-side-chain fragments, the design precision of the given catalytic arrays, and the design of custom binding pockets. To test the effectiveness of our approach, we designed retro-aldol enzymes based on a previously reported, highly optimized catalytic tetrad^21^ and Morita-Baylis-Hillmanases starting from two distinct evolved active site constellations. Detailed biochemical, biophysical, and structural characterization, as well as molecular dynamics simulations, allowed us to test current hypotheses about the precision of the placement of active site atoms, and conformational dynamics in the designed enzymes. The extraordinary wealth of this data comes from the exact same catalytic environment placed into different backbones, allowing us to draw conclusions for de novo enzyme design that were not obvious before.

### Artificial motif libraries scaffold catalytic arrays

We set out to create Riff-Diff as a computational pipeline that designs enzymes from scratch when given an array of amino acids as input. Previous attempts at grafting catalytic arrays into proteins have shown that the achievable catalytic effect of an array depends on how precisely it can be reproduced in the designed active site^2,15,23,26^. We utilized RFdiffusion to scaffold the initial backbone around the input array and added modifications to improve precision. First, each amino acid of the input array is embedded into a helical fragment. The rotamers of each amino acid are selected to be compatible with the phi-psi angle combination of the fragment backbone^27^. This ensures minimal energetic penalty is associated with the preferential catalytic rotamer. We call the combination of the individual fragments an *artificial motif*.

The arrangement of helical fragments in an artificial motif depends on the positions of its functional group atoms and the chosen rotamers. This arbitrary arrangement of helical fragments can result in physically implausible motifs and low scaffolding success rates. To increase scaffolding success, we developed a script that generates libraries of artificial motifs from catalytic arrays from which high-quality motifs can be sampled. The script starts by inverting rotamers of the catalytic array, which fixes the functional groups in space and varies the position of their backbone atoms. Rotamers are selected for backbone compatibility with the helical fragments and subsequently placed onto the fragment’s backbone atoms. Next, a search for all possible combinations of fragments that are not sterically clashing with each other is performed. Non-clashing assemblies are ranked according to the rotamer probabilities of the individual active site residues and by their abundance in the protein data bank (PDB). The identified assemblies are aggregated and stored in an artificial motif library (Figure 1a). This artificial motif library is then used as input to RFdiffusion to scaffold enzyme backbones (Figure 1b).

**Fig. 1.**
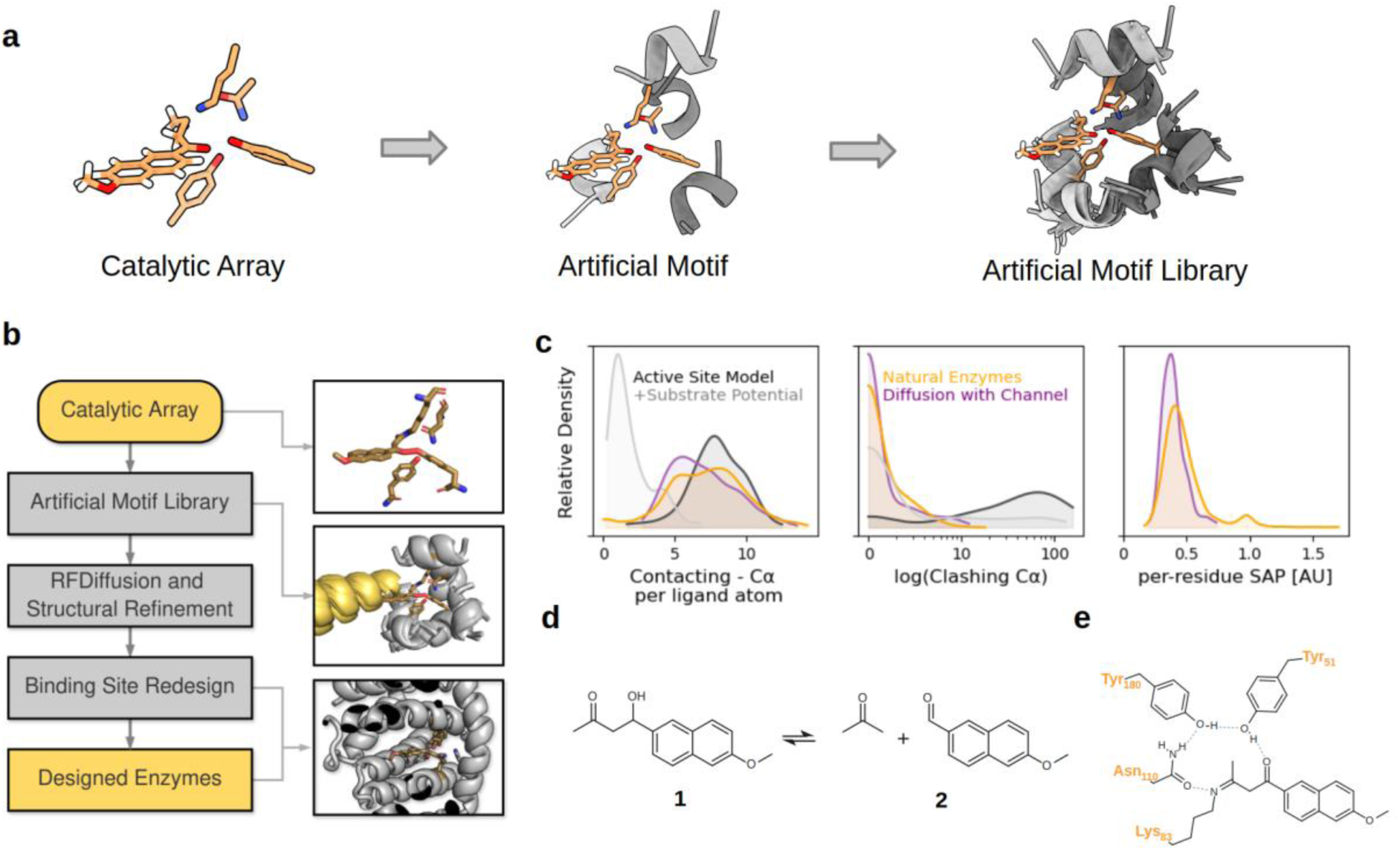
Riff-Diff scaffolds de novo enzymes starting from catalytic arrays. **a**, Artificial motif libraries are collections of artificial motifs that are constructed from arrays of side chains. **b,** Schematic overview of the semi-automated Riff-Diff pipeline. The channel placeholder helix is shown in yellow. **c,** Binding pockets of natural enzymes (yellow) typically bury their substrates, measured by the number of alpha-carbons within 8 Å of the substrate. RFdiffusion’s substrate potential offers only a trade-off between substrate burial and clashes (overlaps) with the pocket (light- and dark gray). Riff-Diff (purple) scaffolds enzyme backbones that bury the substrate in binding pockets reminiscent of natural enzymes. The surface aggregation propensity (SAP, right panel) of the designed enzymes is similar to that of natural enzymes. **d,** A retro-aldol cleavage transforms methodol (**1**) to 6-methoxy-2-naphthaldehyde (**2**) and acetone. **e,** Four residues complete the catalytic array to catalyse the retro-aldol cleavage. The lysine is shown in the covalently linked state to the mechanistic inhibitor 1-(6-methoxynaphthalen-2-yl)butane-1,3-dione, taken from (PDB: 5AN7).

### Ensuring accessible and highly optimized active site pockets

An essential feature of enzymes as biocatalysts is their substrate specificity, which is dictated by well-defined interactions of the substrate with the enzyme’s asymmetric binding pocket. To analyse how deep substrates are typically buried within binding pockets, we extracted an enzyme dataset from the PDBbind database^28^. In this dataset, an average of 7.1 α-carbons were within 8 Å of the substrate, normalized by the number of ligand-heavy atoms. We used this metric to evaluate ligand burial in the substrate pockets of backbones scaffolded by RFdiffusion, which offers an auxiliary potential to mimic the physical presence of a substrate. We found that this potential decreased the number of clashes between ligand and backbone but failed to promote the formation of well-defined substrate binding pockets. This means that using the substrate auxiliary potential resulted in a trade-off between clashes and substrate interactions in the scaffolded artificial motifs. For this reason, we implemented a novel approach to enforce pockets during diffusion by adding another alpha-helix as an entry-channel placeholder to each artificial motif at the position of the binding pocket. A custom auxiliary potential then places the centre of the denoising trajectory on the helix and enforces a distance constraint of all backbone atoms to this centre (see Methods). This ensures that the generated binding pockets exhibit characteristics mimicking those of natural enzymes (Figure 1c). The ‘placeholder’ helix is removed after diffusion, leaving a vacant binding pocket.

### Riff–Diff robustly scaffolds catalytic arrays

An integral part of Riff-Diff is its backbone refinement protocol in which the diffused enzyme backbones are refined iteratively. Recent work showed that the quality of de novo backbone-side-chain pairs improves when predicted structures are used for subsequent sequence design^29,30^. Therefore, we implemented iterative refinement cycles into Riff-Diff during which the coordinates of idealized helical fragments of the artificial motifs were used as constraints to minimize the motif backbone RMSD. In the initial version of Riff-Diff, Rosetta’s FastDesign protocol^31^ explicitly optimized the packing of active site residues and ligand interactions within the substrate binding pocket based on the geometry specified in the input catalytic array. To sample efficiently, FastDesign could only exchange amino acids within the active site and was constrained at each position to residue identities that were assigned high probabilities by ProteinMPNN^32^. In an updated version of Riff-Diff, we exchanged ProteinMPNN and FastDesign with LigandMPNN and FastRelax, reducing the computational effort of the refinement protocol. The generated sequences were predicted using ESMFold^33^ and the highest-scoring structures were fed into the next refinement cycle. We chose ESMFold to predict structures throughout the refinement cycles because of its faster inference compared to AlphaFold2. By default, five cycles of refinement are performed in a design trajectory.

Following backbone refinement, an additional script utilized the CoupledMoves protocol^34^ to refine ligand interactions with the binding pocket. ProteinMPNN was used with its soluble weights^35^ to redesign the remaining protein backbone, generating the final enzyme sequences. In the final evaluation step, these sequences were predicted with AlphaFold2 and ranked with a combination of metrics for structure quality and active-site positioning (see Methods). We combined the scripts for generating artificial motif libraries, backbone diffusion and refinement, and CoupledMoves into a single method for the semi-automated design of enzymes, called Riff-Diff (Figure 1b). Riff-Diff requires only minimal user input to generate sequences that scaffold a given catalytic array. The code, alongside detailed instructions for use, can be found at https://github.com/mabr3112/riff_diff_protflow. Detailed information on filtering steps and cutoffs is provided in the Supplementary Methods.

### Design of de novo retro-aldolases

With the ability to accurately build de novo scaffolds with binding pockets around arbitrary catalytic arrays, we investigated whether this approach produces proficient enzymes starting from an existing, potent catalytic array. As our model system, we selected the catalytic tetrad of an artificial retro-aldolase called RA95.5-8F (PDB: 5AN7^21^), which catalyses the retro-aldol cleavage of (*R*)-methodol (**1**) into 6-methoxy-2-naphthaldehyde (**2**) and acetone (Figures 1d and 1e). This catalytic tetrad emerged after extensive rounds of laboratory evolution and its amino acids were analysed in mutational, mechanistic and computational studies. The fluorescently active cleavage product **2** enables sensitive detection of catalytic activity, making this an ideal model system for designed enzymes.

Sequences for this tetrad designed by the Riff-Diff pipeline were generally predicted to fold as designed. Metrics for AlphaFold2 pLDDT, active-site RMSD and Rosetta energy for the predicted structures can be found in the supplementary materials (Supplementary Figure S1). To select designs for experimental testing, we ranked the pool of designed enzymes predominantly by the side chain RMSD of the catalytic tetrad. After additional manual inspection, we excluded designs for which the binding pocket appeared inaccessible to the substrate. The final selection consisted of 36 sequences derived from 12 unique backbones generated from 8 unique artificial motifs, named RAD1 to RAD36 (Supplementary Table S1). The backbones adopted novel folds, having an average maximum TM-score of 0.49 when searched with FoldSeek^36^ against the PDB. In addition, no similar sequences could be found for any of the 36 sequences in a protein-protein BLAST search against the non-redundant protein sequence database^37^. The set of 36 designed retro-aldolases were ordered as synthetic genes, of which 35 could be cloned. An initial screen for expression and activity revealed that all 35 designs expressed soluble and 32 (91%) variants exhibited activity towards *rac*-methodol above the negative control, with seven variants exhibiting rates of product formation several orders of magnitude higher than the average conversion within the screen. (Supplementary Figure S2).

### Detailed characterization and analysis of de novo retro-aldolases

To investigate the origins of the design’s divergent catalytic rates, we purified them from large-scale expressions and assessed folding, kinetic parameters, and investigated catalytic contributions of the tetrad amino acids. Gel filtration analysis showed monodisperse peaks corresponding to monomeric species (Figure 2a) for all designs and intact mass spectrometry validated the identity of the purified enzymes (Supplementary Table S2). Circular dichroism (CD) spectroscopy also confirmed their helical architecture (Supplementary Figure S3). In addition, we collected small-angle X-ray scattering (SAXS) data to evaluate the overall folding of our enzymes in solution. To judge whether a protein roughly adopted the designed structure, we used Dimensionless Kratky plots (Supplementary Figure S4), computed the expected radius of gyration, and calculated χ² values for the fits between predicted and measured scattering data (Supplementary Table S3). Overall, SAXS measurements could confirm the expected fold for 29 of 35 designs and expression data from batch fermentation confirmed all constructs to express as soluble, alpha-helical monomers (Figure 2b). Next, we measured Michaelis Menten kinetics, allowing us to compare catalytic performance to previous designs and the highly evolved variant RA95.5-8F.

**Fig. 2.**
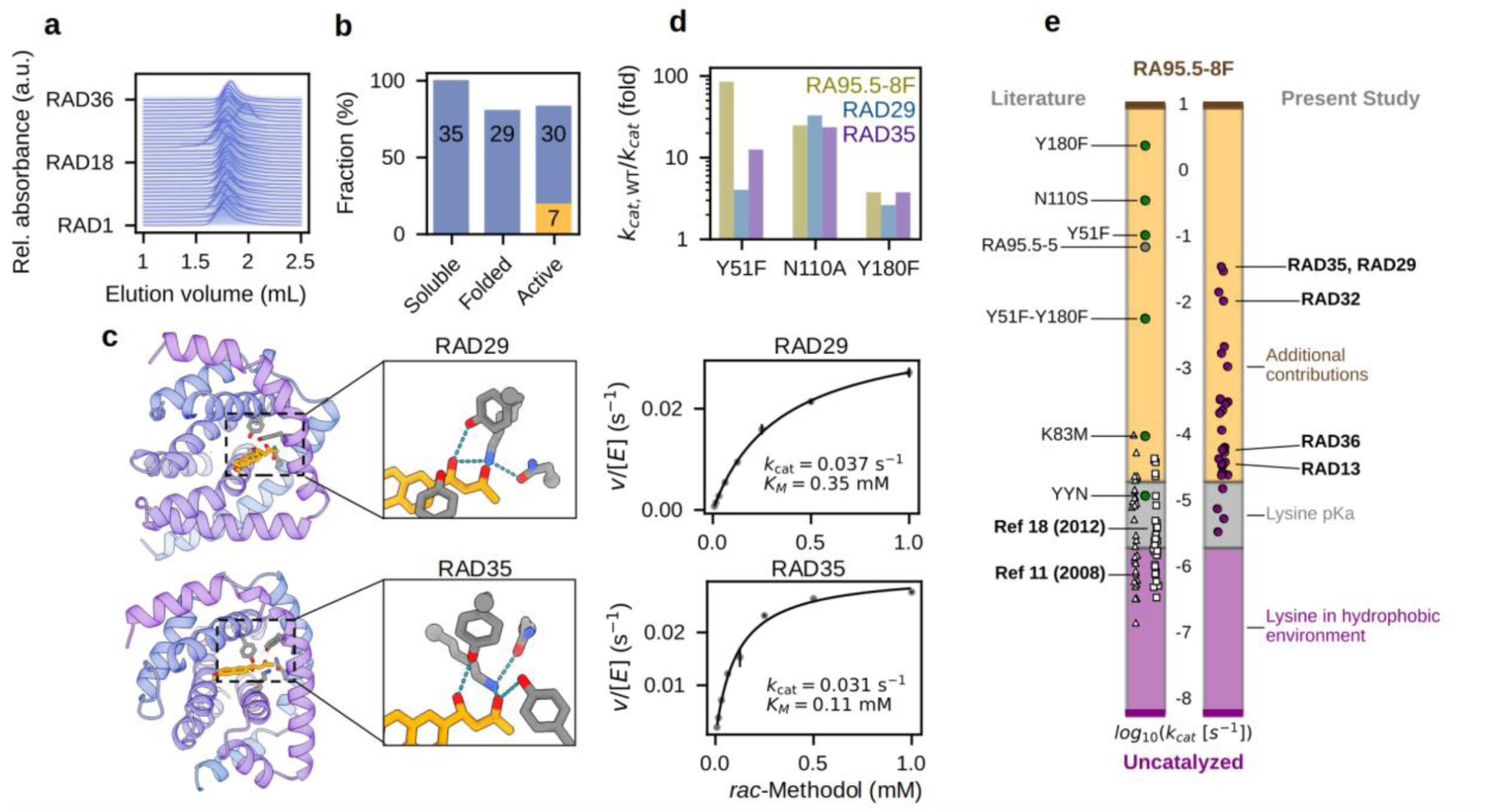
Activity of designed retro-aldol enzymes exceeded those of previous *one-shot* designs. **a**, All retro-aldolases eluted at the volume corresponding to the monomeric peak, as evidenced by size exclusion chromatography (SEC). SEC traces were normalized and stacked. **b,** 29 of 35 designed retro-aldolases were correctly folded according to SAXS (FoXS ꭓ² < 5). For 30 designs, product formation in the initial activity screen exceeded the background reaction. Seven designs exhibited *k*_cat_ > 10^-3^ s^-1^ (yellow bar). **c,** RAD29 and RAD35 exhibited the highest activity among the designed retro-aldolases. The left panels depict the design’s structures in complex with (*R*)-methodol predicted by AlphaFold3. **d,** Site-directed mutagenesis confirmed the participation of the tetrad residues in the catalytic mechanism. The residue numbering refers to the position of the catalytic residues in the model tetrad, not in the final designs. For asparagine-to-alanine variants, we compare the decrease in activity to RA95.5-8F N110S. **e,** For most enzymes, rate accelerations exceeded catalytic effects provided by the lysine in the hydrophobic pocket alone. Left bar: White triangles and squares represent computationally designed enzymes of previous design campaigns. Green dots represent variants that emerged after directed evolution. Right bar: computationally designed retro-aldolases of this study.

RA95.5-8F cleaves methodol at a maximum velocity (*k*_cat_) of 10 s^-1^ at 29 °C, which is more than 10^9^-fold faster than the uncatalyzed reaction, occurring at 6.5 x 10^-9^ s^-1^ ^19,21^. The artificial motifs used in this study were constructed from the tetrad only, leaving out conformational effects of other first- and second-shell residues. Accordingly, we expected the maximum *k*_cat_ of our designs to be lower than the reported rate of RA95.5-8F. We determined Michaelis Menten constants for 30 of the 35 produced retro-aldolases (Supplementary Figure S5, Supplementary Table S4), the catalytically most proficient variants were RAD35 and RAD29 (Figure 2c). They catalysed the cleavage of *rac-*methodol with a *k*_cat_ of 3.6 x 10^-2^ s^-1^ and 3.1 x 10^-2^ s^-1^, respectively, a roughly 5 x 10^6^-fold rate acceleration over the uncatalyzed reaction. Thus, our method produced enzymes orders of magnitude faster than previous computationally designed retro-aldolases, surpassing e.g. the activity of the evolved catalytic antibody 38C2 (*k*_cat_ 1.1 x 10^-2^ s^-1^)^38^. A table of catalytic rates of past designed and evolved retro-aldolases was added to the supplementary materials for reference (Supplementary Table S5). Of particular note is the high affinity of RAD29 for *rac-* methodol. With a *K*_M_ of ∼100 µM, its catalytic efficiency of 290 M^-1^ s^-1^ approaches that of the extensively evolved RA95.5-5 (320 M^-1^ s^-1^)^38^. The improved catalytic rates over previous computationally designed retro-aldolases confirmed that efficient enzymes can be designed with Riff-Diff when starting from potent catalytic arrays. We next sought to investigate to which extent the tetrad residues participated in catalysis.

During the first steps of the reaction, the tetrad’s lysine attacks the carbonyl group of methodol to form a high-energy, hemiaminal intermediate (Supplementary Figure S6). The central role of this lysine to catalysis in our enzymes was corroborated by the diminishing activities upon its substitution to alanine (Supplementary Figure S2). For 22 variants, the catalytic rate (< 10^-3^ s^-1^) falls within the range achievable by an isolated lysine in a hydrophobic pocket (Figure 2e)^18,19^. Seven designs exceeded this range, suggesting their tetrad residues might participate in the catalytic mechanism. To gain further insights, we determined Michaelis Menten parameters for variants targeting the tetrad residues of our most active designs RAD29 and 35 (Supplementary Figure S5). The substitutions reduced activity, although to a lesser extent than for substitutions in RA95.5-8F (Figure 2d). For RA95.5-8F the strongest effect on catalysis was observed when substituting Tyr51, which acts as a base catalysing the retro-aldol cleavage step. The effect on catalysis of exchanging the corresponding tyrosine in RAD29 and RAD35 was less pronounced.

The original tetrad requires its residues to occupy distinct protonation states over the course of the reaction. The catalytic lysine must be deprotonated for the initial nucleophilic attack and Tyr51 serves as both an acid and a base in different steps (Supplementary Figure S6). As a consequence, the reaction rate depends on the pH of its environment. We determined the pH-rate dependence for several RAD variants and found a range of apparent pKa_1_ values between 7 and 9.0 (Supplementary Figure S7). This is higher than the apparent pKa_1_ of the original tetrad (pKa_1_ = 6.2), but close to the pKa determined for the variant RA95.5-8 (pKa_1_ = 8.0). Importantly, the pH rate profiles of RAD35 and RAD29 revealed that their kinetic parameters were determined below their pH optimum, leading to underestimated kinetic parameters. Nevertheless, the pH-rate curves for RAD29 and RAD35, in combination with the results from site-directed mutagenesis, provide evidence for the participation of the tetrad residues in catalysis.

### RAD29 and RAD35 are highly stable and stereoselective biocatalysts

An enzyme’s catalytic rate is one among several metrics that define its utility as a biocatalyst. Other crucial factors include stereoselectivity, substrate scope, the total number of turnovers the enzyme can perform, and its stability in organic solvents or at elevated reaction temperatures. De novo designed proteins are known for their high thermal stability, offering an advantage over natural enzymes. CD melting curves confirmed that all but one of the designed retro-aldolases remained folded to >90 °C (Figure 3a). To further investigate the design’s thermodynamic stability, we measured chemical denaturation midpoints with guanidinium hydrochloride (GdnHCl). Twenty of 33 enzymes displayed cooperative unfolding with mid-points of denaturation ranging from 2.5 to > 6.5 M GdnHCl (Figure 3b, Supplementary Figure S8), indicating a wide range of thermodynamic stability, with RAD29 and RAD35 being among the most stable ones. Making use of this finding, we next investigated whether computational metrics could predict the measured denaturation midpoints of our enzymes. Indeed, we found statistically significant (p < 0.05) correlations for prediction confidence by AlphaFold2 (average pLDDT), the total score of the Rosetta Energy Function, core atomic density, and Rosetta’s surface aggregation propensity (SAP) score (Supplementary Figure S9). Using these metrics, we built a simple linear regression model that predicted chemical denaturation midpoints with a Pearson R of 0.8 (Figure 3c).

**Figure 3:**
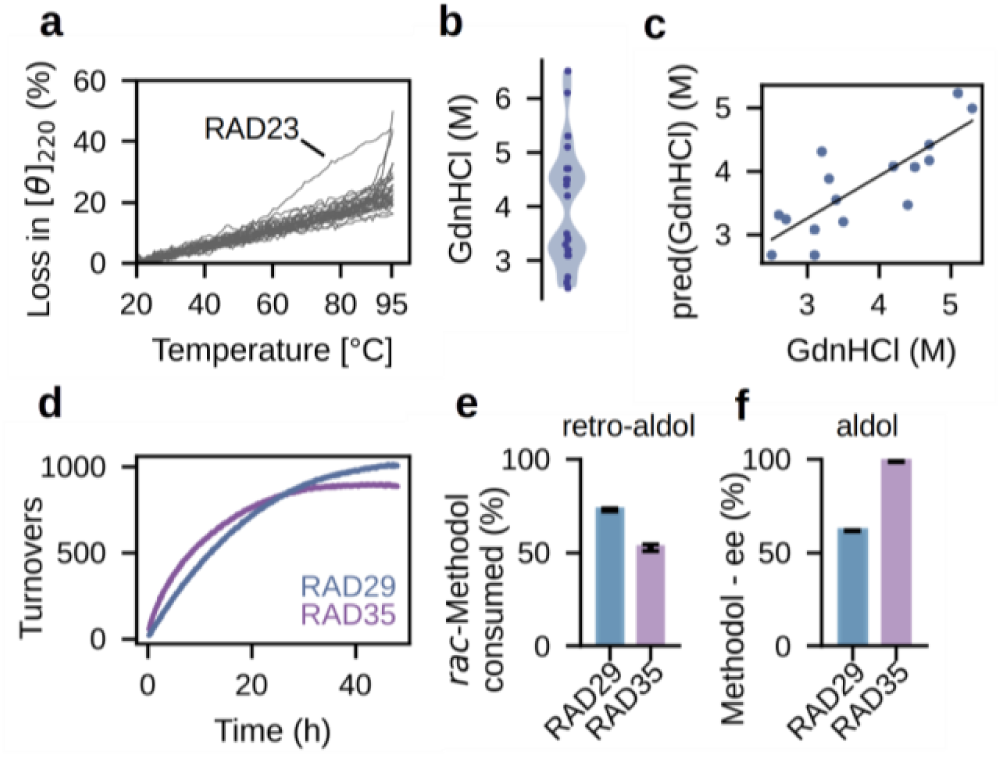
Designed retro-aldolases are highly stable, enantioselective and can catalyse multiple turnovers. **a**, CD melting curves confirmed high thermodynamic stability. Except for RAD23, only negligible loss in signal intensity at 220 nm can be observed up to 95 °C. **b,** According to CD experiments, chemical denaturation midpoints for 20 of 35 constructs ranged from 2.5 to >6 M of guanidinium hydrochloride (GdnHCl). **c**. A linear regression model based on computational design metrics (Rosetta total score, average Alphafold2 pLDDT, surface aggregation propensity and core contacts) can predict chemical denaturation midpoints with a Pearson R of 0.8. **d**, RAD29 and RAD35 can catalyse 1000 and 895 turnovers, respectively. **e**, Consumption of *rac*-methodol after 24h reaction time, leaving 30% unreacted (*S*)-**1** with 77% ee (RAD29) and 52.5% unreacted (*S*)-**1** with 88% ee (RAD35). **f**, RAD29 and RAD35 exhibit stereoselectivity toward (*R*)-**1**. Bars correspond to the measured ee values (%) of product (*R*)-**1** at 5% substrate conversion in the aldol direction.

Determination of the turnover number (TON) of RAD29 and RAD35 resulted in 1000 and 895 turnovers after reacting for 48 hours in the retro-aldol direction (Figure 3d). Stereoselectivity was assessed by product formation in the aldol direction - the condensation of **2** with acetone to methodol. While RAD29 was modestly stereoselective with an E-value of 4 toward the formation of (*R*)-methodol (60% ee), RAD35 displayed exquisite (*R*)-stereoselectivity and E-value >200 toward (*R*)-methodol (99% ee, Figure 3e). These values were corroborated by the analysis of the kinetic resolution of *rac*-methodol, with RAD29 and RAD35 displaying selectivity also in the retro-aldol reaction. Under optimized reaction conditions, RAD29 and RAD35 converted 70% and 47.5% of the racemic substrate (Figure 3f). The high stereoselectivity, turnover and high thermal stability make RAD35 a promising biocatalyst toward application under process-relevant conditions^39^.

### Crystal structures reveal unexpected relationship between structure and function

We obtained crystal structures of the four variants RAD13 (*k*_cat_ 3.8 x 10^-5^ s^-1^), RAD17 (*k*_cat_ n.d.), RAD32 (*k*_cat_ 1.1 x 10^-2^ s^-1^), and RAD36 (*k*_cat_ 6.2 x 10^-5^ s^-1^). The variant’s divergent catalytic rates are ideal for investigating how much of the variation can be explained by side chain packing in the active site. The crystal structures are in agreement with the design models, having backbone C_α_ RMSDs between 0.68 Å and 1.2 Å (Figure 4a). The catalytic residues of RAD13 and RAD17 adopted the correct conformation with heavy atom side chain RMSDs of 0.42 Å and 1.09 Å to the design model and 0.89 Å and 1.2 Å to the catalytic geometry, respectively (Figure 4b). Despite the near-atomic precision with which the tetrad was scaffolded in both variants, RAD13 and RAD27 were more than 10^5^ fold slower than RA95.5-8F. This indicates that precise positioning of catalytic side chains is a valuable design objective but by itself not sufficient to design efficient biocatalysts, even when scaffolding a catalytic array which is known to be potent.

**Figure 4.**
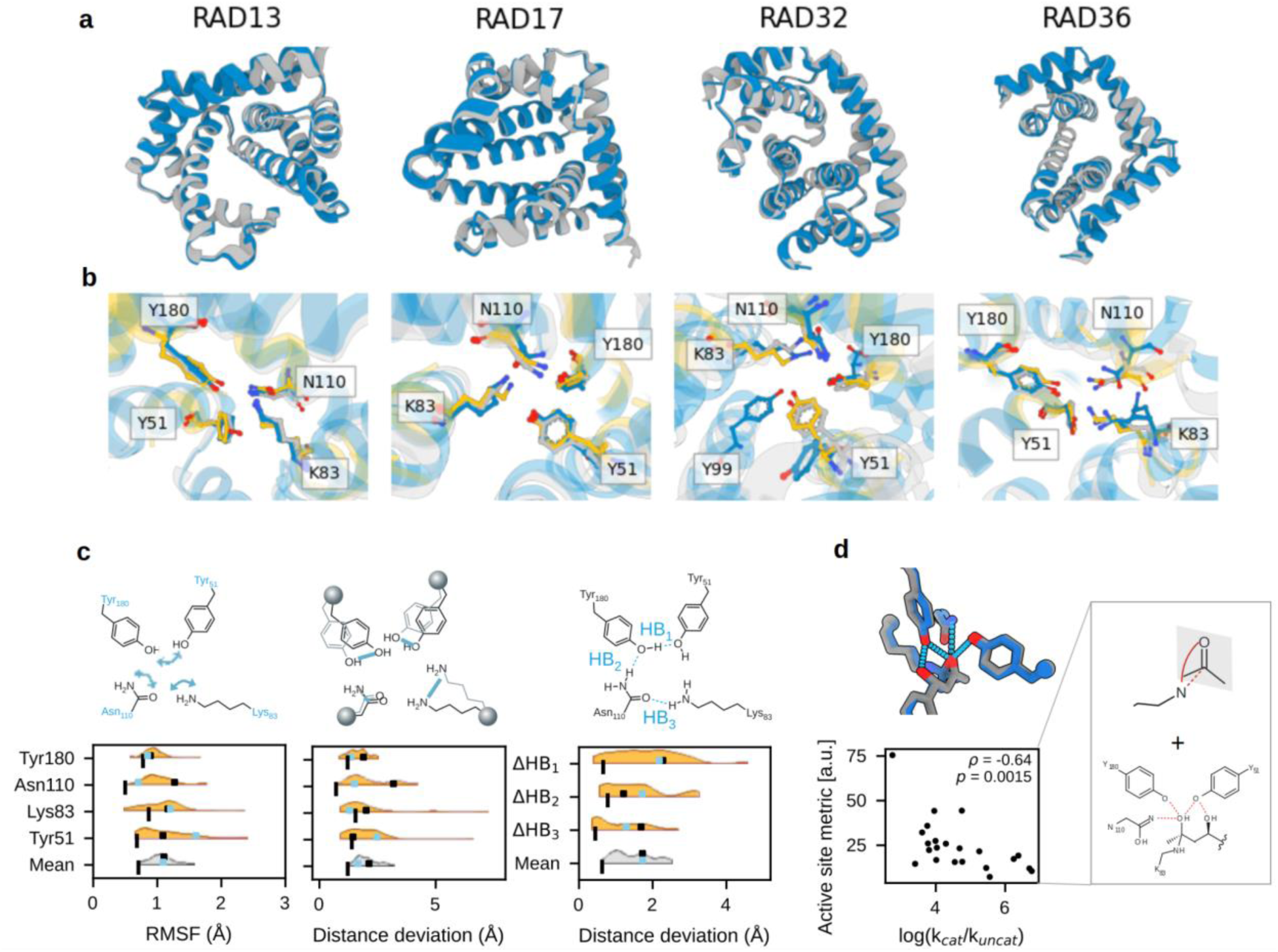
Crystal structures of RAD constructs reveal high accuracy of the scaffolded catalytic tetrad. **a,** The backbones of the design models (gray) are almost identical to the experimentally obtained crystal structures (blue), displaying overall C_α_ RMSDS below 1.2 Å. PDB IDs: 9GBT, 9FW5, 9FW7, 9FWA. **b,** Active site residues in the crystal structures (blue) agree well with those in the design models (gray) and the tetrad (yellow). In the crystal structure of RAD32, the intended position of the tyrosine hydroxyl group is assumed by another tyrosine residue that was not part of the design model. Multiple conformations of the catalytic lysine residue are visible in the crystal structure of RAD36. The conformer with the highest occupancy is adopting a catalytically incompetent orientation. **c,** Three panels showing individual metrics of active site rigidity. The root mean square fluctuation (RMSF) of all catalytic residues during MD simulations (left), measured heavy-atom distances of the model to the theozyme throughout the MD trajectories (middle) and the resulting difference in H-bonding distances between the catalytic residues (right). While the measured values are small, it can be seen that the designed active sites were still more flexible and less positioned than the original tetrad on average. Vertical black lines correspond to RA95.5-8F, black squares to RAD29, and blue squares to RAD35. Amino acid indices indicate positions in RA95.5-8F. **d**, A composite metric calculated from complex predictions of AlphaFold3 correlated with activity. The composite metric combines interaction distances of the tetrad functional groups to the hemiaminal intermediate hydroxyl groups, and the positioning of the substrate carbonyl group in the enzyme-substrate complex.

In the active site of RAD36, we observed density for three distinct conformations of the catalytic lysine. One of the three lysine conformations points away from the active site, partially explaining its low catalytic rate. In contrast, in the active site of RAD32, the most active design for which an experimental structure could be obtained, a tyrosine of the catalytic tetrad (Tyr120) adopts a catalytically unproductive conformation. This tyrosine is supposed to act as a catalytic base, promoting C-C bond cleavage. However, Tyr99, another tyrosine in the RAD32 active site, can compensate for this interaction by placing its deprotonated hydroxyl group close to the position of the methodol β-hydroxyl group in the model (Figure 4b). Exchanging either of the two tyrosines for phenylalanine reduced activity by over tenfold (Figure 2d). This loss of activity confirms the participation of Tyr99 in the catalytic machinery. However, the decreased activity of RAD32 Y120F contrasts the observed unproductive conformation in the RAD32 crystal structure. The unproductive conformation could simply be an artifact of crystal packing or could indicate a conformational flip induced by substrate binding.

### Active site dynamics reveal uncoupling of precision and positioning

Flexibility in catalytic side chains can reduce their likelihood of adopting active conformations, thereby slowing catalysis. Such high conformational flexibility was suggested as one reason for low activity in previously designed enzymes^40,41^. Molecular Dynamics (MD) simulations supported this idea and demonstrated how initially flexible side chains of designed enzymes rigidified in subsequent directed evolution campaigns^42,43^. Similarly, we computed short MD replicates of our designs in the apo state to compare their active site dynamics to those of RA95.5-8F, using RMSF as a metric to estimate side chain flexibility. To evaluate the agreement between the simulated conformations and the catalytic geometry, we calculated two additional metrics: (1) the distance deviation of the hydrogen bonding functional groups, and (2) the distance deviation of the active-site functional groups after superimposition on the tetrad backbone atoms. Consistent with the literature, the active site of RA95.5-8F was rigid and remained positioned for catalysis throughout the trajectories. In contrast, all designed active sites were less consistently positioned than the original tetrad, except for isolated residues (Figure 4c). In five designs, the catalytic lysine even drifted from the active site into a non-catalytic state. The generally more pronounced flexibility and alternative conformations of the active sites in the designed enzymes offer a further explanation for their lower activity compared to RA95.5-8F. However, neither of the calculated metrics from the MD simulations correlated with activity, suggesting that the difference between designs might be linked to other factors.

To address the active site’s interactions with the bound substrate during the reaction, we used AlphaFold3 to model enzyme-substrate complexes directly. We predicted the designs complexed with (*R*)-methodol to assess the accessibility of the carbonyl group to the catalytic lysine and further modelled the hemiaminal intermediate with a covalent link to the catalytic lysine. The predictions revealed the geometry of several catalytically relevant interactions: (1) the near-attack conformation of the lysine to the carbonyl group, (2) the three hydrogen bonds to the alpha hydroxyl group, and (3) Tyr51 positioned between the substrate’s two hydroxyl groups to shuttle protons. Figure 4d depicts the predicted interactions for RA95.5-8F. Across all designs, we observed that substrate positioning and hydrogen bond distances of catalytic residues around the hemiaminal intermediate correlated with experimental activity. Notably, an active-site metric derived from these features exhibited a Spearman ρ of -0.64 with measured catalytic rates (Figure 4d), outperforming apo-state metrics such as side chain RMSD. This highlights the possibility of generating predictive models from de novo design datasets that capture catalytic interactions for ranking and selecting designed enzymes.

### Probing the proficiency of catalytic arrays using a non-biological reaction

To demonstrate the general applicability of Riff-Diff to design useful enzymes, we selected the Morita-Baylis-Hillman (MBH, Figure 5a) reaction as a secondary model system. In organic synthesis, this reaction - the carbon-carbon coupling of activated alkenes with aldehydes to yield functionalized allylic alcohols - typically is resource intensive and requires high loading of nucleophilic catalysts such as 1,4-diazabicyclo[2.2.2]octane (DABCO), 4-dimethylaminopyridine (DMAP) or imidazole and usually needs prolonged reaction times^44–46^. Though low levels of promiscuous MBH activity have been reported for several proteins^47–49^, there are no natural enzymes known to date that catalyse this transformation efficiently. Computational design of MBHases based on a streptavidin cofactor and cysteine or histidine residues as nucleophiles have resulted in constructs with low activity^13,50^. However, directed evolution campaigns of one of these designs have revealed several highly active variants^51,52^.

**Figure 5:**
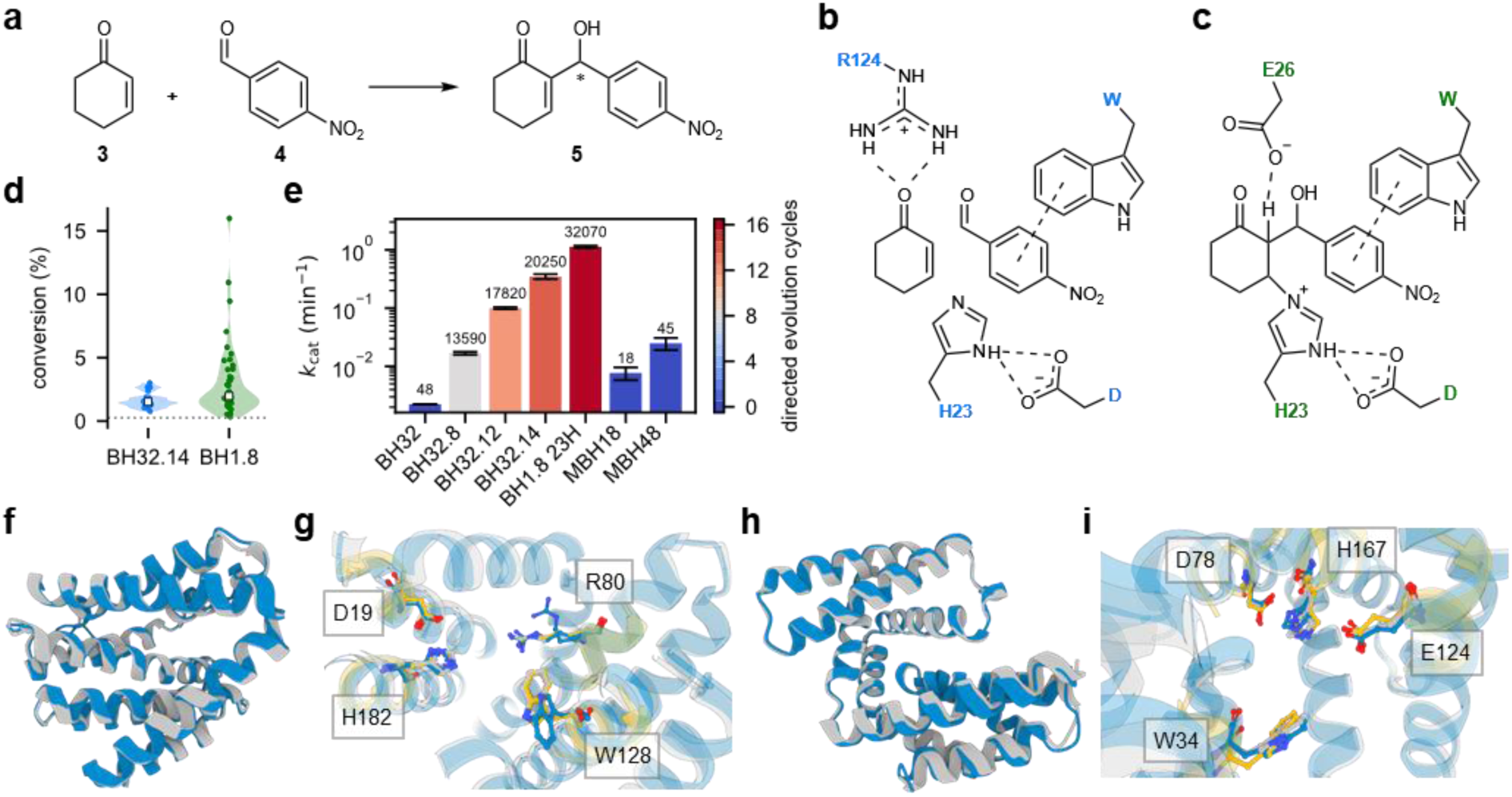
De novo enzymes for the MBH reaction are active and agree with the design models. **a,** Reaction scheme for the Morita-Baylis-Hillman (MBH) reaction between 2-cyclohexenone (**3**) and 4-nitrobenzaldehyde (**4**) to yield 2-(hydroxy(4-nitrophenyl)methyl)cyclohex-2-en-1-one (**5**). **b** and **c,** Catalytic arrays based on transition state 1 from BH32.14 (**b**) and transition state 3 from BH1.8 (**c**). **d,** Conversion of substrates **3** and **4** after 8 hours at 2 mol% catalyst loading for designs based on the active sites of BH32.14 and BH1.8. The dotted line marks the background reaction with lysozyme. **e,** The catalytic constant of MBH48 outperforms that of BH32.8, a variant that emerged after 8 rounds of directed evolution. In BH1.8 23H, the noncanonical amino acid Nδ-methylhistidine was substituted with a regular histidine residue. Numbers above bars indicate the total number of screened clones - for evolved variants, this does not necessarily correspond to the number of unique sequences that were screened. **f,** The crystal structure (blue) of MBH2 (PDB 9QDP) agrees with the design model (gray), showing a C_α_ RMSD of 0.53 Å. **g,** In the active site of MBH2, the tryptophan side chain in the crystal structure (blue) adopts a different conformation compared to the design model (gray) and the input geometry (yellow). Additionally, the catalytic arginine residue is present in two different conformations, one pointing away from the active site. **h,** The crystal structure of MBH48 (blue, PDB 9R7F) is in high agreement with the design model, exhibiting a C_α_ RMSD of 0.78 Å. **i,** Active site residues in the crystal structures of MBH48 (blue) agree well with those in the design model (gray) and the input geometry (yellow).

We used two active site constellations from evolved MBH variants as show-case examples. The active site of BH32.14, an enzyme that has emerged after 14 rounds of directed evolution, features a histidine residue (His23) acting as a nucleophile, forming a covalent bond to the activated alkene substrate. To position His23 and prime it for nucleophilic attack, a glutamic acid (Glu46) is hydrogen-bonding to the histidine Nε. Importantly, an arginine residue (Arg124) is believed to play a crucial role in the stabilization of negative charges on three oxyanion intermediates at the C1 and C3 position that occur throughout the reaction trajectory and thus needs to adopt different rotameric states (Supplementary Figure S10). The second active site was taken from BH1.8, the MBHase with highest activity to date. This variant has emerged after eight additional rounds of directed evolution upon substitution of the nucleophilic histidine in BH32.8, an evolutionary predecessor of BH32.14, with the noncanonical amino acid Nδ-methylhistidine. According to the proposed mechanism, a glutamic acid residue (Glu26) with an unusually high predicted pKa of 8.1 stabilizes the negative charge on the C3 alkoxide in intermediate 2 and mediates a rate-limiting proton transfer step from the C2 proton to the C3 alkoxide (transition state 3). In BH32.14, the proton transfer is facilitated by a water molecule. In the final step of the reaction, the bond between the intermediate and His23 is eliminated to generate the product **5**.

We used Riff-Diff to create designs based on both active sites (see Methods). The geometry of His23 and Arg124 as well as the substrates’ orientations were extracted from a previously reported density-functional theory (DFT) model of transition state 1 from the BH32.14-catalyzed reaction of cyclohexenone (**3**) with 4-nitrobenzaldehyde (**4**). These geometries were then used as input for the Riff-Diff pipeline. As the reported essential residue Glu46 was not present in the DFT model, we modelled a carboxylic acid interacting with the histidine Nδ-hydrogen using XTB^53^. An additional Trp side chain forming π–π-stacking interactions with **4** was also added to facilitate better substrate binding using XTB and CREST^54^. We selected 18 sequences (MBH1 to 18) derived from 14 unique backbones for experimental characterization based on this active site constellation (Figure 5b). For designs modelled after the active site of BH1.8, we extracted the coordinates of the Nδ-methylhistidine, Glu26 and the substrates from transition state 3 of quantum mechanics/molecular mechanics (QM/MM) calculations of the BH_MeHis_1.8-catalyzed reaction of **3** with **4** and combined them with the previously generated carboxylic acid and tryptophan orientations to create a catalytic array (Figure 5c). Forty-five sequences (MBH19 to 63) derived from 23 unique backbones were selected for experimental testing (see Methods for details). Sequences and computational metrics of ordered designs can be found in the supplementary materials (Supplementary Table S6, Supplementary Figure S11).

Endpoint assays showed that 17 of the designs based on BH32.14 (94%) and 42 of the designs based on BH1.8 (93.3%) formed product above background level and outperformed the small molecule nucleophile catalysts imidazole and DMAP, emphasizing the robustness of the Riff-Diff design approach (Supplementary Table S7). While median conversions in both design sets are similar, constructs with highest conversion are based on BH1.8 (Figure 5d).

We determined Michaelis Menten parameters for MBH18 and MBH48, the constructs showing highest conversion after 8 hours in their respective active site set (Supplementary Figure S12). MBH18 displays a *k*_cat_ of 7.7 x 10^-3^ min^-1^, making MBH18 roughly 3.5-fold more active than BH32 (*k*_cat_ of 2.2 x 10^-3^ min^-1^), the computationally designed starting point for directed evolution. A considerable amount of side product formation - a competing aldol reaction product according to similar previously reported HPLC retention times^51,52^ - can be observed for this construct (Supplementary Figure S13). With a *k*_cat_ of 0.025 min^-1^ MBH48 activity is 1.5 times higher than in BH32.8 (*k*_cat_ of 0.0168 min^-1^), a variant that emerged after screening 13590 clones in 8 rounds of directed evolution (Figure 5e). Additionally, by-product formation in MBH48 is minimal (Supplementary. Figure S13). We suspect that high apparent Michaelis constants of both MBH18 (*K*_M_ of 8.4 mM and 6.2 mM for **3** and **4**, respectively) and MBH48 (*K*_M_ of 25.2 mM and 1.4 mM for **3** and **4**, respectively) are likely due to the challenging bimolecular nature of the MBH reaction, complicating the design of binding pockets with high affinity for two substrates. This is also reflected in the apparent Michaelis constants of the evolved variants BH32.14 (*K*_M, **3**_: 2.56 mM, *K*_M, **4**_: 1.14 mM) and BH1.8 (*K*_M, **3**_: 12.02 mM, *K*_M, **4**_: 323 µM). A table comparing Michaelis Menten parameters for previously reported designed and evolved enzymes for the MBH reaction is included in the supplementary materials (Supplementary Table S8).

### Structural characterization of MBH designs

Experimental structure determination resulted in a 1.13 Å resolution crystal structure for MBH2 (PDB 9QDP), a construct ranking among the designs with lowest conversion. The structure is in good agreement with the design model, showing an overall backbone C_α_ RMSD of 0.53 Å; the catalytic side chains generally superimpose well with the catalytic geometry except for Trp140, which adopts a flipped conformation (side chain RMSD 0.54 Å excluding Trp140, Figures 5f and g). This flip leads to a wider entrance channel and ultimately abolishes the designed π–π-stacking interactions with the substrate **4**. In addition, Arg92 could be modelled in two conformations: one resembling the design model, the other pointing away from the active site. Taken together, these observations provide a possible reason for the low conversion rates of MBH2.

Structural investigation of MBH48 via CD experiments revealed the expected helical structures and showed the onset of unfolding at ∼85 °C (Supplementary Figure S14). The crystal structure of MBH48 (PDB 9R7F) displays an overall backbone C_α_ RMSD of 0.78 Å compared to the design model (Figure 5h). In addition, the active site residues superimpose well with the input geometry, resulting in a side chain RMSD of 1.06 Å between the crystal structure and the desired active site geometry (Figure 5i).

### Conclusions and Outlook

Even though exciting de novo design results were reported recently^5^, current enzyme design and engineering rely on high-throughput screening methods to produce viable enzymes. While highly valuable, this method conflicts with an original goal of the field, which was to produce catalysts for reactions that are difficult to optimize via high-throughput screens. Our study constitutes a significant step towards this goal. Using the semi-automated Riff-Diff method, we demonstrated that scaffolding of catalytic arrays yields active enzymes while screening as little as 35 designed sequences. We successfully employed this method to create de novo enzymes for two distinct chemical transformations, the retro-aldol and Morita-Baylis-Hillman reactions. Thorough biochemical characterization allowed us to identify correctly folded designs that exhibited vastly divergent catalytic rates and substrate affinities. Detailed analysis of the catalytic efficiency of the enzymes was supplemented with rich dynamic data obtained from MD simulations and crystal structures for four (RADs) and two (MBHs) of the designed biocatalysts. The resulting datasets enabled us to detail reasons for higher and lower catalytic proficiencies and are an ideal test set to benchmark models of catalytic activity. We emphasize that de novo design campaigns like these for a diverse set of enzyme catalysed reactions will complement the analysis of natural enzymes towards building general models of enzyme catalysis.

An essential component of the high activity of our designs were the potent catalytic centres we used as starting points. For novel reactions, such potent catalytic centres are likely not available and they will have to be created from scratch using methods such as DFT calculations. With the ability to scaffold catalytic centres in arbitrary geometries - regardless if naturally or computationally derived - we can now systematically evaluate their catalytic potency in controlled environments of de novo backbones that were previously inaccessible. This study offers an example of this, with crystal structures of four retro-aldol enzymes that have different backbones but harbour the same catalytic motif and two structures of MBHases with distinct catalytic motifs. The structures allowed us to conclude that designing for active site precision alone is not sufficient to produce proficient enzymes. Our findings support the notion that designing enzymes with activities similar to their natural counterparts will require accounting for catalytic interactions and their conformational dynamics throughout the complete reaction cycle.

Next to the catalytic centres themselves, the prediction of catalytic activity remains a bottleneck for *one-shot* enzyme design. Protein language models represent a new frontier for this task, but their poor generalizing abilities currently limit their effectiveness in novel chemical reactions, especially with de novo enzymes. Physics-based methods like QM/MM appear more suited to handle generalizability but they are computationally costly and require highly accurate structural data as inputs. This severely limits their predictive utility. An alternative route for physics-based activity prediction is conformational ensembles of the active-site. MD simulations conducted in this study revealed populations of active site conformations that were unattainable by structure prediction methods or crystal structures alone. The comparably broader distributions of active site conformations, observed in the conformational ensembles of the designed retro-aldolases, offer one explanation for their lower activity compared to the evolved enzyme. This highlights how computational tools that optimize an active site’s conformational ensemble could yield future improvements in activity of designed enzymes. Additionally, our datasets show how generic catalytic contributions within the active site must be considered in predictive activity models, as observed with the catalytic tyrosine at an alternative position in the crystal structure of RAD32. Thus, the development of computationally cheap estimates for conformational populations and generic models of catalytic contributions are critical for future improvements in physics-based prediction of enzymatic activity.

Finally, enzyme design and engineering campaigns have played a pivotal role in biocatalysis to create superior catalysts with tailor-made properties to manufacture new pharmaceuticals and complex fine chemicals. Yet very few of these studies started from scratch and all those that did relied on directed evolution or screened several hundred de novo sequences to reach practically useful activities in the designed biocatalysts. RiffDiff offers a way to go beyond screening campaigns and help transform enzyme design from an orphan field into an approach that the broader biotechnological community can apply.

## Methods

### Cloning, protein production and purification

RAD genes were obtained from IDT, and MBH genes were obtained from GenScript. The amino acid sequences can be found in Supplementary Table S1. All genes were cloned into a vector containing a hexa histidine-tag and TEV-cleavage site using Golden Gate assembly with a BsaI restriction enzyme^55^. The sequence of the plasmid can be found at https://doi.org/10.5281/zenodo.15494922. All enzymes used during DNA amplification, mutagenesis and cloning were sourced from New England Biolabs. For the initial screen, the plasmids were transformed into *E. coli* BL21 (DE3) STAR cells using standard procedure (30 minutes incubation on ice, followed by 30 seconds heat shock at 42 °C and regeneration in SOC medium at 37 °C). A 96-well plate containing 500 µL of LB medium with 50 µg/mL kanamycin in each well was inoculated with a single colony for each construct, including a positive control, a negative control and a sterile control. We used a recent de novo retro-aldolase developed in our lab as positive control^56^, and a modified natural protein without enzymatic activity as negative control. Sequences for positive and negative control are in the supplementary materials. Some wells were left empty to serve as sterile controls and blank samples for subsequent analysis. Cultures were grown overnight at 37 °C and 250 rpm. 100 µL of these cultures were used to inoculate 24-well plates containing 3 mL of ZY autoinduction medium. For each sample, two replicates were performed. The cultures were grown for 6 hours at 37 °C and 200 rpm, afterwards the temperature was decreased to 18 °C and shaking continued for another 18 hours.

Samples were harvested by centrifugation and resuspended in lysis buffer (20 mM sodium phosphate (NaPi), 500 mM NaCl, 1% n-Octyl-β-D-glucopyranoside, 20 µg/mL DNase I, 250 µg/mL lysozyme, one cOMPLETE protease inhibitor pill/100 mL, pH 7.4). Chemical lysis was performed at RT for one hour on a vibrating plate at 1000 rpm. The lysate was centrifuged for 20 minutes at 4800 rcf and the supernatant was purified in a 96-well plate using magnetic Ni-NTA beads and an Opentrons OT-2 pipetting robot. The lysate was incubated with the magnetic beads for 20 minutes at RT, and afterwards, the supernatant was discarded. Beads were washed with 200 µL wash buffer (20 mM NaPi, 500 mM NaCl, 2 mM tris(2-carboxyethyl)phosphine (TCEP), pH 7.4) once. The supernatant was discarded, and bead-bound proteins were eluted by adding 100 µL elution buffer (20 mM NaPi, 500 mM NaCl, 250 mM imidazole, 2 mM TCEP, pH 7.4) twice per well, yielding 200 µL eluate for each replicate. Protein concentration was determined via Bradford assay using bovine serum albumin in a concentration range of 0.05 to 0.65 mg/ml as a calibration. Protein purity was confirmed using SDS-PAGE.

For the generation of active site lysine to alanine variants, primers were designed using the NEBasechanger web interface. PCR reactions containing 10 µL of 5X NEB Q5 buffer, 32.5 µL ddH2O, 1 µL of dNTP mix (10 mM in ddH2O), 2.5 µL of forward and reverse primers (10 µM in ddH2O), 1 µL of template DNA (2 ng/µL) and 0.5 µL Q5 DNA polymerase were set up. Amplification was performed for 25 cycles, with a denaturation temperature of 98 °C, an annealing temperature according to the used primer pairs and an extension temperature of 72 °C. Initial denaturation and final extension steps were performed for 30 s and 2 min, respectively. A complete list of primers and annealing temperatures used can be found in Supplementary Table S9. Following PCR, to 1 µL of the PCR reaction 1 µL each of T4 DNA ligase buffer, T4 DNA ligase, T4 polynucleotide kinase and DpnI as well as 5 µL of ddH2O were added and incubated at 22 °C for 1h. Plasmids were transformed into *E. coli* TOP10 cells and plated on agar plates containing kanamycin. Plasmids from single colonies were isolated using a Monarch Spin Plasmid Miniprep kit and sent for sequencing. Screening of lysine-to-alanine variants was performed in the same way as the original designs.

For batch production, 10 mL of TB medium containing 100 mg/L kanamycin were inoculated with a single colony of BL21 (DE3) STAR cells containing the respective plasmid. Cultures were grown overnight at 37 °C and 140 rpm on a shaking incubator. 10 mL of overnight culture was used to inoculate 1 L of TB medium containing 100 mg/L kanamycin. Cultures were grown to an OD_600_ of 0.6 to 0.8 at 37 °C and 140 rpm and induction was initiated by adding isopropyl β-D-thiogalactopyranoside (IPTG) to a final concentration of 0.1 mM. Cells were harvested 4-5 hours after induction via centrifugation at 4000 rcf. For MBH19 to MBH63, cultures were placed in a shaking incubator at 20 °C and 140 rpm overnight following induction and harvested the next morning. Pellets were washed once with 30 mL 0.9% NaCl solution at RT and stored at -20 °C.

Pellets were thawed and resuspended in lysis buffer (20 mM NaPi, 500 mM NaCl, and a spatula of DNase I and lysozyme per 200 mL, pH 7.4). The suspensions were sonicated for 15 min on ice, afterwards the lysate was centrifuged at 43 000 rcf for 40 min. The supernatant was loaded onto gravity columns containing 1-2 mL nickel immobilized metal affinity chromatography (Ni-IMAC) resin equilibrated with lysis buffer and washed with wash buffer (see above). The purified proteins were eluted using an elution buffer (see above). Buffer was exchanged to storage buffer (20 mM NaPi/300 mM NaCl/2 mM TCEP pH 7.4 for RAD designs, 20 mM NaPi/150 mM NaCl pH 7.4 for MBH designs) using centrifugal filters. For RAD designs, His-tag cleavage was performed via addition of 0.062 mg of TEV protease (produced in-house) per mg of protein and incubation at 4°C overnight. The cleaved tag was removed using reverse Ni-IMAC. MBH His-tags were not removed. The final purification step consisted of gel filtration on a S75 Increase 10/300 GL or S75 10/300 column equilibrated with the respective storage buffer. Protein concentrations were determined by specific absorbance at 280 nm and by BioRAD assay. Samples were flash-frozen in liquid nitrogen and stored at -80 °C.

### Circular Dichroism

Circular Dichroism (CD) and thermal denaturation experiments for retro-aldolase constructs were performed on a JASCO-1500 CD-spectrophotometer in 10 mM NaPi buffer pH 7.4 containing 150 mM sodium fluoride (NaF) and 0.5 mM TCEP. Sample concentration was set to approximately 0.25 mg/mL. Spectra were recorded in 1 mm quartz cuvettes with cap. Thermal denaturation was performed at 3 °C/min while monitoring CD signal intensity at 220 nm. At 20, 45, 70 and 95 °C spectra (190 to 260 nm) were recorded. Additional spectra were recorded after cooling down to 20°C. Each spectrum consisted of three accumulations.

Chemical denaturation experiments were performed in 100 mM NaPi buffer pH 7.4 containing 300 mM NaF and 1 mM TCEP at a final protein concentration of 0.6 mg/mL. Guanidine hydrochloride (GdnHCl) from a 7.4 M stock solution was added to final concentrations between 0 M and 7.1 M. The concentration of the stock solution was determined using a refractometer. Protein samples were incubated at RT overnight in the respective buffer and the CD signal at 220 nm was recorded in quartz capillaries. Denaturation midpoints were calculated from a sigmoidal fit with the Python SciPy library.

MBH48 CD spectra and thermal denaturation experiments were performed on an Applied Photophysics Chirascan V100 CD-spectrophotometer in a 20 mM NaPi buffer containing 150 mM NaF at a concentration of 0.1 mg/mL. Spectra were recorded in a 1 mm quartz cuvette with cap. Thermal denaturation was performed at 1.5 °C/min, recording full spectra at every degree from 25 °C to 95 °C and an additional spectrum after returning to the starting temperature.

### SAXS

SAXS profiles were recorded at the ESRF BM29 BioSAXS beamline^57^. Samples were prepared in 20 mM NaPi buffer pH 7.4 containing 150 mM NaCl, 1 mM TCEP and 3% glycerol. Protein concentration was set between 2 and 4 mg/mL. Blank measurements were performed using buffer from the flowthrough of centrifugal filters for each sample.

The individual frames for every sample were analysed manually using the ATSAS software package^58^. Frames were averaged and the buffer signal was subtracted. All parameters listed in Supplementary Table 2 were calculated using autorg/autognom functions in ATSAS. The scattering profiles were trimmed at low q values according to the autorg suggestion and a fit of the computational models as well as the available crystal structures to the experimental scattering profiles was performed using FoxS with offset and explicit hydrogens enabled, considering q values up to 0.5. C1 and C2 parameters were set to be flexible between 0.99 to 1.05 and -2 to 4, respectively.

### X-ray crystallography

Crystallization drops were set up with commercial crystallization screens using the vapor diffusion method employing a mosquito® Xtal3 crystallization robot (SPT Labtech) and incubated at 293 K. The protein concentration varied between 10-30 mg/ml in 20 mM NaPi/150 mM sodium chloride (NaCl)/1 mM TCEP pH 7.4 (RADs) or 20 mM HEPES/150 mM NaCl pH 7.4 (MBHs). The drop volume was 400nL, with a 1:1 protein and precipitant solution ratio. Crystallization drops were equilibrated against a reservoir containing 40 µL of precipitant solution. Crystals of RAD32 and MBH2 were obtained from manually set-up drops of 2 µL with 80 µL of precipitant solution in the reservoir. Depending on the construct, crystals appeared after one day to two weeks. Successful crystallization conditions and diffraction parameters are summarized in Supplementary Table S10. The obtained crystals were harvested from mother liquor with CryoLoops (Hampton Research) and briefly incubated with mother liquor containing 25% glycerol or 25% PEG400 followed by flash freezing in liquid nitrogen. Diffraction data was collected at 100 K on ESRF beamlines, Grenoble (France). Full datasets (360°) were collected to 2.43 Å (RAD13), 2.9 Å (RAD17), 2.0 Å (RAD32), 1.73 Å (RAD36), 1.13 Å (MBH2) and 1.93 Å (MBH48) resolution.

The collected data were processed using XDS^59^ with the provided input file from the beamline. Data resolution cutoffs were determined by pairef^60^. Structure determination was performed by molecular replacement using PHASER^61^ with the design models as search templates. The best solution was refined in reciprocal space with PHENIX^62^ with 5% of the data used for R_free_ and by real-space fitting steps against σA-weighted 2Fo–Fc and Fo–Fc electron density maps using COOT^63^. Water molecules were placed automatically into difference electron density maps and accepted or rejected according to geometry criteria and their B-factors, defined in the refinement protocol. For RAD17, feature-enhanced maps^64^ were generated to reduce noise and enhance existing weak features, thereby facilitating the interpretation and modelling of side chains throughout the entire structure but especially those of the active site residues (Supplementary Figure S15).

### Activity Measurements

Catalytic activity in the retro-aldol reaction was determined by following the formation of the aldehyde product **2** via measuring fluorescence emission at 452 nm at an excitation wavelength of 330 nm. Reactions were carried out in a reaction buffer 1 containing 20 mM NaPi, 300 mM NaCl, and 1 mM TCEP in 5 vol% dimethyl sulfoxide (DMSO), pH 7.4 at a 200 µL reaction volume in a 96 well-plate format at a temperature of 29 °C. Measurements for Michaelis Menten kinetics were measured with serial dilutions starting at a concentration of 1 mM *rac-*methodol. This resulted in eight concentrations of 1, 0.5, 0.25, 0.13, 0.063, 0.031, 0.016, and 0.0078 mM. Parameters *k*_cat_ and *K*_M_ were determined by fitting reaction velocity and substrate concentration in a Michaelis Menten model using the Python library SciPy, version 1.13.1.

pH profiles for RAD constructs were determined at 50 µM methodol concentration in 384 well plates at a total sample volume of 100 µL and a temperature of 25 °C. Final enzyme concentrations were set to 2 µM (RAD13), 12.6 µM (RAD17), 0.03 µM (RAD29), 0.06 µM (RAD32), 0.03 µM (RAD35) and 3.6 µM (RAD36). To 10 µL of protein samples in 10 mM NaPi/300 mM NaCl pH 7.4 90 µL of reaction buffer 2 at various pH values (100 mM phosphate/100 mM borate/100 mM acetate buffer containing 55.56 µM methodol and 5.56 vol% DMSO) were added. The reaction buffer 2 pH was adjusted to values between 4.6 and 10.6 in 0.4 steps. *k*_cat_ / *K*_M_ was determined via monitoring of fluorescence emission as described above. For each measurement, the background reaction at the respective pH was subtracted. pKa values were obtained according to a fit with a two-pKa model (equation 1) using the Python library SciPy, version 1.15.2.

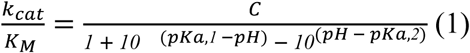

*k*_cat_ and *K*_M_ have their usual meaning, C corresponds to the maximum value of *k*_cat_ / *K*_M_, and pKa_1_ and pKa_2_ are the logarithmic acid dissociation constants.

The aldol addition reaction was performed using a final enzyme concentration of 5 μM (for both RAD29 and RAD35)in reaction buffer 3 (20 mM NaPi, 300 mM NaCl, pH 7.4). 5 mM of **2** was added from a stock solution in acetonitrile (MeCN, 15 vol% final concentration co-solvent) followed by 5 vol% acetone. The reactions (final volume 500 μL) were incubated for 24 h at 30 °C and 120 rpm. Control reactions were run in the absence of enzyme. All reactions were performed in triplicates. The products were extracted using ethyl acetate (2 x 250 μL) spiked with 0.5 vol% acetophenone as internal standard (IS). The combined extracts were dried over anhydrous sodium sulfate and analysed by chiral-phase HPLC as indicated below. The results were corrected for the buffer-catalysed background reaction under the same conditions. The identity of the compounds was further confirmed by GC/MS and the use of authentic reference material. Substrate consumption was measured using a calibration curve.

For conversion and substrate ee determination in the retro-aldol reaction, a final enzyme concentration of 20 µM (for both RAD29 and RAD35) in reaction buffer 3 was used to catalyse the reaction with 2 mM *rac*-**1** as the substrate, added from a stock solution in MeCN (15 vol% final concentration). The samples were incubated for 24 hours at 30°C and 120 rpm. All experiments were performed in triplicates. The reaction was analysed using the same method as the forward aldol reaction, monitoring the consumption of both isomers and the formation of the aldehyde product **2**.

RAD TON experiments were performed at 0.1 µM final enzyme concentration and 2 mM final substrate concentration (*rac*-**1**) in reaction buffer 3 containing 15 vol% DMSO. Samples were incubated for 48 hours at 29°C. All experiments were performed in triplicates. Product formation was monitored using fluorescence and comparison to a calibration curve. The background reaction of substrate without enzyme was subtracted.

Conversion screening of MBH constructs was performed at an enzyme concentration of 100 µM. Reactions were performed in 96-microwell plates in 20 mM phosphate buffer pH 7.4 with 150 mM NaCl and 10% DMSO at substrate concentrations of 25 mM **3** and 5 mM **4**. Samples were taken after 8 hours of incubation at 40 °C and 800 rpm by quenching 10 µL of the reaction mixture with 10 µL of MeCN and subsequently used for HPLC analysis.

Michaelis Menten kinetics for the reaction of **3** with **4** were recorded at 60 µM (MBH48) and 90 µM (MBH18) concentration in 20 mM phosphate buffer pH 7.4 with 150 mM NaCl and 10% DMSO at 100 µL reaction volume in polypropylene 96-microwell plates. Reactions versus **3** were performed at a fixed concentration of **4** (5 mM) and **3** concentrations ranging from 0.5 mM to 32 mM. Reactions versus **4** were performed at a fixed concentration of **3** (25 mM) and **4** concentrations ranging from 0.1 mM to 6.4 mM. Reaction progress was sampled every 50 min (MBH48) or 2 hours (MBH18) by quenching 10 µL of the reaction mixture with 10 µL of MeCN and subsequently analysed by HPLC. All reactions were performed in triplicates. For MBH48, biological replicates were performed as well.

Initial reaction velocities (V_0_) at each substrate concentration were determined from linear fits of conversion versus time. The combined V_0_ versus **3** and V_0_ versus **4** were fitted globally using a random order binding model (equation 2):

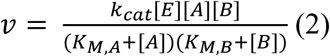

where *k*_cat_ is the catalytic constant, [E] is the total enzyme concentration, [A] and [B] are the initial **3** and **4** concentrations, respectively, and *K*_M,A_ and *K*_M,B_ are the corresponding apparent Michaelis constants. Fits were performed with shared *k*_cat_, *K*_M,A_ and *K*_M,B_ values as a three-dimensional surface fit using the Python library SciPy, version 1.15.2. Fits for individual measurements versus **3** and **4** are shown in Supplementary Figure S12.

Conversion and kinetic measurements for MBH constructs were analysed using a ThermoScientific UltiMate 3000 HPLC equipped with a Gemini SecurityGuard 4x2.0 cartridge and a Kinetex 5 µm XB-C18 100 Å column (50 × 2.1 mm, Phenomenex) at a flow rate of 1 mL/min. An isocratic method using 22 vol% MeCN in water at 20 °C was used for all measurements. The 2-(hydroxy(4-nitrophenyl)methyl)cyclohex-2-en-1-one (**5**) product elution volume and concentration were determined via comparison to a calibration curve prepared from a chemically synthesized standard.

### Molecular Dynamics simulations

For all molecular dynamics simulations reported in this study, gromacs version 2023.4 was used. The GRO coordinates of the design models and the topology files were generated using the pdb2gmx tool with the amber99sb-ildn force field. The MD unit cell was defined as a dodecahedron with a minimum distance of 1.0 nm between the complex and the box edges using the gmx editconf tool. The system was solvated with the SPC/E water model using the gmx solvate tool and neutralized by adding Na^+^ ions using the gmx grompp and gmx genion tools with a salt concentration of 150 mM. The system was then subjected to standard energy minimization and equilibration using the gmx grompp and gmx mdrun tools. Energy minimization was performed using the steepest descent algorithm with a maximum force of 1000 kJ*mol^−1^*nm^−1^ and a maximum of 5000 steps. The equilibration step consisted of two phases: NVT (constant number of particles, volume, and temperature) and NPT (constant number of particles, pressure, and temperature). The NVT phase was run for 100 ps with a temperature coupling of 298 K using the v-rescale thermostat. The NPT phase was run for 200 ps with a pressure coupling of 1 bar using the Parrinello-Rahman barostat. Backbone Ca RMSDs were followed throughout the trajectories to evaluate simulation stability (Supplementary Figure S16).

Prior to the 20 ns replicate simulations, individual 50 ns simulations were run to confirm proper equilibration of the 36 designs and RA95.5-8F. The parameters for the 50 ns and 20 ns production runs of all models were as follows: the temperature and pressure were set at 298 K and 1 bar by the v-rescale thermostat and Parrinello-Rahman barostat, respectively; hydrogen bonds were constrained using the LINCS algorithm; the Verlet cut-off scheme was used to process intra-atomic interactions; the PME method was implemented to account for Coulombic and Lennard-Jones interactions; and a van der Waals cut-off radius of 1.0 was applied. The simulations were analysed using the MDAnalysis python package version 2.8.0^65^.

### Implementation of the placeholder helix to scaffold substrate pockets

The artificial motifs introduced in the Riff-Diff pipeline contain an additional helix that is used as a placeholder for a substrate pocket during structure generation with RFdiffusion. This helix can be placed manually or in an automated manner into the artificial motif library. With manual placement, the user places the helix directly into the initial catalytic array, which can then be identified by the scripts that generate the artificial motif library. The automatic placement mode calculates a vector *V* between the centre of mass of (a) the atoms of the artificial motif and (b) the substrate atoms. The first alpha carbon of the helix is then placed on the centre of mass of the substrate atoms and oriented along vector *V* to point away from the motif. In general, any arbitrary backbone can be used as a placeholder. The automatic placement mode uses a straight alpha-helical fragment of 21 residues with a sequence starting with small side chains (Gly) that increase in size to mimic a cone shape. The sole function of the placeholder helix is to provide a negative shape into which RFdiffusion cannot scaffold a backbone.

Without auxiliary potentials, RFdiffusion generated extended scaffolds that pointed away from the placeholder helix. To ensure that the scaffolds generated by RFdiffusion were both globular and cantered on the artificial motif and its channel placeholder, we implemented a custom auxiliary potential. The auxiliary potential implements a radius of gyration loss to a virtual centre of mass. This virtual centre of mass is calculated using a recentring vector scaled by a customizable *distance* factor. Similar to the placement of the placeholder helix, the user can either specify the direction of the recentring vector ***v*** using xyz coordinates or use this potential in an automated mode. The automated mode calculates the vector *v^* as the vector from the substrate atom centroid to the channel helix centroid. Equation (3) describes this potential, where *w* corresponds to the weight of the auxiliary potential, *d* to the distance factor of the recentring, *v^* to the normalized vector, *r_i_* to the coordinates of every C_ɑ_ atom, and *N* to the number of C_ɑ_ atoms in the structure.

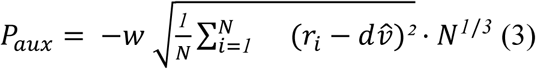

### Addition of carboxylic acid and a tryptophan side chain to MBH catalytic arrays

We used geometry optimization with XTB^53^, version 6.6.1, with the generic force field GFN-FF using the ALPB implicit water solvent model to calculate the interaction between imidazole (histidine side chain) and acetic acid (aspartic and glutamic acid side chains). The orientation of the tryptophan side chain relative to 4-nitrobenzaldehyde was modelled in the same way. Subsequently, we used CREST version 2.12^54^ with the GFN-2 method and ALPB implicit water solvent model to identify the lowest-energy conformer of the tryptophan side chain. The carboxylic acid and tryptophan orientations were combined with the original active site models to create the input for the Riff-Diff pipeline.

### Design of MBH constructs

We used an updated version of the Riff-Diff pipeline that employs LigandMPNN instead of MPNN-informed Rosetta FastDesign during the refinement steps. Rosetta FastRelax was used after ESMFold and AlphaFold2 predictions for evaluation based on ligand movement. For the final refinement step after AlphaFold2 prediction, LigandMPNN was used as well instead of the Rosetta coupled moves protocol in the original version.

For 36 of the designs based on the active site of BH1.8, the helical fragment used for attachment of the active site residue was not predetermined. Instead, the complete rotamer library for each active site residue was ranked according to the probability of each individual rotamer as well as the phi/psi angle occurrence. The resulting rotamer library was further filtered to include only rotamers found in helical regions according to the phi/psi angle preference of this secondary structure. The top 15 rotamers were extracted (Supplementary Figure S17) and a library consisting of all structures in the PDB (filtered for maximum 20% sequence identity) was searched for 7-residue fragments featuring these rotamers and the corresponding phi/psi angle combinations. The fragments were filtered for 100% helical content according to DSSP^66^. Subsequently, all fragments featuring identical rotamers and phi/psi angles over all residues in the fragment were grouped. The resulting fragment library was ranked according to the number of fragments in each group, the active site residue rotamer probability and the active site residue phi/psi angle occurrence. The selected fragments were treated in the same way as the predetermined helical fragment. This alternative approach to fragment generation is also included in the Riff-Diff pipeline. After refinement and evaluation with AlphaFold2, 7 of these constructs were further refined with the original coupled moves protocol. All other 38 designs were refined with LigandMPNN. A complete list of constructs, the design method and substrate conversion can be found in Supplementary Table S7. To select sequences based on the active site of BH1.8, we included the glutamic acid pKa calculated with Propka^67^ and the ligand heavy atom RMSD between the design model and AlphaFold3^68^ predictions with the product in our evaluation metrics.

### Synthesis

All reagents used for synthesis were obtained from Sigma-Aldrich (except for pyridine, obtained from ThermoFisher Scientific) with min. 98% purity and used without further purification. Solvents were of HPLC grade. NMR spectra were measured on a Bruker Avance III 300 MHz NMR spectrometer. Chemical shifts are reported in ppm relative to TMS (δ = 0.00 ppm) and the coupling constants (*J*) in Hertz (Hz).

### *rac-*Methodol (*rac*-1) synthesis

6-Methoxy-2-naphthaldehyde **2** (500 mg, 2.69 mmol, 1 eq) was added to a 1:4 mixture of acetone (5.4 mL) and aqueous phosphate solution (22 mL, 10 mM NaH2PO4, 111 mM NaCl, 2.7 mM KCl in water, p 7.4) (1:4). L-Proline (62.2 mg, 0.2 eq.) was added to the solution and the reaction was stirred at room temperature for 48 h. Since the reaction had not sufficiently proceeded based on TLC monitoring, more acetone (5.4 mL) and L-Proline (62.2 mg) were added after 48 h, and the reaction was kept stirring for another 48 h, after which conversion appeared complete on TLC. Purification by flash chromatography (cyclohexane/ethyl acetate, 2:1) and evaporation of the solvent under reduced pressure yielded the final racemic product methodol (4-hydroxy-4-(6-methoxy-2-naphthalenyl)-2-butanone **1**) as a white solid (400 mg, 1.64 mmol) in 61% yield. The spectral data were in accordance with the literature^69^ (Supplementary Figures S18-S19). ^1^H NMR (300 MHz, CDCl3) δ 7.76 – 7.68 (m, 3H), 7.42 (dd, J = 8.5, 1.9 Hz, 1H), 7.19 – 7.08 (m, 2H), 5.29 (dd, J = 8.7, 3.7 Hz, 1H), 3.92 (s, 3H), 3.28 (br s, 1H), 3.03 – 2.80 (m, 2H), 2.21 (s, 3H). ^13^C NMR (75 MHz, CDCl3) δ 209.28 (s), 157.87 (s), 137.94 (s), 134.24 (s), 129.58 (s), 128.86 (s), 127.33 (s), 124.42 (s), 124.41 (s), 119.18 (s), 105.78 (s), 70.11 (s), 55.44 (s), 52.07 (s), 30.96 (s).

### Synthesis of MBH product 2-(hydroxy(4-nitrophenyl)methyl)cyclohex-2-en-1-one

The product standard for the MBH reaction was synthesized as described previously^70^. The product was purified by flash column chromatography (9:1 pentane/ethyl acetate) yielding white crystals (190 mg, 7.7% yield). The spectral data were in accordance with the literature^71^ (Supplementary Figures S20 and S21). ^1^H NMR (300 MHz, CDCl3) δ 8.18 (d, J = 8.8 Hz, 2H), 7.54 (d, J = 8.2 Hz, 2H), 6.82 (t, J = 4.2 Hz, 1H), 5.60 (d, J = 6.0 Hz, 1H), 3.57 (d, J = 6.0 Hz, 1H), 2.49 – 2.39 (m, 4H), 2.00 (p, J = 6.2 Hz, 2H). ^13^C NMR (75 MHz, CDCl3) δ 200.26, 149.42, 148.33, 147.35, 140.30, 127.26, 123.66, 72.17, 38.55, 25.91, 22.50.

### HPLC analysis of methodol absolute configuration and enantiomeric excess

HPLC analyses were performed on a Shimadzu system (DGU-20A On-line Degasser, LC-20AD pump, SIL-20AC autosampler, CBM-20A system controller, SPD-M20A Photodiode Array Detector, Shimadzu CTO-20AC Column Oven). The samples (5 µL) were analysed with an isocratic flow according to the following methods:

For elution order and absolute configuration: analysis on a Daicel Chiralpak IB (250 mm, ID 4.6 mm, particle size 5 µm) using *n*-heptane/2-propanol 90:10 (isocratic, flow rate of 1 mL/min, 30 °C, wavelength 254 nm); Retention times: (*S*)-methodol 12.6 min, (*R*)-methodol 13.2 min^72^, see Supplementary Figure S22a and c.

For determination of enantiomeric excess: Since baseline separation was obtained more easily on an OD-H column, analysis was performed on a Daicel Chiralcel OD-H (250 mm, ID 4.6 mm, particle size 5 µm) using *n*-heptane/2-propanol 92:8 (isocratic, flow rate of 1 mL/min, 30 °C, wavelength 254 nm). Retention times: (*S*)-methodol 18.5 min, (*R*)-methodol 20.0 min; see Supplementary Figure S22b and d.

## Acknowledgments

We thank A. Winkler and P. Macheroux for discussions about mass spectrometry and enzyme kinetics and Anthony Green for providing the coordinates of a calculated transition state 3 of MBH. Additionally, we acknowledge the European Synchrotron Radiation Facility for provision of synchrotron radiation facilities, and we would like to thank the staff of the ESRF and EMBL Grenoble for assistance and support in using beamlines MASSIF-3, ID30B, and BM29. For open access, the authors have applied a CC BY public copyright license to any Author Accepted Manuscript version arising from this submission.

## Author contributions

Conceptualization: M.B., A.T., G.O.; Methodology: M.B., A.T., G.O.; Software: M.B., A.T.; Validation: M.B., A.T., G.O., W.E, C.F..; Formal analysis: M.B., A.T., S.K., W.E.,; Investigation: M.B., A.T., M.C., S.K., M.T., D.S., A.B., W.E., S.Y.H., M.A., C.F., M.M., H.L., F.R., M.H., G.O.; Resources: M.H., G.O.; Data Curation: M.B., A.T., A.B., G.O.; Writing: M.B., A.T., G.O.; Visualization: M.B., A.T., S.K.; Supervision: M.H., G.O.; Project administration: G.O.; Funding acquisition: G.O.

## Materials & Correspondence

Supplementary Materials are available for this paper. Custom Python scripts, RosettaScripts, example input command lines can be found on github (https://github.com/mabr3112/riff_diff_protflow), and corresponding data has been deposited to Zenodo (https://doi.org/10.5281/zenodo.15494858). Correspondence and requests for materials should be addressed to GO.

