## Supplementary Materials for "Computational enzyme design by catalytic motif scaffolding"

**The PDF file includes:**

Supplementary Methods
Figs. S1 to S22
Tables S1 to S10

### Supplementary Methods

#### The Riff-Diff pipeline

The full codebase to the latest version of RiffDiff including detailed instructions for use can be found at https://github.com/mabr3112/riff_diff_protflow. This repository will be kept up to date and also includes all Rosetta .xml scripts that are used during the refinement steps of the protocol. The original implementation of RiffDiff, used in the generation of the retro-aldol enzymes, can be found at https://github.com/mabr3112/riff_diff_original. The following section is a detailed description of the design steps performed by the scripts of the RiffDiff pipeline.

##### Motif library generation

Motif library construction can proceed via two modes: (1) using a predefined backbone structure (“backbone rotamer finder”), or (2) extracting fragments from a filtered PDB database (“fragment finder”).

###### *Backbone rotamer finder*

By default, a 7-residue helical fragment (with phi/psi angles of -64.8/-41.0 at each position) is used as input. For each residue in the catalytic array, the rotamer library of the corresponding amino acid is retrieved. For each position on the fragment that is not on the N- or C-terminus, the phi and psi angles are determined (default -64.8/-41.0) and the corresponding rotamers are extracted from the Dunbrack[1] rotamer library as provided with Rosetta. Rotamers with $<5\%$ overall probability and $<30\%$ of the probability of the highest likelihood rotamer at a given fragment position are filtered out. To expand conformational diversity, additional rotamers are generated by systematically perturbing each chi angle by $\pm1$ and $\pm2$ standard deviations from the mean values provided in the rotamer library. The rotamers are ranked according to a composite score of the log rotamer probability and the log phi/psi angle occurrence for the corresponding amino acid identity. The selected rotamers are attached to each specified fragment position. A maximum number of 15 fragments (corresponding to 15 unique rotamers) per input position on the fragment is returned.

###### *Fragment finder*

For each residue in the catalytic array, the rotamer library of the corresponding amino acid identity is retrieved. The rotamers are ranked according to a composite score of the log rotamer probability and the log phi/psi angle occurrence for the corresponding amino acid identity. If a fragment secondary structure was provided (e.g. helical), the rotamer library is filtered for all phi/psi angles that correspond to the selected secondary structure. Only rotamers meeting all of the following criteria are retained: (i) the backbone phi/psi combination must occur in at least 0.5% compared to all phi/psi combinations for the given amino acid, (ii) the rotamer itself must have a probability of at least 5%, and (iii) its probability must be at least 30% of that of the most probable rotamer. The rotamers are ranked according to a composite score of the log rotamer probability and the log phi/psi angle occurrence for the corresponding amino acid identity. A maximum number of 15 rotamers per fragment position is returned.

From a version of the PDB containing structures with a maximum sequence identity of 20%, all positions with the same amino acid identity, identical phi, psi (both rounded to the nearest ten) and chi angles (± two standard deviations) as well as the three adjacent residues in both directions are extracted, creating fragments of 7-residue length. The extracted fragments are filtered for secondary structure content (e.g. 100% helical). All extracted fragments are grouped by the phi/psi/omega angles (rounded to the nearest ten) of all residues in the fragment and assigned a score that corresponds to the number of fragments in each group. This score (referred to as "backbone probability") indicates how common this fragment is in the PDB. For all fragments within a group, the median of phi/psi/omega is calculated, and an updated fragment is generated using these values. The resulting fragments are ranked according to their log backbone probability, log rotamer probability and log phi/psi occurrence.

***Motif library***

The selected fragments are superimposed with the input catalytic array via their functional groups. A clash detection is performed, removing all fragments that show clashes with the ligand. Additionally, fragments for each residue in the catalytic array are filtered for a minimal backbone RMSD (0.3 Å) to all other selected fragments. Next, clash detection between the fragments of each residue in the catalytic array is performed. All clashing fragment combinations are removed. The resulting motifs are ranked according to their average backbone probability (for fragments that are derived from Fragment Finder protocol), rotamer probability and phi/psi occurrence. The placeholder helix is added in the last step to create the final catalytic motifs.

#### Structure generation

This section describes the RiffDiff pipeline as used for the generation of MBH constructs with default settings. The original version that was used to design the RAD constructs is largely similar but used a ProteinMPNN-informed Rosetta FastDesign for sequence generation instead of LigandMPNN. The updated version also includes more stringent filtering steps to reduce computational costs. The Riff-Diff structure generation script is divided into 4 phases: Screening, Refinement, Evaluation, and Variant Generation. We refer to design models as "poses" throughout the design trajectory. Unless noted otherwise, RMSDs are calculated relative to orientations in the input catalytic motif and TM scores are calculated between poses and predicted structures. In all LigandMPNN runs, the specified catalytic residues are kept fixed, all other positions are set to designable. After each prediction step (ESMFold or AlphaFold2), the ligand is copied back into the pose via a superimposition of the input motif residues.

***Screening***

The top 200 catalytic motifs are selected as input for structure generation. Per input motif, 5 RFdiffusion trajectories are performed, using 30-residue stretches on the N- and C-terminus and 10-52 residue stretches in between fragments for four 7-residue fragments (example contig: [Q1-21/0 30/A1-7/10-52/B1-7/10-52/C1-7/10-52/D1-7/30], where Q indicates the 21-residue placeholder helix, and A, B, C and D indicate 7-residue fragments). The total length of diffused backbones is exactly 200 residues.

After RFdiffusion, the placeholder helix is removed and the ligand is copied back in, as described above. The poses are filtered based on radius of gyration (< 18 Å, ROG), number of CA-atoms per ligand atom within 8 Å (> 5, referred to as "ligand contacts"), backbone RMSD of the catalytic residues (< 1 Å, referred to as "catres bb RMSD") and backbone RMSD of all residues in the input motif (< 1 Å, referred to as "motif RMSD").

LigandMPNN is used to generate initial sequences on all poses that passed the initial filtering (3 per input structure). The returned sequences are threaded onto the input structures and relaxed once with Rosetta, while constraining the catalytic residue side chains as well as the motif backbone atoms to the input motif coordinates. This step will be referred to as "constrained relax" in subsequent paragraphs. Next, sequences are generated based on the relaxed output using LigandMPNN (30 per structure). The resulting sequences are filtered based on a composite score composed of the Rosetta constrained relax total score, ligand contacts after constrained relax, the number of ligand clashes with backbone atoms after constrained relax (referred to as "ligand clashes", ROG, and the overall confidence score of LigandMPNN. The top 5000 poses are predicted with ESMFold. Output poses are filtered based on ROG (< 18 Å), motif RMSD (< 1.5 Å), catres bb RMSD (< 1.5 Å), TM score to the design model (> 0.9), average per-residue ESMFold pLDDT (> 70), and ligand contacts (> 5). All passing poses are ranked according to a composite score composed of average per-residue ESMFold pLDDT, TM score, catres bb RMSD, motif RMSD, ligand clashes, ligand contacts, and ROG. The top scoring pose according to these metrics for each RFdiffusion output is kept for the next phase.

***Refinement***

For each refinement cycle, poses are constrained-relaxed and 25 sequences are generated using LigandMPNN. Next, the design sequences are predicted using ESMFold and filtered based on catres bb RMSD, motif RMSD, average ESMFold pLDDT, ligand contacts (> 5) and TM score (> 0.9). Filter cutoffs for catres bb RMSD, motif RMSD and average pLDDT are ramped with increasing number of refinement cycles, starting at 1.2 Å/ 1.5 Å/ 75 and ending at 0.7 Å / 1.0 Å / 85. All passing poses are ranked according to a composite score (referred to as "ref comp score") composed of these scoreterms and RMSD of catalytic residue side chains (referred to as "side chain RMSD"), and ligand clashes. The top three poses per unique RFdiffusion output are passed to the next refinement cycle. After five cycles of refinement, the top 500 poses according to ref comp score are passed on for evaluation.

##### *Evaluation*

For each designed sequence, five predictions using AlphaFold2 in single sequence mode are performed. The top pose according to average pLDDT is selected, and poses are filtered according to average pLDDT (> 85), TM score (> 0.9), catres bb RMSD (< 0.7 Å), and ligand contacts (> 5). Poses (without ligand) are repacked using AttnPacker and side chain RMSD is calculated. All passing poses are relaxed with the ligand present (without constraints) 15 times, and the mean sidechain RMSD, ligand RMSD, and catres bb RMSD are calculated. The poses are ranked according to the evaluation composite score (all above mentioned values and the number of ligand clashes and surface aggregation propensity (SAP)).

##### *Variant generation*

In the final step of the Riff-Diff pipeline, custom mutations can be introduced upon inspection of the evaluation poses. This is useful e.g. to open channels that are blocked by residue side chains.

Poses are relaxed with constraints. If mutations were specified, the amino acid identities at the specified positions are restrained during the following LigandMPNN step (50 sequences per pose). The resulting sequences are predicted using ESMFold and filtered in the same way as the final refinement step. All passing sequences are predicted again with AlphaFold2 and filtered and ranked in the same way as during the evaluation step.

Instead of LigandMPNN-based variant generation, an alternative Rosetta coupled-moves based protocol can be employed. Following constrained relax, this protocol is performed 50 times for each pose. Designable positions are selected based on the DetectProteinLigandInterface TaskOperation with cut parameters of 6, 8, 10 and 12 Å. For each designable position, the occurrence of each amino acid in the coupled moves output sequences is evaluated. Amino acid identities chosen in 25% of all trajectories (or all above 10%, if none passes this cutoff) are selected, and every possible combination of residue identities is generated. The resulting sequences are then predicted with ESMFold and AlphaFold2, as described above.

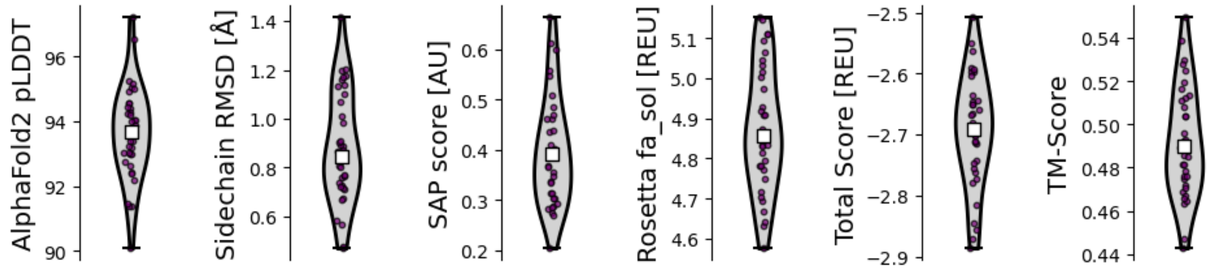

**Fig. S1.**

**Computational metrics of selected RAD models.** All 36 sequences were assigned high confidence by AlphaFold2. The displayed AlphaFold2 pLDDT corresponds to the average per-residue pLDDT per structure. Average sidechain RMSD to the catalytic geometry was below one angstrom. SAP and Rosetta scores were calculated per residue. The displayed TM scores represent the TM score calculated by TMAlign to the closest match in the PDB by Foldseek. White squares correspond to the median of the distributions, purple dots to individual designs.

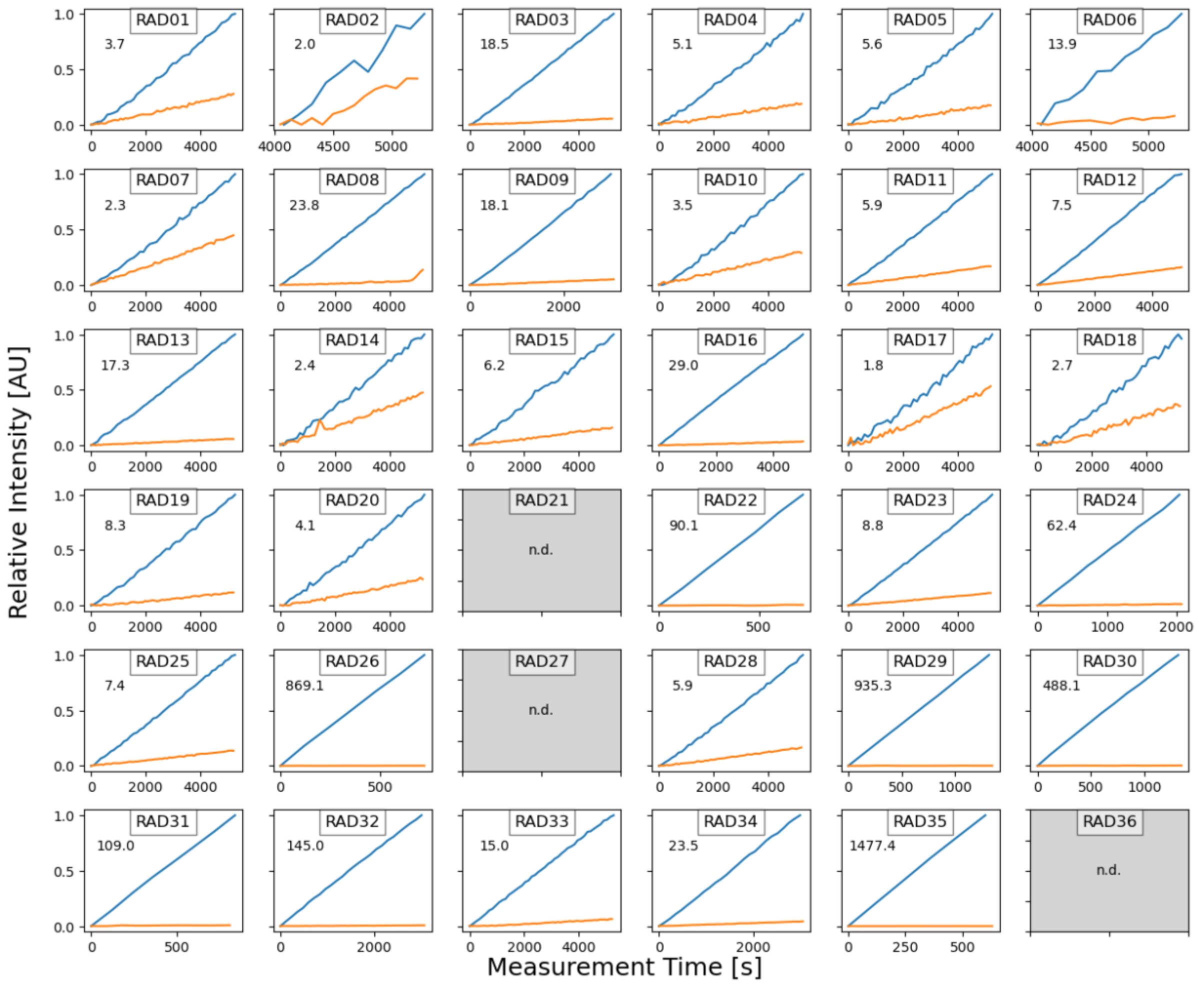

**Fig. S2.**

**Mutation of active site lysine residues to alanine leads to significant loss of activity of designed retro-aldolases.** Relative reaction progress for original constructs (blue) and active site lysine to alanine variants (orange). Numbers denote n-fold decrease in activity. Variants of highly active constructs RAD29 and RAD35 display 935.3 and 1477.4-fold decrease, respectively.

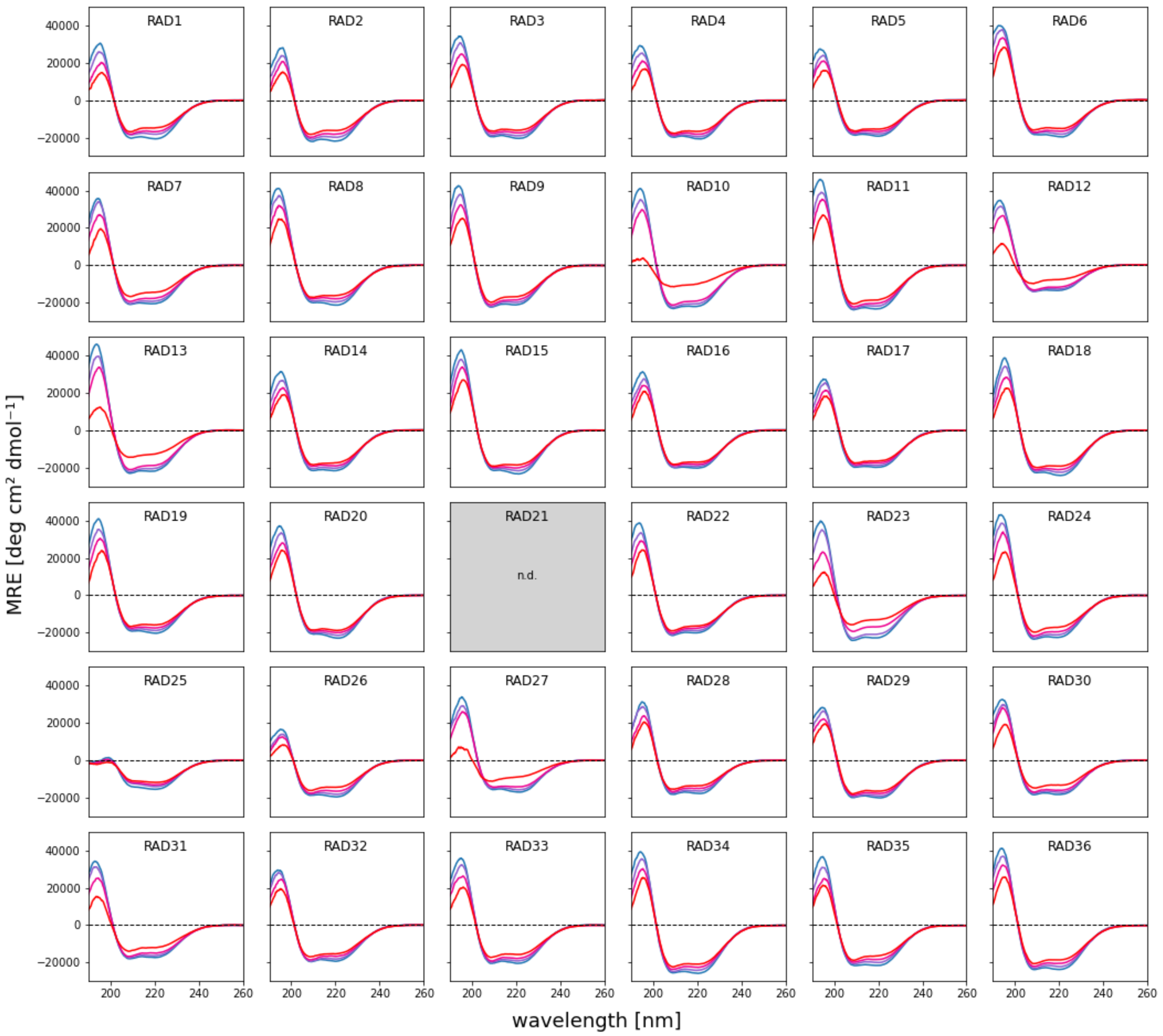

**Fig. S3.**

Circular dichroism spectra of designed RAD enzymes at 20 °C (blue), 45 °C (purple), 70 °C (pink) and 95 °C (red) confirm alpha-helical fold and high thermodynamic stability. For 30 constructs, heating the samples to 95 °C yielded a decrease in CD signal intensity at 220 nm between 75% to 85% of the initial value, indicating only marginal unfolding (Figure 2B). RAD10, RAD12, RAD13, RAD27 and RAD31 show the onset of unfolding at 95°. The thermal denaturation curve for RAD23 shows a higher degree of unfolding compared to all other constructs starting at 60 °C but no clearly defined melting temperature can be assigned, indicating uncooperative unfolding. Except for RAD10 and RAD13, the CD spectra after heating to 95 °C and cooling to 20 °C closely resemble the spectra before the thermal scan was performed.

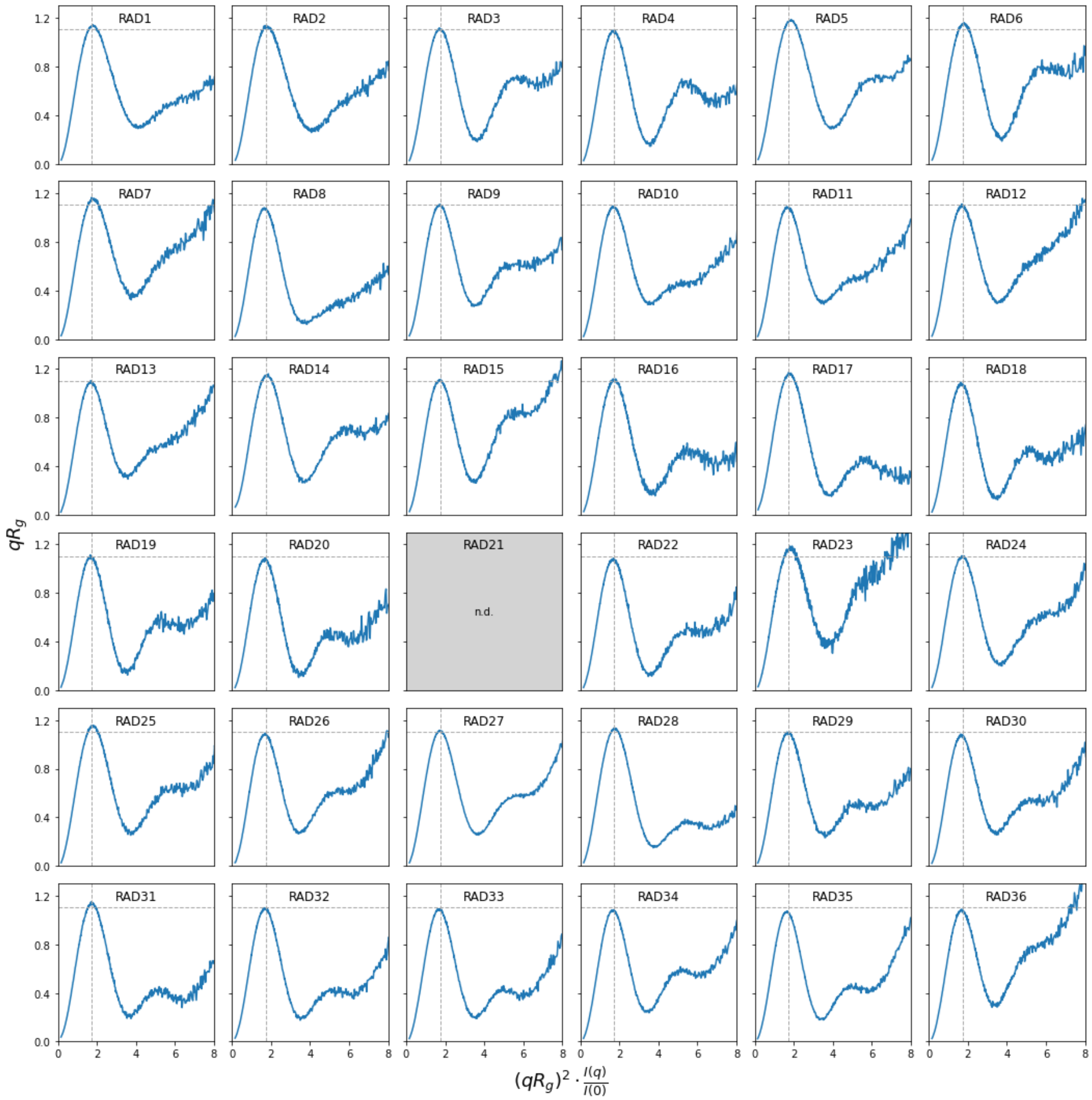

**Fig. S4.**

Dimensionless Kratky plots derived from experimental SAXS profiles confirm globular shape and high degree of foldedness of the designed enzymes, indicated by the peak maxima close to Rg = $3^{1/2}$ and $(qR_g)^2 I(q)/I(0) = 1.104$ (dotted lines).

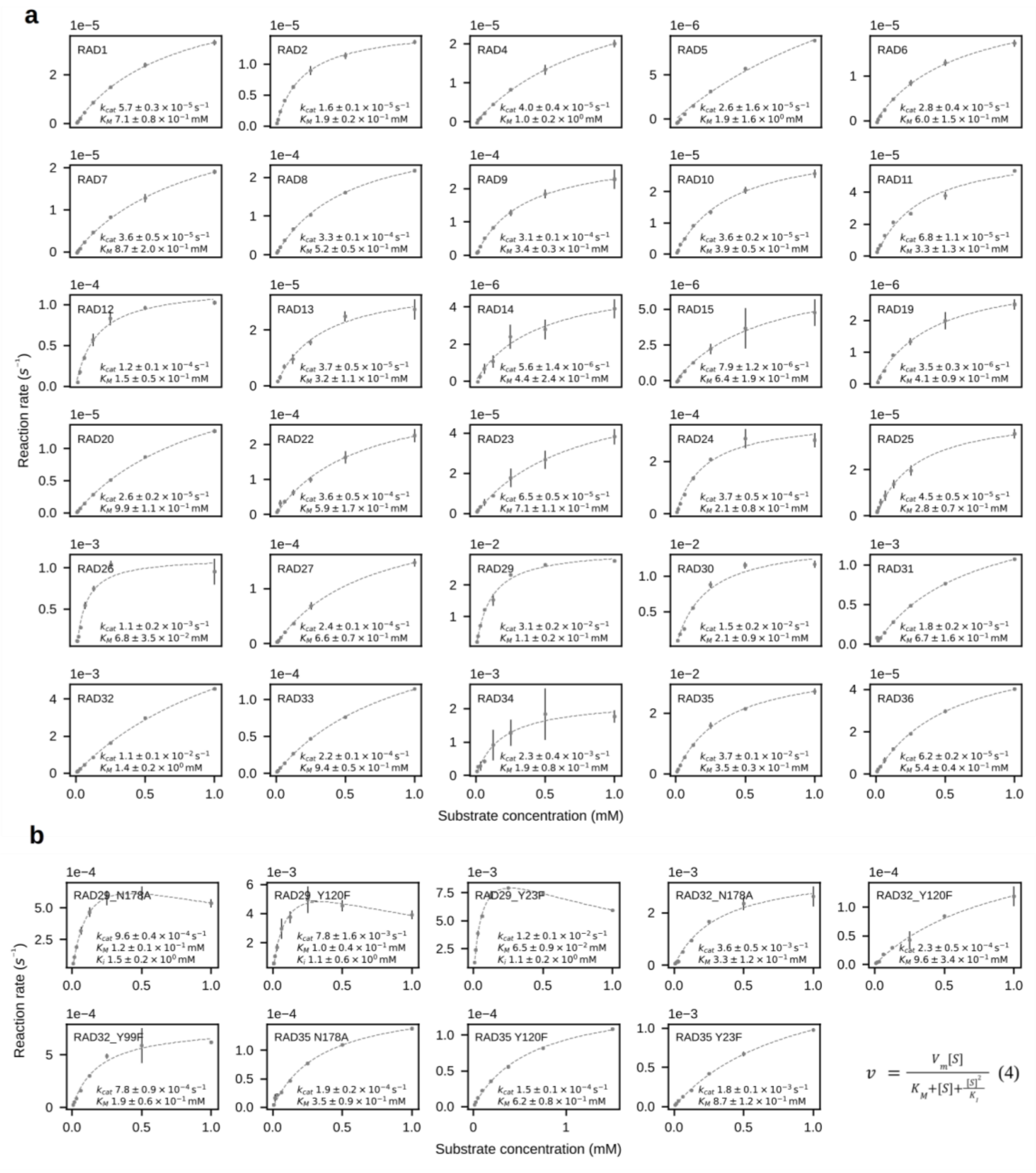

**Fig. S5.**

Michaelis Menten kinetics of (**a**) designed retro-aldolases and (**b**) their site-directed mutagenesis variants with *rac*-methodol as substrate. Reaction rates at individual concentrations were measured in triplicates. Error bars indicate standard deviations. Designs for which Michaelis Menten parameters could not be obtained were omitted. RAD29 variants in panel (b) were fitted using the Haldane substrate inhibition model (equation 4) and the inhibition constants are reported as $K_i$ in the corresponding panels.

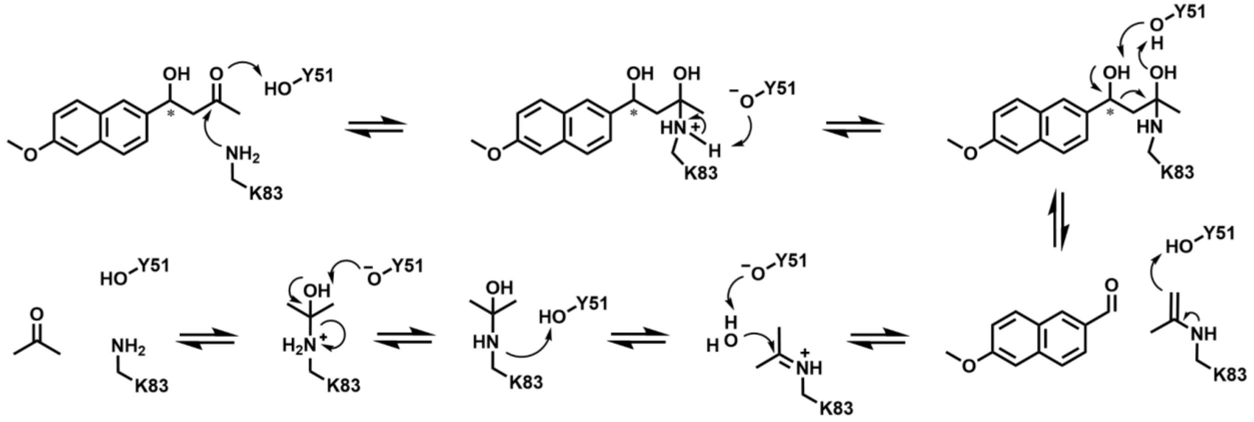

**Fig. S6.**

Proposed reaction mechanism for the RA98.5-8F-catalyzed retro-aldol reaction of methodol.

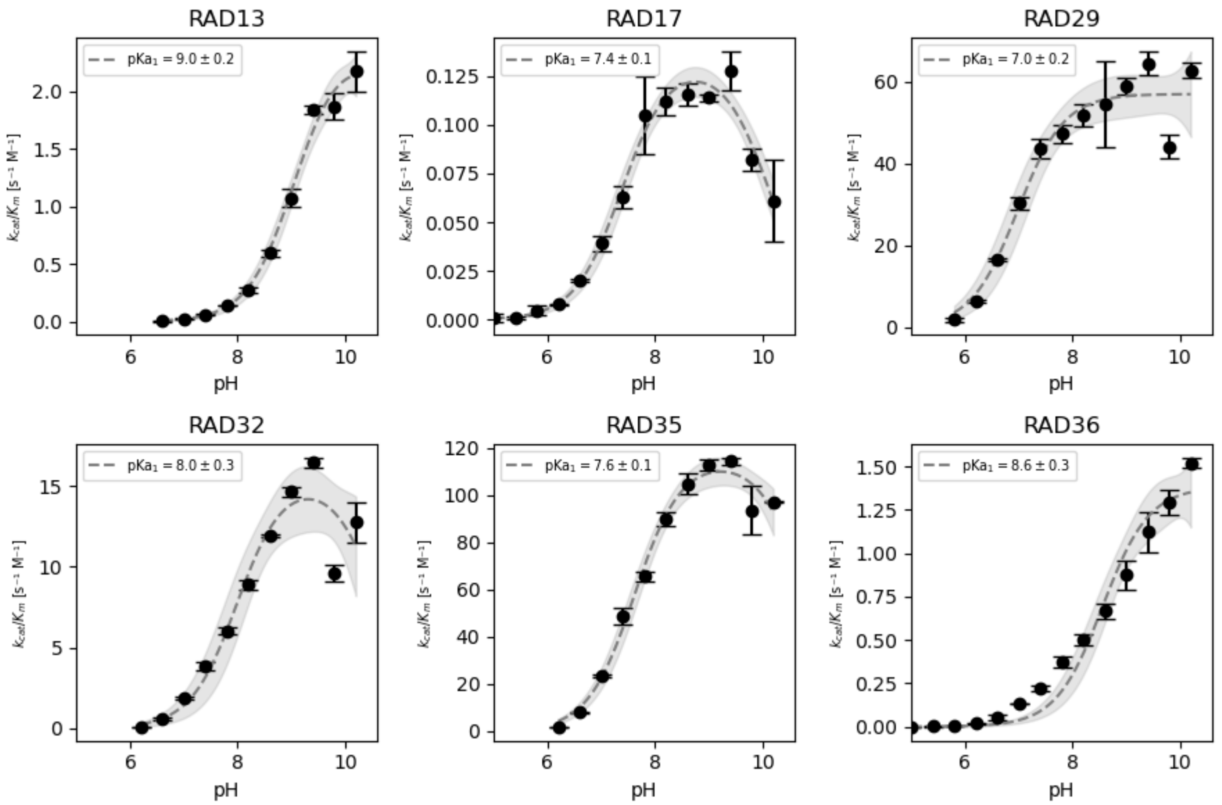

**Figure S7.**

pH profiles of RADs. Gray areas indicate the 95% confidence interval of the fit, error bars indicate standard deviations.

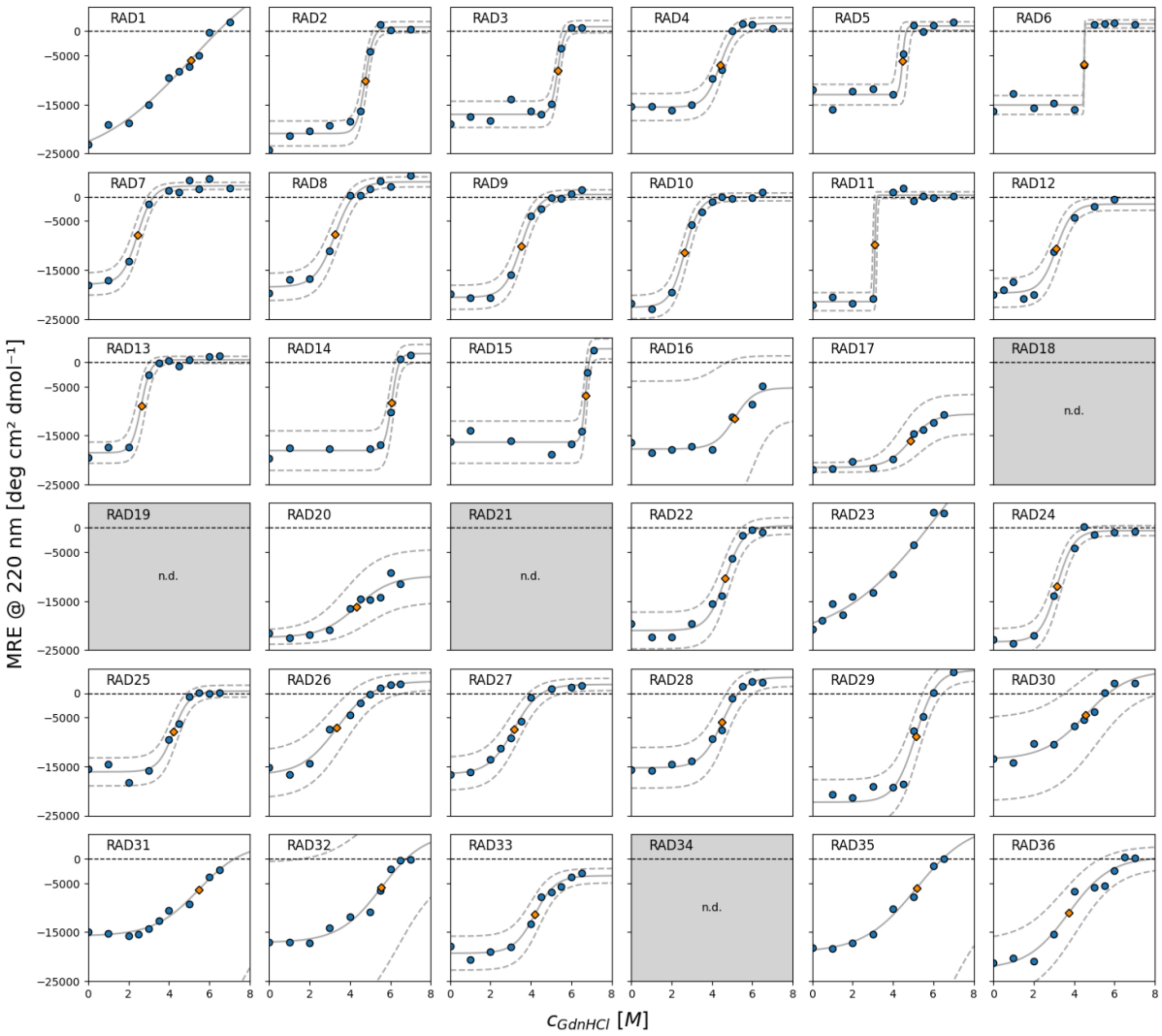

**Fig. S8.**

Chemical denaturation with GdnHCl monitored via CD spectroscopy at 220 nm (MRE = mean residue ellipticity). Orange dots represent the denaturation midpoints calculated from a sigmoidal fit (gray lines). The confidence interval (95%) of the fit is shown in dotted gray lines. Highest resistance to chemical denaturation was observed for RADs 14-17 and RAD20, with midpoints of the denaturation transition above 6 M of guanidinium hydrochloride (GdnHCl). RAD1, RAD23, RAD30, RAD31, RAD32 and RAD36 displayed non-cooperative unfolding, impeding determination of accurate denaturation midpoints. RAD18, RAD19 and RAD34 were not measured. All other constructs display denaturation midpoints between 2.6 and 5.3 M GdnHCl.

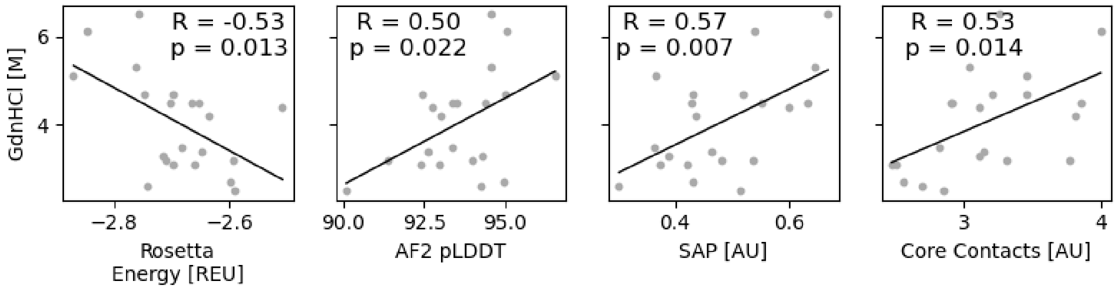

**Fig. S9.**

**Individual correlations of computational metrics used for the denaturation model.** Correlations and statistical significance were calculated using python libraries numpy and SciPy. AF2 pLDDT corresponds to the average per-residue pLDDT per structure of AlphaFold2 predictions. Rosetta score and SAP score were calculated per-residue as metrics to be independent of the protein size. Core contacts were calculated using the RosettaScripts AtomicContactCount filter and represent the atomic density of the protein.

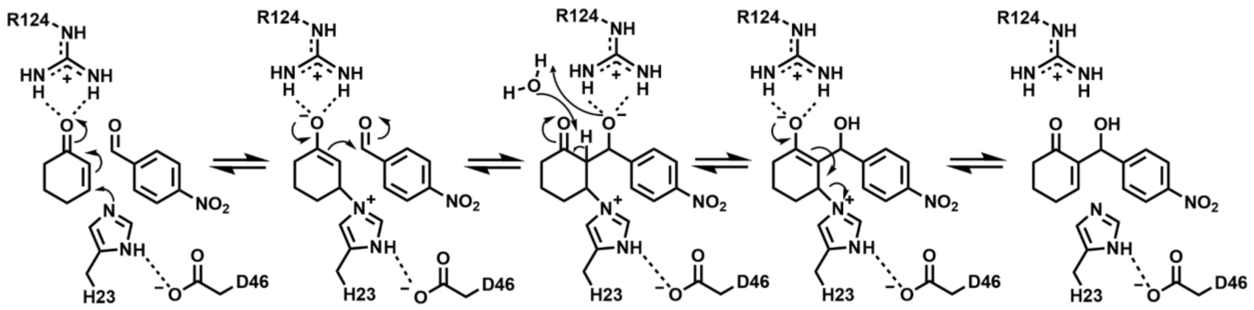

**Fig. S10.**

Proposed reaction mechanism for the BH32.14-catalyzed Morita-Baylis-Hillman reaction of 2-cyclohexenone and 4-nitrobenzaldehyde. In BH1.8, Arg124 is not present, and the water-mediated proton transfer step is facilitated by Glu26.

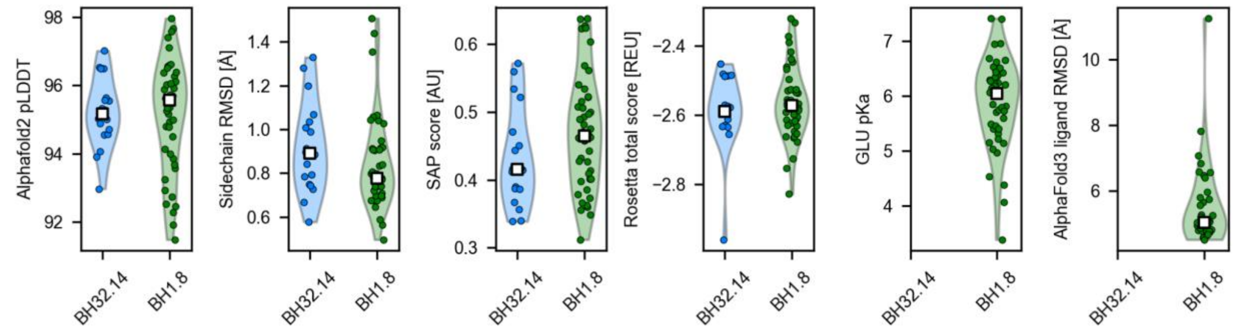

**Figure S11.**

Computational metrics for de novo designed MBHases for each set of active site models. White squares indicate median values; dots indicate values of individual designs. Alphafold2 pLDDT corresponds to the average per-residue pLDDT per structure. Sidechain RMSD was calculated from AttnPacker-repacked[2] AlphaFold2 predictions compared to the geometry of the catalytic array. Surface aggregation propensity (SAP) and Rosetta total energy are divided by the number of residues in each design. For designs based on the active site of BH1.8, the PROPKA-calculated pKa of the glutamic acid residue facilitating the proton transfer and the RMSD of the MBH reaction product compared to the substrate in the catalytic array is provided as well.

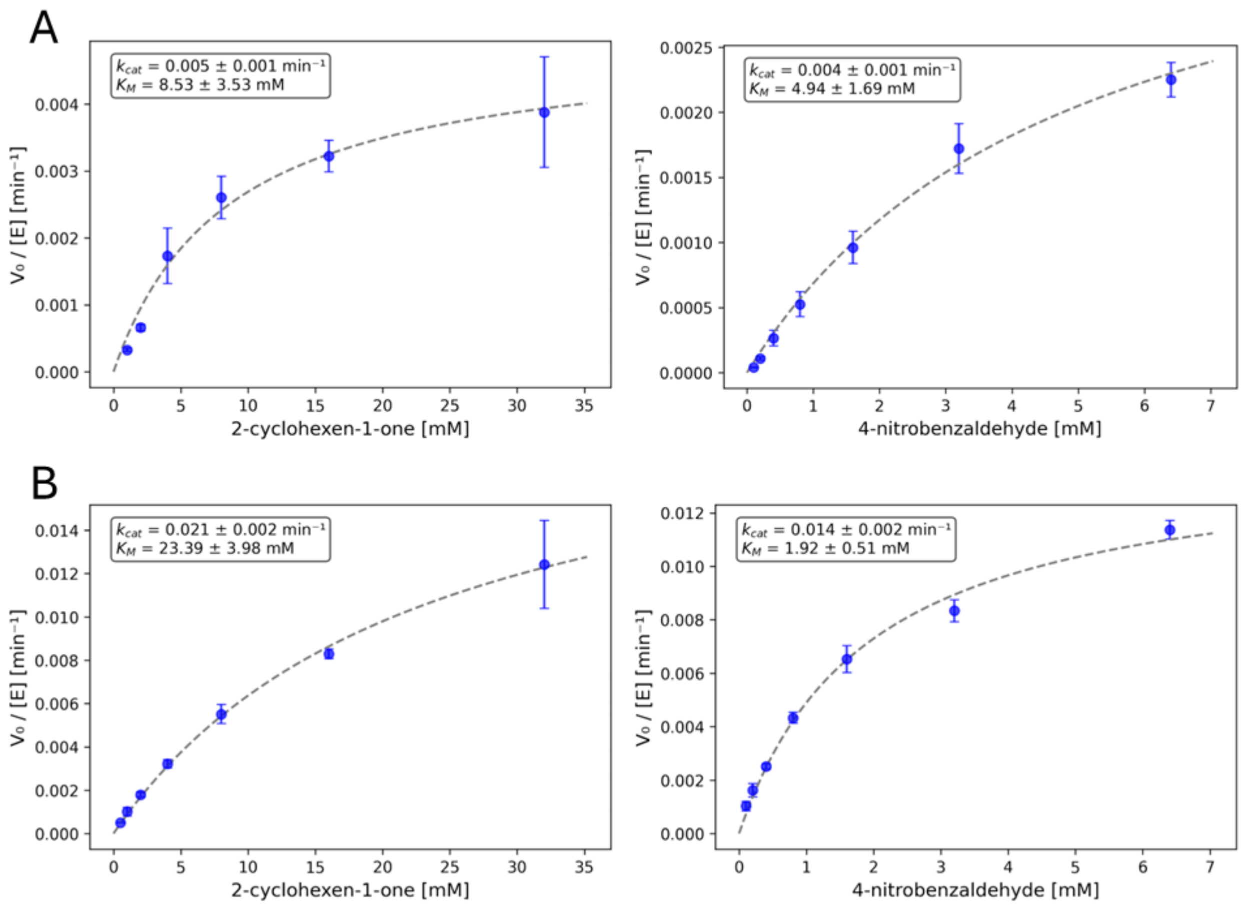

**Fig. S12.**

Fits of initial velocities against 2-cyclohexen-1-one and 4-nitrobenzaldehyde for MBH18 (A) and MBH48 (B). Error bars indicate standard deviations of triplicate measurements.

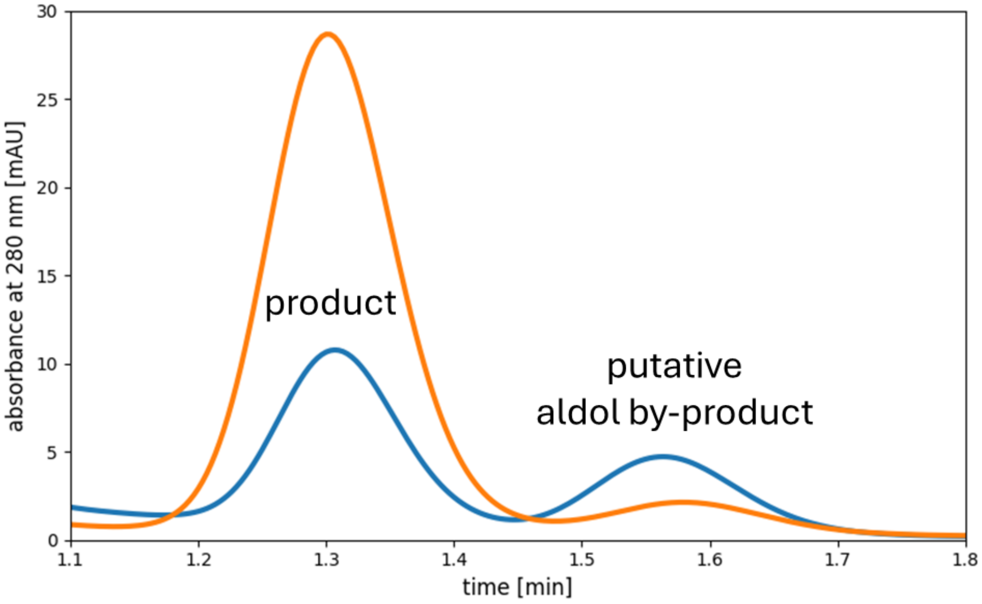

**Fig. S13.**

**HPLC traces for the MBH reaction of 2-cyclohexen-1-one and 4-nitrobenzaldehyde catalysed by MBH18 (blue) and MBH48 (orange).** The peak on the right shows similar retention times as previously reported for an aldol by-product in the MBH reaction[3]. The HPLC trace of MBH18 was shifted by 0.1 minutes to generate this plot, as we observed a reduction in retention times with increasing number of runs on the HPLC column.

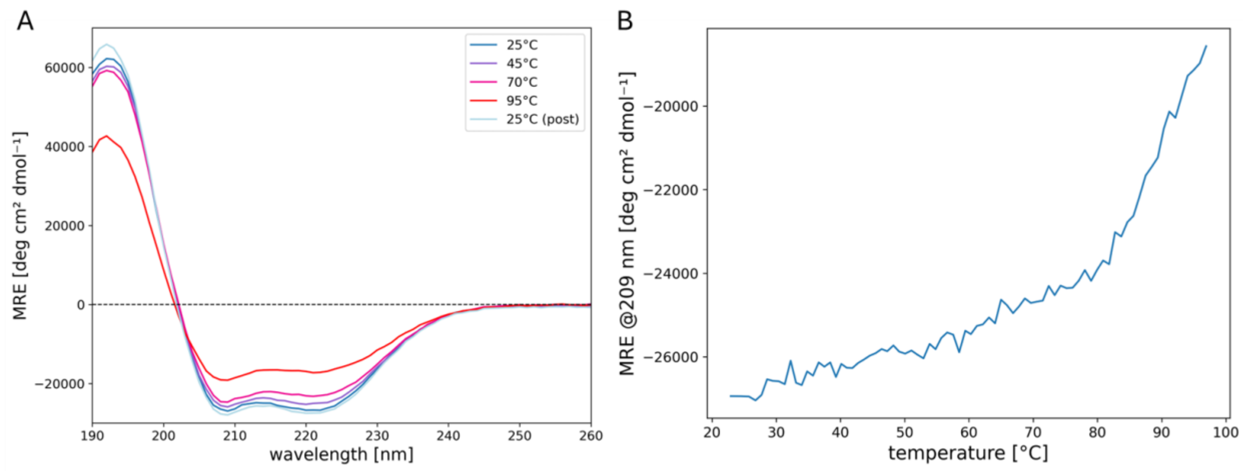

**Fig. S14.**

**Circular dichroism data of MBH48.** A: CD spectra at 25, 45, 70, 95 °C and after cooling back down to 25 °C confirm helical fold. B: CD signal intensity at 209 nm versus temperature indicates onset of unfolding at ~85 °C.

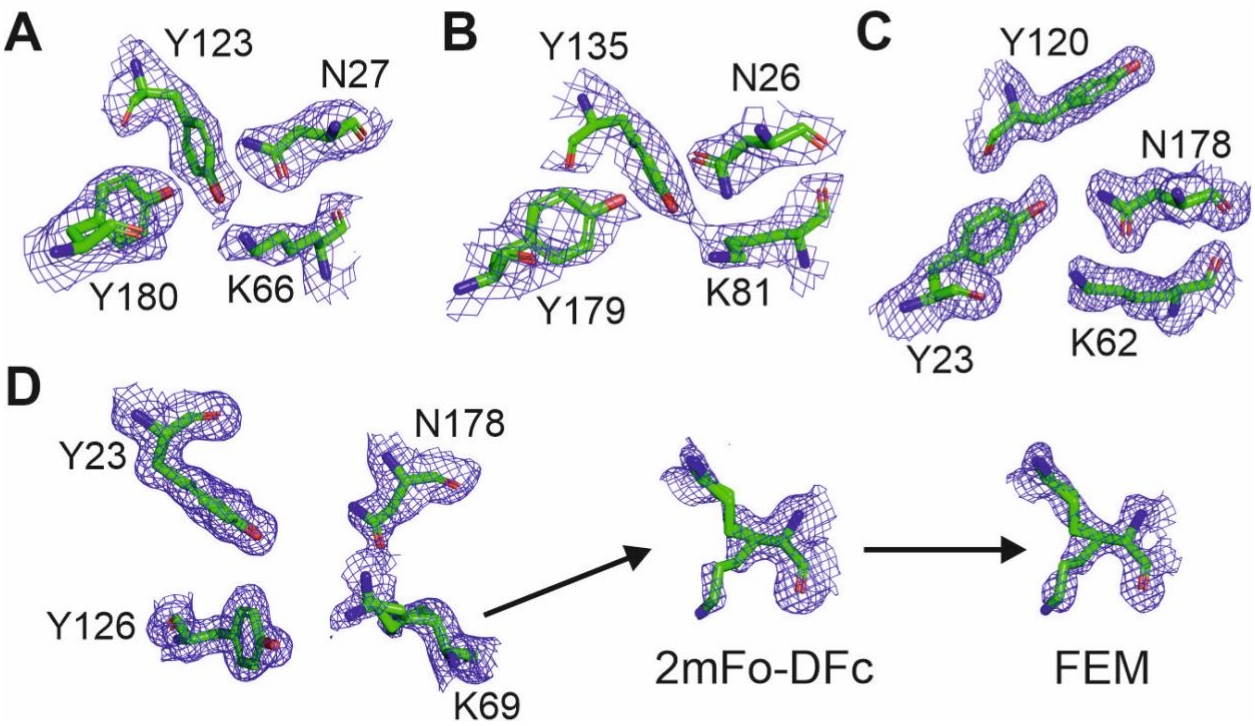

**Fig. S15.**

Electron density for the catalytic residues of RAD13 (A), RAD17 (B), RAD32 (C), and RAD36 (D). The 2mFo-DFc electron density map contoured at 1.0σ is shown for the tetrad residues of each structurally solved RAD. As residue K69 of RAD36 showed weak and diffuse electron density for multiple conformations (2mFo-DFc), a feature-enhanced map (FEM) was created to improve the features of the conformers. Panel (D) highlights the improvement of the electron density for K69 by FEM.

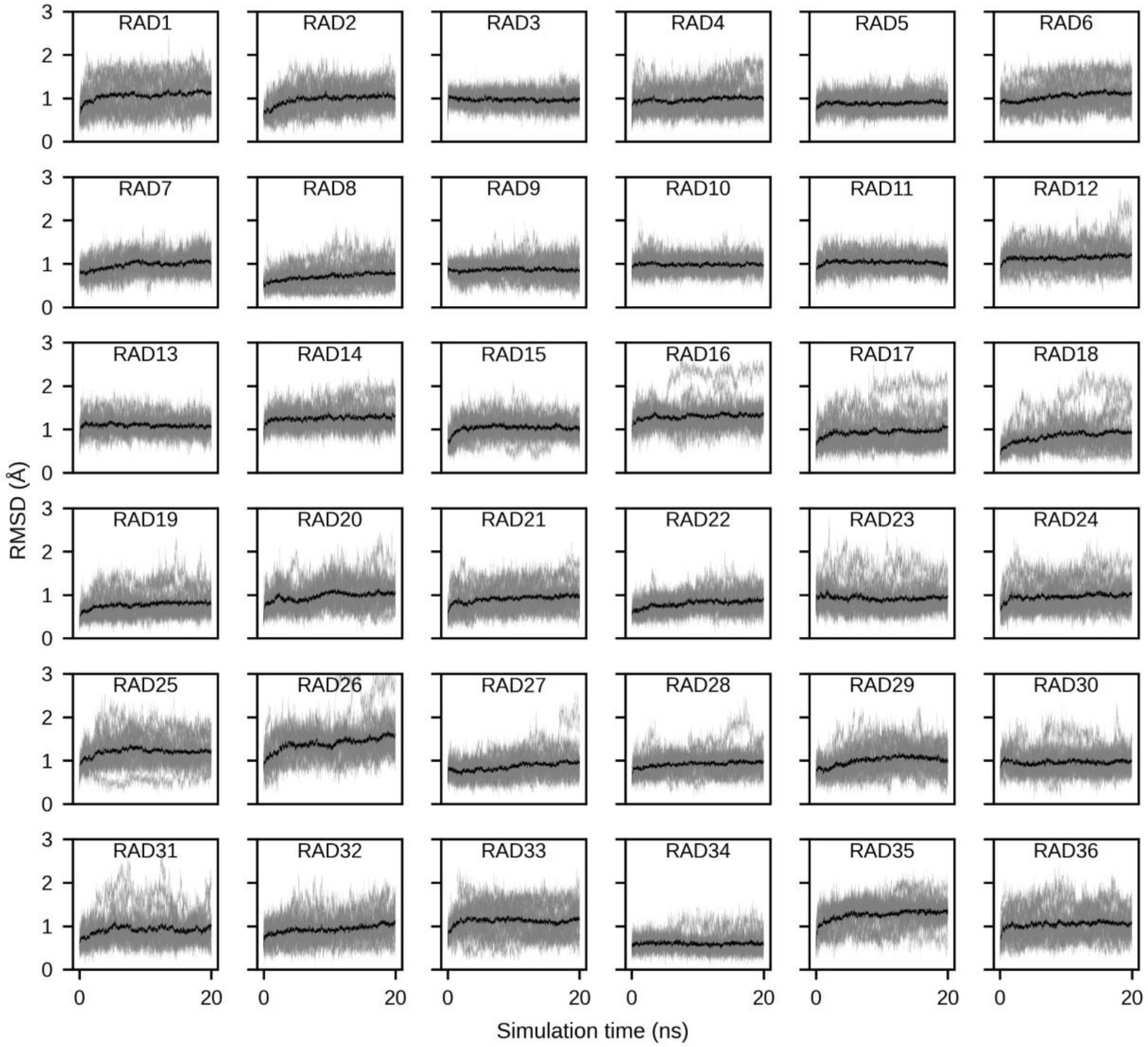

**Fig. S16.**

**Backbone Cα RMSDs of the simulated RADs for all MD trajectories.** Grey lines represent RMSDs measured for each individual trajectory and black lines correspond to the average over all 20 trajectories.

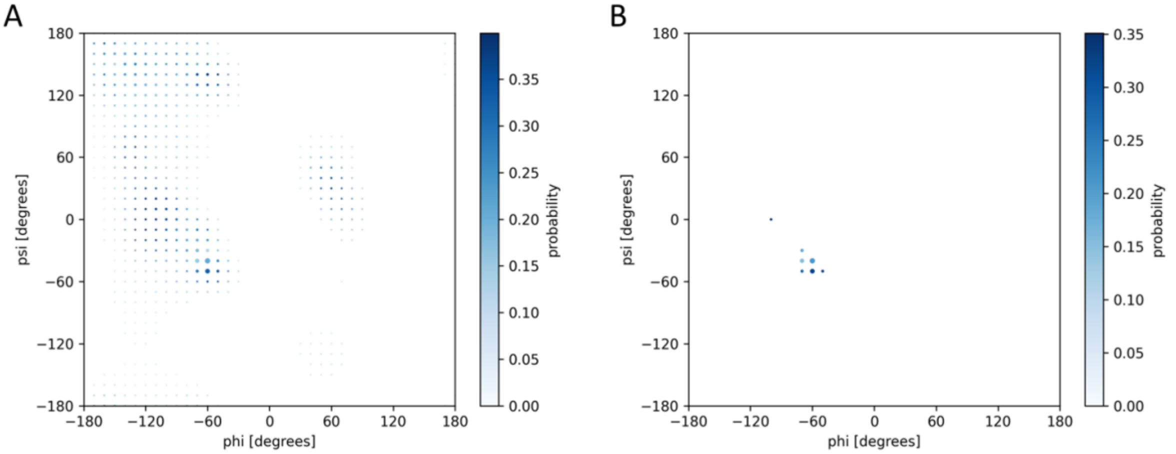

**Figure S17.**

**Selection of rotamers for the nucleophilic histidine residue in the MBH reaction.** A. Ramachandran plot of histidine. The colour gradient indicates the probability of the most common rotamer at each phi/psi angle combination. Dot size corresponds to the occurrence of the phi/psi angle combination for this residue. Only rotamers on phi/psi angles in the helical region of the Ramachandran plot were considered for selection. All data was extracted from a previously published backbone-dependent rotamer library[1]. B. The top 15 rotamers according to probability and phi/psi angle occurrence were selected (multiple rotamers for a single phi/psi angle combination are possible).

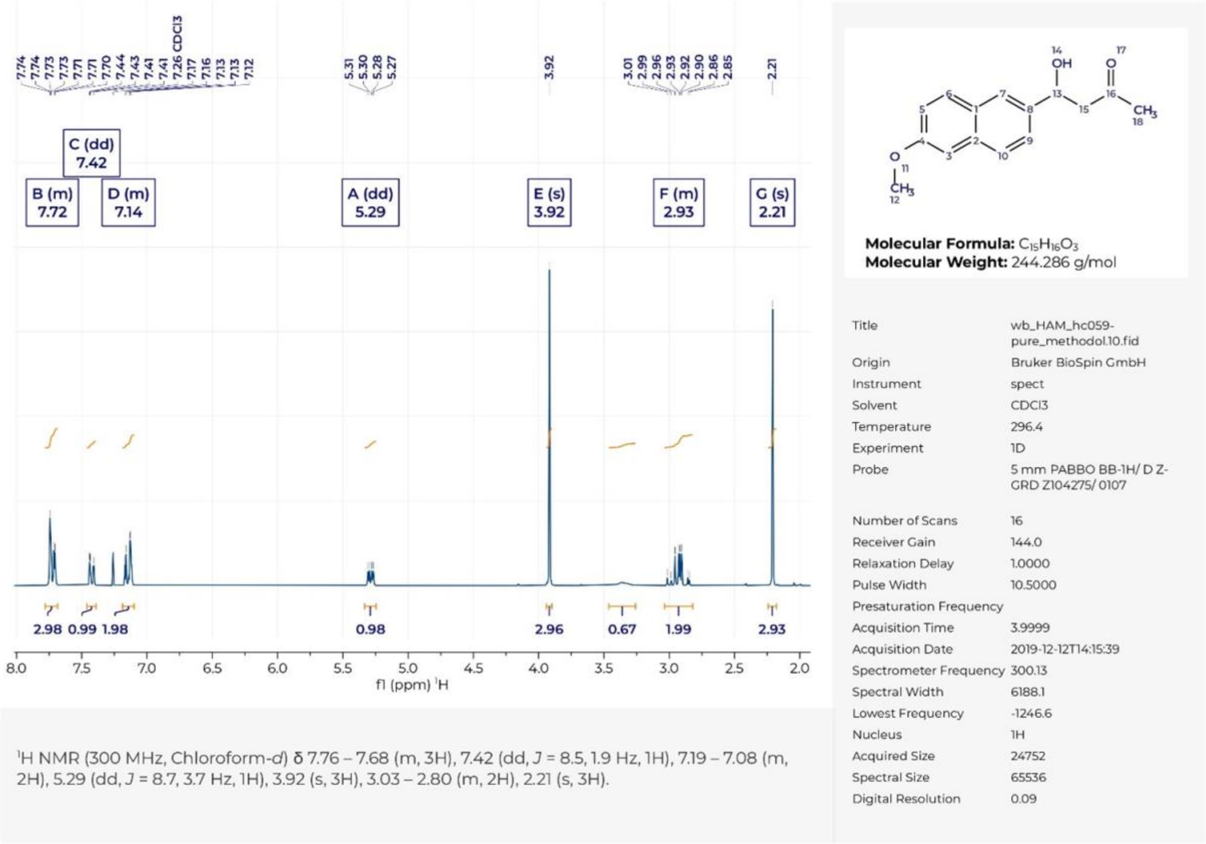

**Fig. S18.**

**$^1$H NMR spectrum of *rac*-methodol 1.**

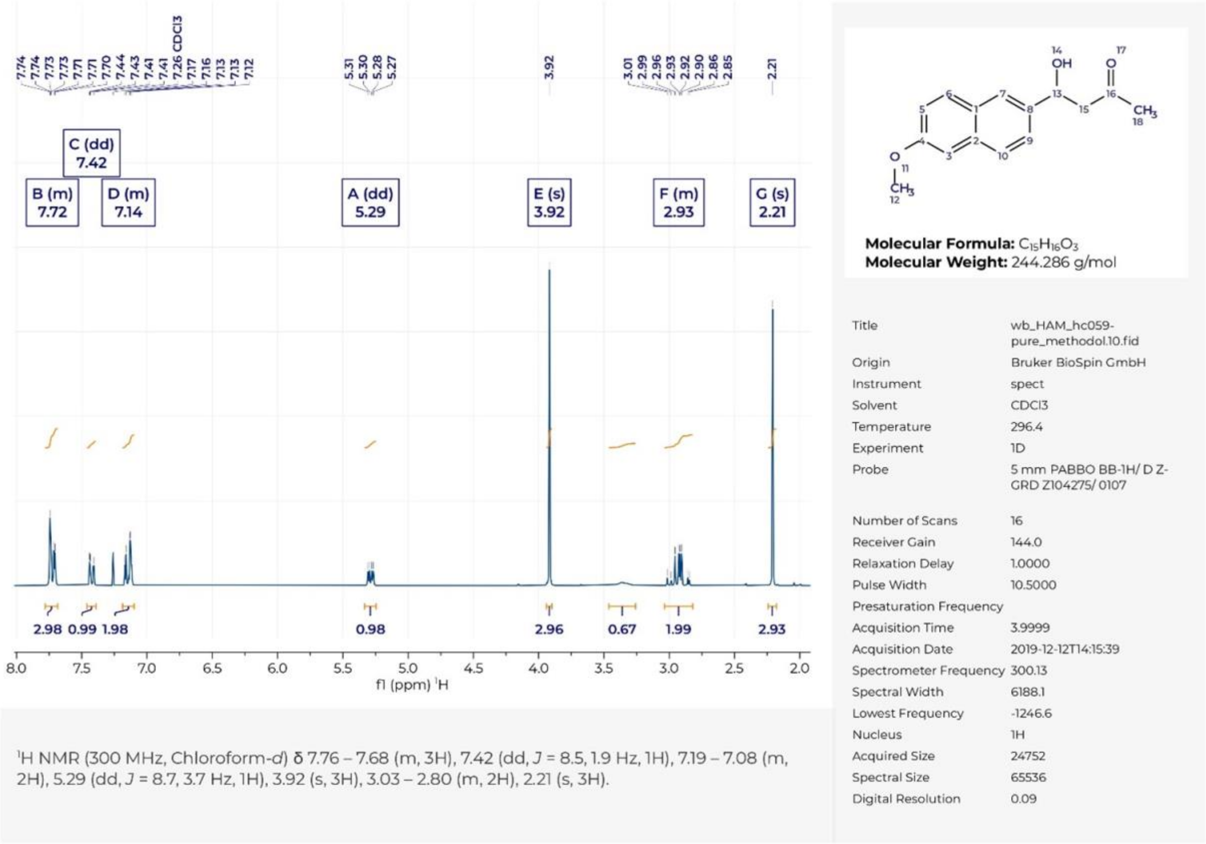

**Fig. S19.**

**$^{13}$C NMR spectrum of *rac*-methodol 1.**

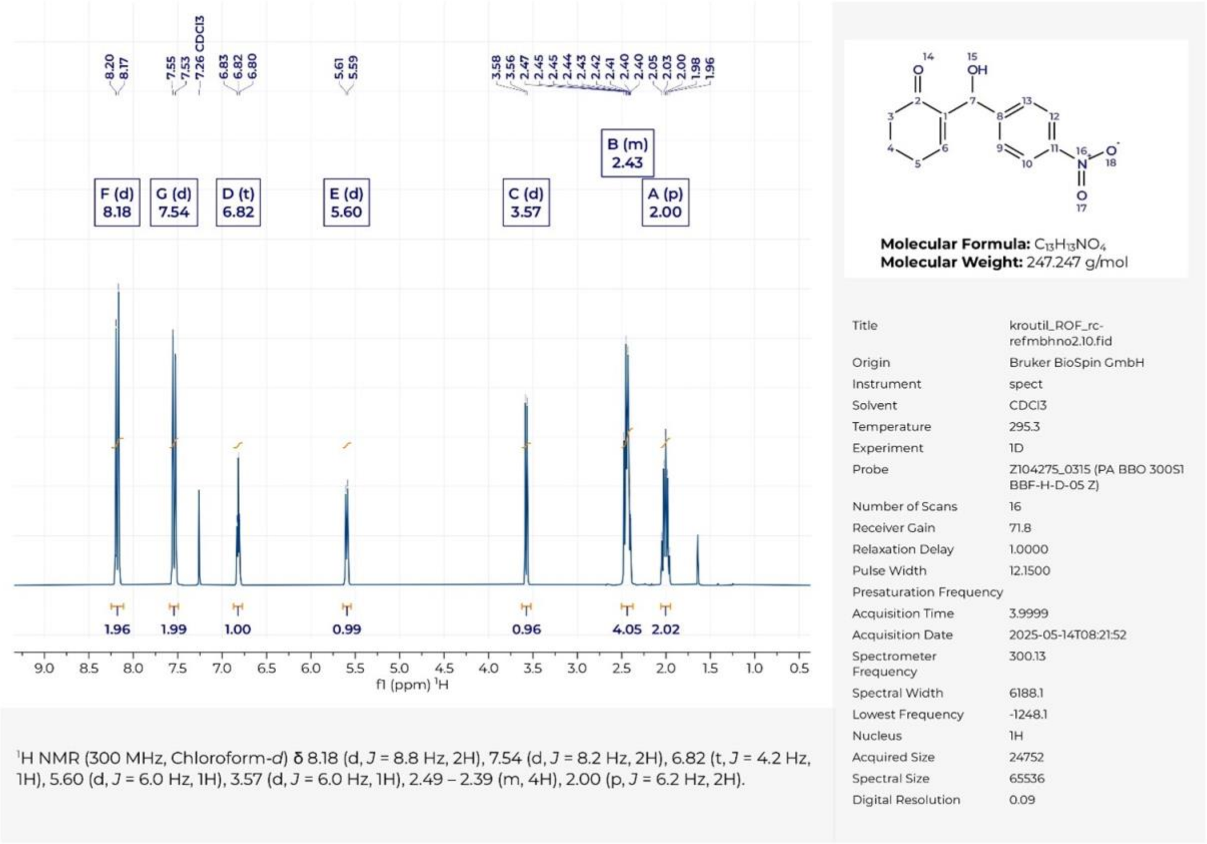

**Fig. S20.**

**$^1$H NMR spectrum of 2-(hydroxy(4-nitrophenyl)methyl)cyclohex-2-en-1-one 5.**

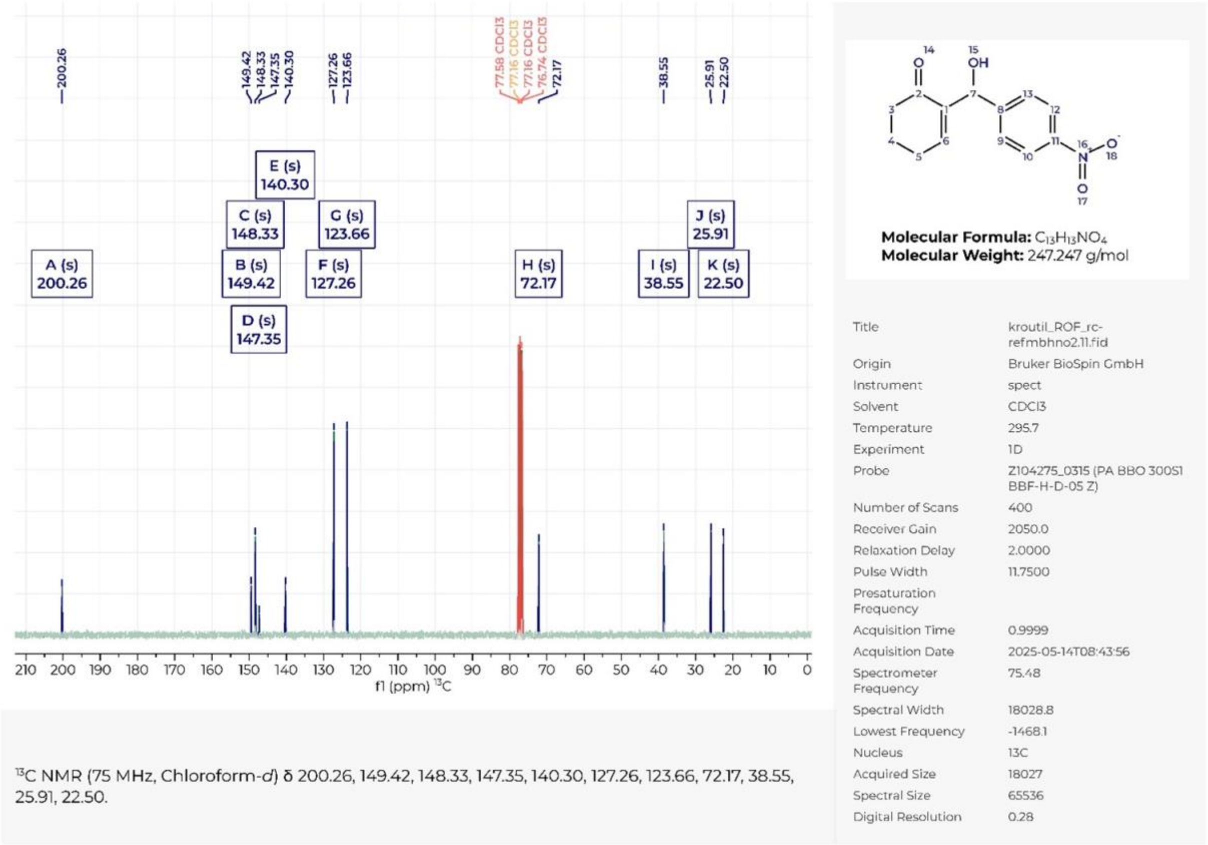

**Fig. S21.**

**$^{13}$C NMR spectrum of 2-(hydroxy(4-nitrophenyl)methyl)cyclohex-2-en-1-one 5.**

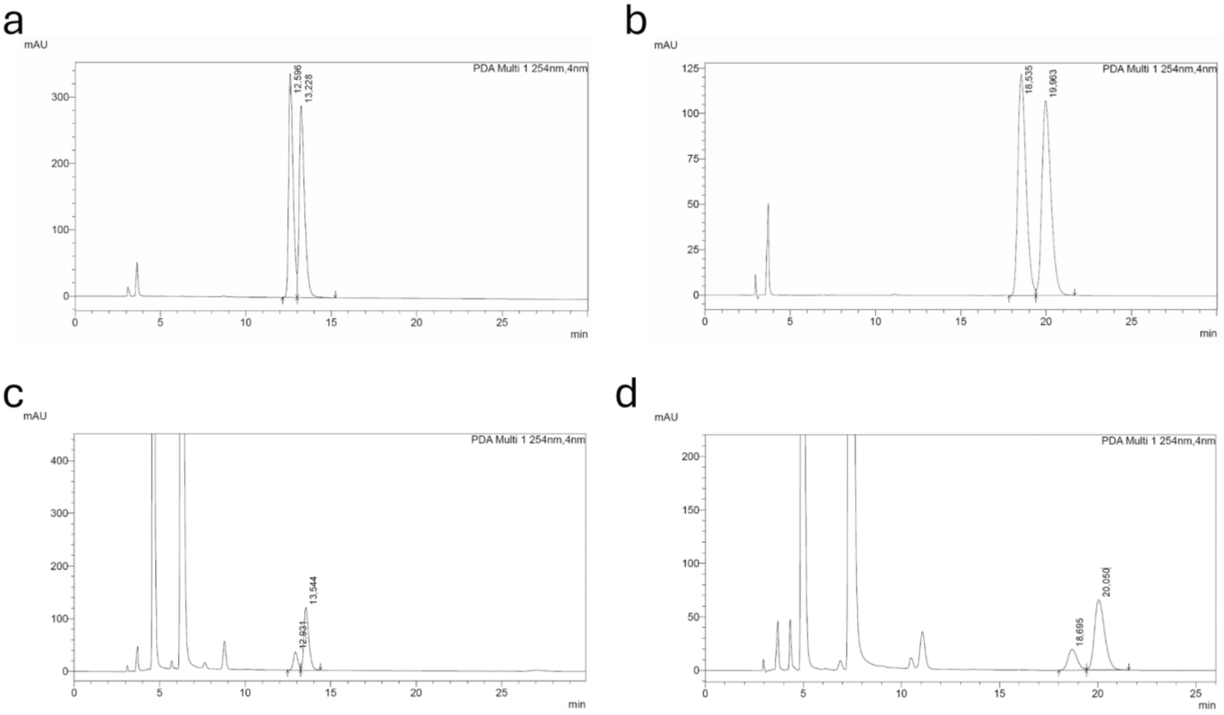

**Fig. S22.**

**HPLC analysis for determination of methodol absolute configuration and enantiomeric excess. a.** and **b.** HPLC traces of freshly synthesized *racemic* methodol obtained on a Daicel Chiralpak IB column (**a**, retention times: (*S*)-methodol 12.6 min, (*R*)-methodol 13.2 min) and a Daicel Chiralcel OD-H column (**b**, retention times: (*S*)-methodol 18.5 min, (*R*)-methodol 20.0 min). **c**. and **d**. Representative traces from samples of the aldol reaction with RAD29 showing formation of the (*R*)-enantiomer with 60% ee on a Daicel Chiralpak IB column (**c**) and a Daicel Chiralcel OD-H column (**d**) obtained under the same conditions as in **a.** and **b**.

**Table S1:**

**Amino acid sequences of RAD constructs.** All genes were cloned using Golden Gate cloning into a vector featuring a N-terminal hexa-histidine tag and a TEV cleavage site with the sequence MGSSHHHHHHSSGENLYFQSG. Extinction coefficients were calculated using ProtParam.

| RAD | sequence | extinction coefficient [$M^{-1}$ $cm^{-1}$] |
|---|---|---|
| 1 | MSEEEKRKKEEEEEYKKKLEEAREYLEKTMGKFTSLLLKAAGREDLIPTYAPLSSEYVKNVPDEKLLEI KEKNEKLGEKAKEKEEEFEEYKKLLLELASKDLSPEEAKLVGLGSLAFFLGVGKYEGMEKFYELWEEF KKVASPSVLDALREFFAGLYEKIAKSKTLEEFLEGVFEGYVEFMEFIFENYDELLEKVKELAK | 20400 |
| 2 | MSKEELEKLKKEEERKKEIEEARRYLERTMKRFTTLLLKAAGREDLIPELAPAASEYVKNIPDEELLEILK KNKELGEKAKKLEEEFEKLKETFLTLAGKDLSPEEARLVGLGSLAFFLGIGKAKGEEYFYELWEEFVKV ASPAVLEALREFFRGLFEKINKAKTKEEFLKGVYDGYKEFMDFIFNNFEELVEKVLELSK | 14440 |
| 3 | MEKLKEILIELAKLAGDPSEESQKEYLGKAIAFMGALALAALDPSLQSPEALAELTEELLPLGEEGAKTL SEEEIKEITDFNLSMASYFGDMTPEELVAAVAAAPSALLALTIAVGGIKGVAVKDPSKVPEYVEAFLEGL GTWAAKHPEEWHAFFAAMGEEILADPSKLAENMAKVYKDLVDELKKLLPEAVKLYHEVKA | 18450 |
| 4 | MERLKELLIELAKLAGDPSEEKQREYLGKAIAFLGALALAALRPELQSPEALAELTEELLPLGEKGAETL SEEEFQFISDFNLSAADDFADATPEELVEAIVAAPSIELALAIAVLGIKGVAVKDASKVPEYVEAFLRGLG EWAKRHPEEVRAFFEGAGRRLVEDPSTLAQRMADLYKELVKRLKELRPQAKALLEEVKK | 9970 |
| 5 | MEKLKEILIELAKLAGDPSEESQREYLGKAIAFLGALALAALDPSRRSPEAIAQLTEELLPLGREGAKTLS EEEFKYITEFNLKAGEYFKDVTPEELVKAVVAAPSPELALTIAVSGIKGVAVEDKSKVPEYVDAFIEGLG EWAKKHPEEVAAFFEAMGRELLADPSTLAERMARVYRDLVKELEKLLPEAVALVEEVKK | 12950 |
| 6 | MEKVREILIELAKLAGDPSEEAQRKYLGKAIAFFAALALSALDPALRSPEALAALFEELLPVGEEGAEGL SEEEKEKIFRFNREVGLKFKDVTPEELVAAIVAADSPELALTIGLGAVKGHAVLDPSKVREYVEAFLRGL GEARARHPEAWDAFFRAAGRRLLADPSRLAERAAGLYADLVSRLESLLPEALALVEEVKK | 9970 |
| 7 | AEEEAKKIIEKGRKFMEEHGREYVEKVLEEVGKFLAKVAKDPVLLAGLTKLDPRFANLNLSREERLELYI WIAGFGNKVAAEALREEGLEKAAEIFEKASKKALKGVELAKEKGVEEAIKYALEANAENGGAALLALKE SGKGVPLVERLLRKAKEEPEKAGEYFLGALVLGFSIIKGLAFDPDNEVLREAAERVRELLG | 11460 |
| 8 | LSPAEREELRRRLRLLAIFGAKGLLNRGEELEVTPEDEPYLERGRELAEKGIELYEEYPEVIDYLKPHLE ELKRINEEESLEFLEVIEEAPEELQPLLRREMAFAYAALSTLREKAKELGEDEDLNYYGLAAVHQALMR VLEEVRREDPSLSEEEALEKAIERTRELLKNPKEIADYYVEATYRMLEEARALGEEIRGLH | 14900 |
| 9 | MEEALEASREAIELDPEGAKEVAELNREAGRIVEEAGSYEEVAKKVLEAAKEGKLSDETIIAAAKGLVY TPEGREVARKTAEEAEKLAKESEGEERKRLALLSFLLRLLVEIYEKSKDDEGYLVLAGVHWLAAKIAKK ELEKNPEWYYDVEGIEEAFKKGLEEAAKAPREEIIKAGLDYFKEAKKIMEKGNKELKELLFK | 21430 |
| 10 | MEKALEAAREAIKENPEEAKEVAELNREAGKVVEEAGSYEEVAKKVLELAEKGKLSDDAIIAAAKGLTY TPEGVEVALKTAEEAKEKAKKSEGEGKERLTLLSFLLELQAELTLQSKDDEGYLTLVTVYWLAAKIAKE YLEKNPEASTDYEGIKEAFEKGLEEALKAPAEEIKKAGFDYFKNAKKILKKGNKELRELLFE | 15930 |
| 11 | MEKALEAARKAIEANPELAAKAAELNKKAGEIVEEAGSYEEVAKKILEAAKEGKLSYNAIIAAAKGLAYT PEGQKVALETAEKAEEEAKKSSGESKEKLILLSFLLRLQVKLKEESEDDEGYLTLATVYWLAAKIAKKKL EEDPSLAESLEGIEKAFKEGLEEAEKAPEEEIIKAGKDYFENAIGIMTKGNEELKKLLFS | 14440 |
| 12 | MEKALEASREAIKLNPELAKKVAELNEEAGRIVKEAGSYEEVAEKILEAAREGKLSRETIIAAAKGLVYT PEGQEVARATAERAEEAARESEGEGRERLTLLAFLLRLLVELYEASEDDEGYLVLAGVHWLAAQIAKK KLEENPEAALDIEAIEKAFEEGLEEAAKAPEEEIIKAGLDYFEEAQKIMEKGNKELKELLFS | 12950 |
| 13 | MEKALEAARKAIEEHPEEAKEVAELNKKAGEIVKEAGSYEEVAKKVLELAREGKLSDDAIIAAAKGLAY DEEGQEVALKTAEEARKAAEESSGKGKERLTLLSFLLRLQVRLTRESEDDEGYLTLATVYWLAAKIAK KKLEEDPSASTDLEGIEKAFEEGLEEAKKAPEEEILKAGFDYFEKAKEIMEKGNKELRELLFK | 12950 |
| 14 | FEKALKRGREFYEKDPEGFKKMMELNEELADLAYKLIEEGKIEKIVELAAEYYKKGDELTAALVLAALVK GIALHDLPEEYVEKAAEYAKKLIGEEWGEFIRGLVAFLRKLKEAFPDPEELHLAYFGYVTALFLAIERLK ENPELYKDKPYFDRILLTAEEILNDESTLPELEKYAEFYLTEVMEKLEKIKKELEKILKE | 23380 |
| 15 | FEKALEYGRRFYEEDPEGFERMMKLNFELAKLAKELVEKGKIREVIELAAEYYKKGDELTAALVLAALV KGMALYDLPEEYVEEAAAYAEELIGPEWGEFIRGLVAFLRELKKAFPDPEELALAYFGFVTFLFLAIDEL KKDPSLYADEPFFDRILKKAEELLNDESTLPKAREYARFYLTEVMEKLEEIRKKLEEILEK | 21890 |
| 16 | FEKALEWGRYFYAKDPEGFKKMMDLNEEMAKLGAELVKQGKIEEVVELAAEYYKKGDELTAALVLAV LVKAMALYDLPEEYVEKAAKLAEEKIGPEWGEFIRGLVAFLRELKAAFPDPEELALAYFGFVTALLLAIE ELKKNPELYKDKPFFERILLTAEEILNDPSKLPELEKYAEFYLTEVMKKLEEIRKKMEEILKK | 25900 |
| 17 | MTPEEEARAAVDSFPEALRQRAWDLNVKSAEKLAKYGIEKVTELALKLLKEIFEKYVEGKITREDLPEV VKKILVLLSLVKATAIYSKEGLEKILELLKEIAKELRERGETLLAEAIDYLIEALEKLHKGDADGYLTLLTIAL YLYFKHIVENGARDPELAAAVRPLVEGGYEAVARYYFEVFAPKLEEGTEEAVKLFEE | 20400 |

| | | |
|---|---|---|
| 18 | MSDEELARAAVDSFPEAERQRAWDLNVKSAEKLAKFGIEKVTELALEKLKEIYKKFIEGKITKEDLPKLI<br>EEILILLALIKATAIYSEEGLEKILEEYKKIAEELRKKGYTLLAEAIDYLIKALKALHAGDADGYLTLLTIALYL<br>YFKHIVDNGKFDPELAEKVRPLVEGGYEAVARYYFEVFAPKLEEGTEKAVKLFEE | 21890 |
| 19 | MSDEEEAAAAVHSFPQELRDRAWNLNVESAEKLAKYGIEKVTERAVELLKELYENFVEGKITREDLPK<br>VLEDLLVLLALVKATTIYSEEGLEKIIALLKEIAEKLRELGYTLLAEAIDYLIEALKKLHEGDADGYLTLLTIA<br>LYLYFKYIVENGHLDPELAALVKPLVEGGYEAVARYYFEVFAPKLEEGAEKAVKLFEE | 23380 |
| 20 | MTDLEKAREAVLSLPKELRERAWKLNVESAEELAKFGIEKVLEKALELTKELAELYFEGKITPEDLPEVA<br>KKMLILLAYIKAATIKSKEGLEKIIKLYEEIAEELRKRGETLLAEAIEYLIKALEALHKGDAEGYLSLLTVALY<br>KYYKWLVEKGDKFPELAALHKPLVEGGYEAVARYYFEVFAPAFEKGAKEAVESFKK | 27390 |
| 21 | MTDEERALAAVRSLPEELREYAWKLNVESAEKLEKFGIEKVTEIALKKLEELSELYFEGKITPEDLPEVF<br>RKMLILLAYIKATAIKDEKGLEKILEKLREIAEELRKKGETLLAEALEILIEALKALHKGDADGYLSLLTIALH<br>KYFQWVVSQGDKFPELAAAVQPLVDGGYEAVARYYFEVFAPKFEEGAKEAVEEFKK | 22920 |
| 22 | NLLEFWIKFLEGLVKSLSEEGREDYREGLGGLLRRAARGLEKLDPSLKEAIELVRRAAEKAGPNLTDEE<br>ILEINLEAGRAGKIVVEKGVEKVIDNGIKLIKENPLDLELLLEVTAALTVIAKGAYFDPSVVELLKKKAEEL<br>KKEGDDFTAFILELLAETGEKFAELAKEENGGEKYLEWLFTVVREEIEKKIEELKKKLE | 15470 |
| 23 | NLLEFWIEYLRELVDSLSEEGREVYKNSLGETLKLAAEGLKLADPSMKEAIELVEKAAEKAGPNLTDEEI<br>GEINLKAGEAGEIVMKKGYKPVVDKAIELWKKNPLDKEILLKVTNAITVIAKGSYKYPEVVDYLREKAKE<br>LRKEGDDFLALILELLAETGEKWAELAKEKEGTKKYLEWLFSKVKGRIEEKIKELEKELK | 32430 |
| 24 | DLKKFYIEYLEGLVESLSEEGLKVYKEGLGELLTRAAEGLELLDPSLKEAIEKVREAAKKAGPNLTDEQI<br>KEINLEAGEAGKIVIEKGYKPVVENGIKLIKENPLDLELLLKVTNALTVIAKGAVYNPEVVEYLKKKAEELE<br>KEGDTFTAFILRLLAETGEKWAELAKEENGGKKYLEWLFTEVRKEIEERIEELKKELE | 21430 |
| 25 | NKELQEIADKIYNGLKSLSGKEYYEWVKNAPEKFEKNKKIFLEDPSKLDFDGFFTVITATLKGVLGKSKL<br>EEELKKLAEIIEKTVSLLSKVNPFWLKLAEAGLLFEEIAKKLYKENPEIVEEYFESLGNTFEEALEVVEKL<br>NTIPRGVLFILLAGMAIHASKFKELSKEEIEEVFEFNKKMAEIGLPFIEEALEALKKKE | 18450 |
| 26 | MEELVEKALELAPKKAKENRKPYFDFVMKYVEDPELLRRTLAEGARKFAEAVALRRGEEAERLLAEAA<br>FAVIVGAKGYVVAGDKERLDELKAMLEEMAESVRTAQLLVAAFGIIEKLIDDPKKYIGAITAFLGTVEELE<br>KEGPLSLEELKAIAEAIIENADKLVDVSDEEVAKMDAFNKRMGEIAIPLLEAALEALRASL | 5960 |
| 27 | MEEYEKILDEIVNGLRNMSGEEYYEMVLNAEKYYYPAVEKFLEDPSSLNGDEFFTVFTLSLKGVLAAG<br>DLDTLLPKYAKIIEETIKEMSKKDPFWLDFAKAALKFEKVFQEEYEKNPEVVDEYFRANGVSPQEALDY<br>ALKFDDISRGVLLILLAGMAIQGLKLLSLSEEQREKVFAFNRSMKDIGLEFVDKALEALEKKK | 20400 |
| 28 | MEKLEELAKKILDFLESMSGQDYYEMVKNFEEYFEKAKKKFLEDPSKLNFGEFFTVITYTLKGVLGKAD<br>LEEEFPKLAEVIRETLEILSKKDPFWLDLAEAGLLFEEVAEKYYKEDPEIVEEYFSSLGVSFEEALEVALK<br>FDKIPRGILLILLAGMAIHARKLKDLSEEERKEVFEFNKKMAEIGMKLVEEALKELEKKK | 15930 |
| 29 | MEELVEKALELAPEAAKENAQPYYDFVKKYIEDKELLKKEIEEGSKNFAKALELKEGEEAEELLKKSAF<br>AVIVYAKGLALSGDEEKFEELLEKIKEMAESLETARLLYAALLIIKKLIKDPKKYFGAITAFLGVVERLEKE<br>RPLSLEELVAIAEAIIERAEELQAFTDEQVEEMRKFNEEAGKLAIEYFEKALEELKKKK | 10430 |
| 30 | MEELVKRALELAPRTAKERREPYFEFVSRYIDDPELFRREMEEGARLFEEALRLRTGPEAERLLREAA<br>FAVIVGAKGFAVAGDEERFNRLVEALREMAEDVPTARLLVAAFEIIKRLIDDPEKYIGAVTAFLGVVEEL<br>EKERPLSLEELVAIAEAIIERADELVDVSKEEVERADAFNRRAGEVAVPYLDRALDAVEAER | 5960 |
| 31 | MMEKAEEAIKEAIENRDERVEPYVERAKKLEKEIKKYVEAGVEAIVQAIKDGDLLVLAMLLKGLFYHGFY<br>GKEEEAIELLERLAEAVPTLEGRLMALLYAELLRYVKEKGISKEEFAPLYLTVILILLPTYKKLVEEGVLSE<br>ETSLEELREIIRLVIERTPEPSEKIKKATEEVEPINEKMGELLSFEEMKEVVEGVANS | 11920 |
| 32 | MLEKAKEAIKEAIENRDELVEPYVERAKKLAEEIKKYVEGGVEAIVEAIKNGDLEVLAMLLKGIFYHGFY<br>GEREEAIELLEKLAKAVKNLEQRLMSLLYAELLRYMEEKGISWEEFAPQYLTLITILLPTYEKLKEAGVVT<br>ESTSLEELREIIKLVLENLPEPSELEKEATKEVEPINKKMGEYLSFEELKEVVEGVANG | 18910 |
| 33 | MMEEAERAIEEAIKEIDERAEKYVERAKWMIEEVEKYVEAGLQAVIDAILNGDMIVLAMFLKGIFFHGLK<br>GKVEEAIELLRRLAEAVPYLEGRLIALMYALFLEYVTEKGLSLEESYPIYLSVILVLLPTFEKLVEKGVLSE<br>ETSLEELEELIRLVIEKTPEPSEKVKEYVKEVAPINEKMGELLSFEEGREVVEGVAAA | 17420 |
| 34 | MREAVEEALARARERRDELVGRYMAWAREFYNNPELQEEILEGAKEAIRDPSDPEKMKALGIALAIAIK<br>GAVFSSEEEIERARRLLERLGKLSPEHAKLAEFMLALLDAMVTLRETLSREDFALAYLGLLMVVLLALS<br>EGDVETATAALRLVAEGEYEPFLELAAPYREEAEEALRINEGLVPLGLRFLEELERVVEEVE | 12950 |
| 35 | MEEAALKAIEEAIKNREKIVKEYVERAKEMIEKLKEYVKKGKEAIVDAILNGDMDVLAMFVKGIFFHGLR<br>GREDEAIELLERLAEAVKTLEGRLIFLTYAEMLRYIRERGISREEFWPLYLSLITVLLPTFERLVKEGVLS<br>EETSLEELRDLIRLVWERTPEPSELERKATEEVAPINEEAAELISFEEGKEVVEGVASA | 18450 |
| 36 | MRERVEEALRRAKERREEIIGKYLEWAKTFANNPELQKEINERALKAIKDPSDEKDLKALGIALAIGMKG<br>PIELGEEAVEELLGLLERLGKLSEKHAELADFLKALVQAYMTLKKTLSEEEYRVTYLGMIAVVLLALSEG<br>DYDTAKAALELVVEGDYEPFLELAEPYAEEAKEAWEINKKLVEYGLKVLEKMKEAIKEVE | 22920 |

**Table S2.**

**Mass spectrometry results of purified enzymes.** The expected molecular weight includes the serine-glycine stub that remains after cleavage of the histidine tag. For RAD14 and 15 this cleavage was incomplete, resulting in a higher detected molecular weight. The highest intensity peaks for each sample are displayed.

| RAD | expected MW [Da] | detected MW [Da] | difference expected – detected [Da] | Intensity |
|---|---|---|---|---|
| **1** | 23569.65 | 23569.1 | -0.55 | 898358 |
| **2** | 23536.89 | 23536.4 | -0.49 | 1552472 |
| **3** | 21670.68 | 21670.3 | -0.38 | 869407 |
| **4** | 22140.16 | 22139.7 | -0.46 | 880003 |
| **5** | 22113.19 | 22112.8 | -0.39 | 479635 |
| **6** | 21888.87 | 21888.3 | -0.57 | 1287525 |
| **7** | 22219.38 | 22218.8 | -0.58 | 1140131 |
| **8** | 23571.27 | 23570.7 | -0.57 | 1373084 |
| **9** | 22658.54 | 22658.3 | -0.24 | 1371438 |
| **10** | 22359.14 | 22358.7 | -0.44 | 1251987 |
| **11** | 22022.86 | 22022.4 | -0.46 | 1514071 |
| **12** | 22286.9 | 22286.4 | -0.5 | 1379381 |
| **13** | 22435.1 | 22434.7 | -0.4 | 1472734 |
| **14** | 23538.93 | 25618.9 | 2079.97 | 609901 |
| **15** | 23491.77 | 25571.7 | 2079.93 | 582547 |
| **16** | 23285.91 | 23285.5 | -0.41 | 432873 |
| **17** | 22817.1 | 22816.8 | -0.3 | 1184266 |
| **20** | 22932.46 | 22931.9 | -0.56 | 1579678 |
| **21** | n.d. | n.d. | n.d. | n.d. |
| **22** | 22664.84 | 22664.5 | -0.34 | 1445679 |
| **23** | 22926.24 | 22926.2 | -0.04 | 1074630 |
| **24** | 22777.9 | 22777.7 | -0.2 | 1691445 |
| **25** | 23115.53 | 23115.1 | -0.43 | 348040 |
| **26** | 22236.5 | 22236.2 | -0.3 | 1418615 |
| **27** | 23162.25 | 23161.8 | -0.45 | 1202863 |
| **28** | 23505.96 | 23505.5 | -0.46 | 1027103 |
| **29** | 23068.37 | 23067.8 | -0.57 | 1292346 |
| **30** | 22920.77 | 22920.6 | -0.17 | 2171495 |
| **31** | 23043.51 | 23043.1 | -0.41 | 721705 |

| | | | | |
|---|---|---|---|---|
| **32** | 23038.34 | 23037.9 | -0.44 | 1392019 |
| **33** | 22899.36 | 22898.9 | -0.46 | 369553 |
| **34** | 22729.77 | 22729.7 | -0.07 | 1379768 |
| **35** | 23187.47 | 23187 | -0.47 | 1061567 |
| **36** | 22885.14 | 22885 | -0.14 | 1450476 |

**Table S3.**

**SAXS parameters.** %MW and %Rg denotes deviation from expected molecular weight and Rg calculated from the design models.

| RAD | FoXS chi$^2$ | FoXS c1 | FoXS c2 | FoXS default chi$^2$ | BI MW [Da] | %MW | Size& Shape MW [Da] | %MW | Rg Guinier [A] | %Rg | I0 Guinier | Rg p(r) [A] | %Rg | I0 p(r) | Rmax (p(r)) [A] | Porod volume [A$^3$] | Porod Volume % | model Rg [A] | model Porod Volume [A$^3$] |
|---|---|---|---|---|---|---|---|---|---|---|---|---|---|---|---|---|---|---|---|
| **1** | 2.3 | 1.04 | -0.22 | 56.5 | 18675 | 79% | 22769 | 97% | 20.4 | 100% | 82.23 | 20.6 | 101% | 82.36 | 70 | 40995 | 140% | 20.44 | 29300 |
| **2** | 1.2 | 1.04 | 0.05 | 35.6 | 18050 | 77% | 22625 | 96% | 20.2 | 99% | 72.11 | 20.2 | 99% | 71.87 | 60.7 | 39406 | 148% | 20.44 | 26710 |
| **3** | 5.3 | 1.02 | 4.00 | 167.4 | 15475 | 71% | 20906 | 96% | 18.7 | 118% | 74.82 | 18.3 | 116% | 74.09 | 53.4 | 30020 | 119% | 15.82 | 25280 |
| **4** | 2.3 | 1.03 | 2.92 | 78.4 | 16125 | 73% | 26379 | 119% | 18.8 | 117% | 54.44 | 18.5 | 115% | 54.15 | 56.9 | 29298 | 109% | 16.06 | 26770 |
| **5** | 39.5 | 1.03 | 4.00 | 311.7 | 16125 | 73% | 23062 | 104% | 20.5 | 127% | 90.13 | 20.6 | 128% | 90.1 | 70.6 | 40165 | 161% | 16.09 | 24930 |
| **6** | 4.4 | 1.01 | 4.00 | 57.8 | 14825 | 68% | 19010 | 87% | 19.2 | 121% | 49.84 | 19.2 | 121% | 49.7 | 64.9 | 30541 | 126% | 15.9 | 24240 |
| **7** | 4.4 | 1.01 | 4.00 | 57.8 | 14150 | 64% | 19163 | 86% | 19.5 | 118% | 46.24 | 19.6 | 119% | 46.14 | 64.6 | 31931 | 140% | 16.49 | 22800 |
| **8** | 8.7 | 1.01 | 4.00 | 62.4 | 20550 | 87% | 23865 | 101% | 18.5 | 110% | 65.05 | 18.4 | 109% | 65.09 | 53.7 | 36386 | 118% | 16.85 | 30830 |
| **9** | 1.6 | 1.02 | 1.94 | 58.6 | 15475 | 68% | 21417 | 95% | 18.5 | 111% | 68.58 | 18.1 | 109% | 70.18 | 51.7 | 31146 | 126% | 16.68 | 24800 |
| **10** | 1.2 | 0.99 | 3.91 | 64.2 | 15475 | 69% | 19492 | 87% | 18.1 | 108% | 76.92 | 18.3 | 109% | 68.44 | 56.4 | 30491 | 110% | 16.73 | 27690 |
| **11** | 1.1 | 1.01 | 1.94 | 46.7 | 14825 | 67% | 17008 | 77% | 18.1 | 109% | 76.92 | 17.9 | 108% | 76.77 | 51.5 | 30660 | 108% | 16.64 | 28340 |
| **12** | 1.4 | 1.01 | 3.61 | 56.3 | 12750 | 57% | 20623 | 93% | 18.6 | 111% | 72.12 | 18.4 | 110% | 72.0700 | 55.8 | 30681 | 134% | 16.71 | 22940 |
| **13** | 1.1 | 0.99 | 3.87 | 43.0 | 13450 | 60% | 18462 | 82% | 18.3 | 110% | 62.33 | 18.1 | 109% | 62.1900 | 55.2 | 29068 | 106% | 16.68 | 27330 |
| **14** | 8.1 | 1.01 | 4.00 | 116.0 | 16125 | 69% | 20722 | 88% | 19.6 | 119% | 67.23 | 19.1 | 116% | 66.0800 | 57.8 | 33931 | 115% | 16.46 | 29380 |
| **15** | 6.0 | 1.03 | 4.00 | 109.9 | 13450 | 57% | 16186 | 69% | 19.2 | 118% | 57.48 | 18.6 | 115% | 56.8000 | 50.3 | 34180 | 118% | 16.24 | 29010 |
| **16** | 2.4 | 1.02 | 4.00 | 62.6 | 19925 | 86% | 25396 | 109% | 19.5 | 119% | 57.05 | 18.9 | 115% | 56.3800 | 53.1 | 31014 | 107% | 16.38 | 29080 |
| **17** | 2.6 | 1.01 | 4.00 | 57.2 | 23675 | 104% | 24839 | 109% | 19.8 | 120% | 62.11 | 19.3 | 117% | 61.2700 | 56.6 | 35300 | 110% | 16.5 | 31970 |
| **18** | 1.7 | 1.03 | 1.15 | 32.0 | 17425 | 76% | 19323 | 84% | 18.6 | 113% | 47.37 | 18.3 | 111% | 47.2400 | 52 | 30492 | 97% | 16.52 | 31460 |
| **19** | 1.6 | 1.02 | 2.68 | 36.3 | 17425 | 76% | 24283 | 107% | 19.1 | 116% | 42.96 | 18.6 | 113% | 42.6500 | 55.4 | 32238 | 104% | 16.45 | 30980 |
| **20** | 1.2 | 1.04 | 0.06 | 22.3 | 18050 | 79% | 20954 | 91% | 18.4 | 111% | 44.26 | 18.1 | 110% | 44.1100 | 55.2 | 27037 | 85% | 16.51 | 31740 |
| **21** | n.d | n.d | n.d | n.d. | n.d. | n.d. | n.d. | n.d. | n.d. | n.d. | n.d. | n.d. | n.d. | n.d. | n.d. | n.d. | n.d. | n.d. | n.d. |
| **22** | 1.2 | 1.04 | 0.94 | 55.2 | 17425 | 77% | 21821 | 96% | 18.5 | 112% | 62.2 | 18.2 | 110% | 62.04 | 54.9 | 29807 | 102% | 16.54 | 29200 |
| **23** | 2.3 | 1.02 | 4.00 | 26.1 | 12025 | 52% | 20084 | 88% | 20 | 121% | 27.97 | 19.5 | 118% | 27.5500 | 58.1 | 26666 | 92% | 16.58 | 29080 |
| **24** | 1.9 | 1.02 | 3.03 | 83.4 | 15475 | 68% | 20592 | 90% | 18.9 | 114% | 58.39 | 18.7 | 112% | 58.2100 | 64.8 | 32596 | 117% | 16.63 | 27930 |
| **25** | 4.9 | 1.00 | 4.00 | 77.9 | 16775 | 73% | 19283 | 83% | 19.5 | 118% | 54.98 | 19.4 | 117% | 54.6800 | 67.9 | 29514 | 100% | 16.55 | 29600 |
| **26** | 1.5 | 1.04 | 0.08 | 43.9 | 14150 | 64% | 20927 | 94% | 18.3 | 109% | 65.11 | 18.1 | 108% | 65.0700 | 52.1 | 29963 | 106% | 16.75 | 28270 |
| **27** | 9.9 | 1.01 | 4.00 | 318.4 | 15475 | 67% | 20809 | 90% | 19 | 115% | 128.59 | 18.8 | 114% | 128.0000 | 60.4 | 36547 | 128% | 16.47 | 28650 |
| **28** | 9.7 | 1.01 | 4.00 | 292.3 | 22425 | 95% | 25748 | 110% | 19.5 | 119% | 136.86 | 19.3 | 118% | 135.9 | 66.3 | 40775 | 149% | 16.38 | 27360 |
| **29** | 1.4 | 1.01 | 1.88 | 30.0 | 16125 | 70% | 22828 | 99% | 18.9 | 111% | 68.73 | 18.6 | 109% | 68.5000 | 60.9 | 31562 | 101% | 16.99 | 31330 |
| **30** | 1.7 | 1.01 | 1.13 | 17.0 | 14150 | 62% | 17680 | 77% | 18 | 107% | 73.84 | 17.8 | 106% | 73.7600 | 58.6 | 26777 | 87% | 16.81 | 30940 |
| **31** | 1.5 | 0.99 | 4.00 | 39.0 | 18675 | 81% | 18878 | 82% | 18.4 | 112% | 65.45 | 18.1 | 110% | 65.2200 | 57 | 28780 | 99% | 16.48 | 29020 |
| **32** | 1.6 | 0.99 | 3.71 | 56.4 | 17425 | 76% | 18043 | 78% | 18.1 | 111% | 91.11 | 17.9 | 110% | 91.0600 | 48.7 | 29801 | 102% | 16.3 | 29130 |
| **33** | 1.8 | 1.00 | 3.18 | 88.3 | 17425 | 76% | 18264 | 80% | 18.3 | 112% | 77.34 | 18 | 111% | 77.1900 | 52.2 | 32135 | 110% | 16.27 | 29200 |
| **34** | 1.3 | 1.03 | 0.64 | 80.9 | 15475 | 68% | 17680 | 78% | 18.2 | 112% | 79.02 | 17.7 | 108% | 78.5200 | 70.1 | 27927 | 94% | 16.32 | 29630 |
| **35** | 2.5 | 1.02 | 1.26 | 85.4 | 15475 | 67% | 17737 | 76% | 17.6 | 109% | 105.12 | 17.3 | 108% | 104.8000 | 57.3 | 28979 | 105% | 16.08 | 27540 |
| **36** | 1.2 | 1.02 | 2.80 | 40.8 | 12025 | 53% | 19237 | 84% | 18.4 | 112% | 55.63 | 18.2 | 110% | 55.5600 | 51.1 | 30402 | 99% | 16.49 | 30710 |

**Table S4.**

Michaelis Menten parameters for the retro-aldol reaction with *rac*-methodol, chemical denaturation midpoints, expression levels and pKa1 of retro-aldolases. ± indicates 95% confidence interval of the fit.

| RAD | kcat [s-1] | Km [µM] | denaturation midpoint [M GdnHCl] | yield [mg/L culture] | pKa1 |
|---|---|---|---|---|---|
| **1** | 5.7E-05 ± 0.3E-05 | 710 ± 80 | n.d. | 16.3 | n.d. |
| **2** | 1.6E-05 ± 0.1E-05 | 190 ± 20 | 4.7 ± 0.1 | 16 | n.d. |
| **3** | n.d. | n.d. | 5.3 ± 0.1 | 2 | n.d. |
| **4** | 4.0E-05 ± 0.4E-05 | 1000 ± 200 | 4.4 ± 0.1 | 3.7 | n.d. |
| **5** | 2.6E-05 ± 1.6E-05 | 1900 ± 1600 | 4.5 ± 0.3 | 7.9 | n.d. |
| **6** | 2.80E-05 ± 0.4E-05 | 600 ± 150 | 4.50 ± 0.01 | 17.4 | n.d. |
| **7** | 3.6E-05 ± 0.5E-05 | 870 ± 200 | 2.5 ± 0.2 | 16 | n.d. |
| **8** | 3.3E-04 ± 0.1E-04 | 520 ± 50 | 3.3 ± 0.2 | 21 | n.d. |
| **9** | 3.1E-04 ± 0.1E-04 | 340 ± 30 | 3.5 ± 0.2 | 12 | n.d. |
| **10** | 3.6E-05 ± 0.2E-05 | 390 ± 50 | 2.7 ± 0.1 | 5.2 | n.d. |
| **11** | 6.8E-05 ± 1.1E-05 | 330 ± 130 | 3.1 ± 0.1 | 9 | n.d. |
| **12** | 1.2E-04 ± 0.1E-04 | 150 ± 50 | 3.1 ± 0.2 | 17.5 | n.d. |
| **13** | 3.8E-05 ± 0.5E-05 | 320 ± 110 | 2.6 ± 0.2 | 9 | 9.0 ± 0.2 |
| **14** | 5.6E-06 ± 1.4E-06 | 440± 240 | 6.1 ± 0.1 | 1 | n.d. |
| **15** | 7.9E-06 ± 1.2E-06 | 640 ± 190 | 6.5 ± 0.1 | 6.5 | n.d. |
| **16** | n.d. | n.d. | n.d. | 1.5 | n.d. |
| **17** | n.d. | n.d. | n.d. | 28 | 7.4 ± 0.1 |
| **18** | n.d. | n.d. | n.d. | 3.4 | n.d. |
| **19** | 3.6E-06 ± 0.3E-06 | 410 ± 90 | n.d. | 2.4 | n.d. |
| **20** | 2.6E-05 ± 0.2E-05 | 990 ± 110 | n.d. | 20 | n.d. |
| **21** | n.d. | n.d. | n.d. | n.d. | n.d. |
| **22** | 3.6E-04 ± 0.5E-04 | 590 ± 170 | 4.7 ± 0.1 | 3.4 | n.d. |
| **23** | 6.5E-05 ± 0.5E-05 | 710 ± 110 | n.d. | 4 | n.d. |
| **24** | 3.7E-04 ± 0.5E-04 | 210 ± 80 | 3.2 ± 0.1 | 14 | n.d. |
| **25** | 4.5E-05 ± 0.5E-05 | 280 ± 70 | 4.2 ± 0.2 | 32 | n.d. |
| **26** | 1.1E-03 ± 0.2E-03 | 68 ± 35 | 3.4 ± 0.3 | 40 | n.d. |
| **27** | 2.4E-04 ± 0.1E-04 | 660 ± 70 | 3.2 ± 0.2 | 28 | n.d. |
| **28** | n.d. | n.d. | 4.5 ± 0.2 | 14 | n.d. |
| **29** | 3.1E-02 ± 0.2E-02 | 110 ± 20 | 5.1 ± 0.1 | 18 | 7.0 ± 0.2 |
| **30** | 1.5E-02 ± 0.2E-02 | 210 ± 90 | n.d. | 32 | n.d. |
| **31** | 1.8E-03 ± 0.2E-03 | 670 ± 160 | n.d. | 11 | n.d. |
| **32** | 1.1E-02 ± 0.1E0-2 | 1400 ± 120 | n.d. | 22 | 8.0 ± 0.3 |
| **33** | 2.2E-04 ± 0.2E-04 | 940 ± 50 | n.d. | 16 | n.d. |
| **34** | 2.3E-03 ± 0.4E-03 | 190 ± 80 | n.d. | 22 | n.d. |
| **35** | 3.7E-02 ± 0.1E-02 | 350 ± 30 | n.d. | 16 | 7.6 ± 0.1 |
| **36** | 6.2E-05 ± 0.2E-05 | 540 ± 40 | n.d. | 27 | 8.6 ± 0.3 |
| **29 N178A** | 9.6E-04 ± 0.4E-04 | 120 ± 10 | n.d. | n.d. | n.d. |
| **29 Y120F** | 7.8E-03 ±1.6E-03 | 100 ± 40 | n.d. | n.d. | n.d. |
| **29 Y23F** | 1.2E-02 ± 0.1E-02 | 65 ± 9 | n.d. | n.d. | n.d. |
| **32 Y120F** | 2.3E-04 ± 0.5E-4 | 960 ± 340 | n.d. | n.d. | n.d. |
| **32 Y99F** | 7.8E-04 ± 0.9E-04 | 190 ± 60 | n.d. | n.d. | n.d. |
| **32 N178A** | 3.6E-03 ± 0.5E-03 | 330 ± 120 | n.d. | n.d. | n.d. |
| **35 N178A** | 1.9E-04 ± 0.2E-04 | 350 ± 90 | n.d. | n.d. | n.d. |
| **35 Y120F** | 1.5E-04 ± 0.1E-04 | 620 ± 80 | n.d. | n.d. | n.d. |
| **35 Y23F** | 1.8E-03 ± 0.1E-03 | 870 ± 120 | n.d. | n.d. | n.d. |

**Table S5.**

Comparison of designed and evolved Retro-aldol enzymes.

| | substrate | pH | $k_{cat}$ ($s^{-1}$) | $k_{cat}$ /$K_M$ ($M^{-1}$ $s^{-1}$) | $k_{cat}/k_{uncat}$ | Ref. |
|---|---|---|---|---|---|---|
| **Computationally designed retro aldol enzymes (this work)** | | | | | | |
| RAD1 | *rac*-methodol | 7.4 | 5.70E-05 | 8.02E-02 | 8.77E+03 | this work |
| RAD2 | *rac*-methodol | 7.4 | 1.60E-05 | 8.21E-02 | 2.46E+03 | this work |
| RAD3 | *rac*-methodol | 7.4 | n.d. | n.d. | n.d. | this work |
| RAD4 | *rac*-methodol | 7.4 | 4.03E-05 | 4.00E-02 | 6.20E+03 | this work |
| RAD5 | *rac*-methodol | 7.4 | 2.59E-05 | 1.37E-02 | 3.98E+03 | this work |
| RAD6 | *rac*-methodol | 7.4 | 2.80E-05 | 4.68E-02 | 4.30E+03 | this work |
| RAD7 | *rac*-methodol | 7.4 | 3.56E-05 | 4.09E-02 | 5.48E+03 | this work |
| RAD8 | *rac*-methodol | 7.4 | 3.29E-04 | 6.34E-01 | 5.06E+04 | this work |
| RAD9 | *rac*-methodol | 7.4 | 3.06E-04 | 8.97E-01 | 4.71E+04 | this work |
| RAD10 | *rac*-methodol | 7.4 | 3.59E-05 | 9.18E-02 | 5.52E+03 | this work |
| RAD11 | *rac*-methodol | 7.4 | 6.76E-05 | 2.05E-01 | 1.04E+04 | this work |
| RAD12 | *rac*-methodol | 7.4 | 1.23E-04 | 8.26E-01 | 1.89E+04 | this work |
| RAD13 | *rac*-methodol | 7.4 | 3.75E-05 | 1.17E-01 | 5.77E+03 | this work |
| RAD14 | *rac*-methodol | 7.4 | 5.60E-06 | 1.26E-02 | 8.62E+02 | this work |
| RAD15 | *rac*-methodol | 7.4 | 7.93E-06 | 1.24E-02 | 1.22E+03 | this work |
| RAD16 | *rac*-methodol | 7.4 | n.d. | n.d. | n.d. | this work |
| RAD17 | *rac*-methodol | 7.4 | n.d. | n.d. | n.d. | this work |
| RAD18 | *rac*-methodol | 7.4 | n.d. | n.d. | n.d. | this work |
| RAD19 | *rac*-methodol | 7.4 | 3.55E-06 | 8.70E-03 | 5.46E+02 | this work |
| RAD20 | *rac*-methodol | 7.4 | 2.55E-05 | 2.57E-02 | 3.92E+03 | this work |
| RAD21 | *rac*-methodol | 7.4 | n.d. | n.d. | n.d. | this work |
| RAD22 | *rac*-methodol | 7.4 | 3.55E-04 | 6.01E-01 | 5.46E+04 | this work |
| RAD23 | *rac*-methodol | 7.4 | 6.50E-05 | 9.22E-02 | 1.00E+04 | this work |
| RAD24 | *rac*-methodol | 7.4 | 3.65E-04 | 1.76E+00 | 5.62E+04 | this work |
| RAD25 | *rac*-methodol | 7.4 | 4.50E-05 | 1.59E-01 | 6.92E+03 | this work |
| RAD26 | *rac*-methodol | 7.4 | 1.13E-03 | 1.67E+01 | 1.74E+05 | this work |
| RAD27 | *rac*-methodol | 7.4 | 2.44E-04 | 3.70E-01 | 3.75E+04 | this work |
| RAD28 | *rac*-methodol | 7.4 | n.d. | n.d. | n.d. | this work |
| RAD29 | *rac*-methodol | 7.4 | 3.14E-02 | 2.93E+02 | 4.83E+06 | this work |
| RAD30 | *rac*-methodol | 7.4 | 1.51E-02 | 7.16E+01 | 2.32E+06 | this work |
| RAD31 | *rac*-methodol | 7.4 | 1.79E-03 | 2.66E+00 | 2.75E+05 | this work |
| RAD32 | *rac*-methodol | 7.4 | 1.11E-02 | 7.82E+00 | 1.71E+06 | this work |
| RAD33 | *rac*-methodol | 7.4 | 2.22E-04 | 2.36E-01 | 3.42E+04 | this work |
| RAD34 | *rac*-methodol | 7.4 | 2.26E-03 | 1.19E+01 | 3.48E+05 | this work |
| RAD35 | *rac*-methodol | 7.4 | 3.66E-02 | 1.05E+02 | 5.63E+06 | this work |
| RAD36 | *rac*-methodol | 7.4 | 6.18E-05 | 1.14E-01 | 9.51E+03 | this work |
| **Computationally designed retro aldol enzymes (from existing scaffolds; no laboratory evolution)** | | | | | | |
| RA34 | *rac*-methodol | 7.5 | 1.22E-04 | 1.90E-01 | 1.90E+04 | [4] |
| RA45 | *rac*-methodol | 7.5 | 2.83E-05 | 3.60E-02 | 4.00E+03 | [4] |
| RA60 | *rac*-methodol | 7.5 | 1.55E-04 | 2.70E-01 | 2.40E+04 | [4] |
| RA95 | *rac*-methodol | 7.5 | 3.33E-05 | 5.30E-02 | 4.80E+03 | [4] |
| RA110 | *rac*-methodol | 7.5 | 8.33E-05 | 4.80E-02 | 1.20E+04 | [4] |
| cRA-50 | *rac*-methodol | - | 2.2E+00 | 1.8E+02 | - | [5] |
| RA117 | *rac*-methodol | 7.4 | 2.76E-05 | 5.8E-02 | 4270 | [6] |
| **Computationally designed retro aldol enzymes (from existing scaffolds; laboratory evolution)** | | | | | | |
| RA34.6 | *rac*-methodol | 7.5 | 3.67E-04 | 1.20E+01 | 5.50E+04 | [4] |
| RA45.2-10 | *rac*-methodol | 7.5 | 2.32E-04 | 5.40E-01 | 3.60E+04 | [4] |
| RA60.2 | *rac*-methodol | 7.5 | 1.17E-03 | 1.80E+00 | 1.80E+05 | [4] |
| RA95.4 | *rac*-methodol | 7.5 | 2.50E-04 | 8.40E-01 | 3.40E+04 | [4] |
| RA95.5-8F | *R*-methodol | 7.5 | 1.08E+01 | 3.4E+04 | 1.7E+09 | [7] |
| RA110.4-6 | *rac*-methodol | 7.5 | 3.83E-03 | 5.50E+01 | 5.90E+05 | [4] |
| RA95.5-8 | *rac*-methodol | 7.5 | 1.7E-01 | 8.50E+02 | 2.6E+07 | [8] |
| RA117.4 | *rac*-methodol | 7.4 | 2.1E-02 | 5.6E+01 | 3.23E+06 | [6] |
| RAβb-16.1 | *S*-methodol | 7.5 | 2.50E-02 | 1.08E+02 | 3.8E+06 | [9] |
| RAβb-16.2 | *S*-methodol | 7.5 | 2.67E-02 | 5.0E+02 | 4.1E+06 | [9] |

**Table S6.**

**Amino acid sequences of MBH constructs.** All genes were cloned into a vector featuring a N-terminal hexa-histidine tag and a TEV cleavage site with the sequence MGSSHHHHHHSSGENLYFQSG. Extinction coefficients were calculated from these sequences using ProtParam[10].

| MBH | sequence | extinction coefficient [$M^{-1}$ $cm^{-1}$] |
|---|---|---|
| 1 | GEAARRAGETFGEVLNAGDEANPALAPLAEANRRFRREHADLVERIYALGAEAIERLGTADPAAARAV<br>VAVMLGLLNVARLAAELRDEGKDEEADRLLDLAEEILRAALAGTPEEVITVANAVGQAAWLAYIAGKRA<br>DLALENLEKVRNANLEEQKAFAEAGAAASVAWLAATYGPEAAAAHLAAVARAVDAAVDVLRAA | 15470 |
| 2 | GEAARRAGEAFGEVLDAGDRANPALQPIADENRRFVRDHADLVERIYALAAEAIARLGTADPAAAAAV<br>VAVMLGLLNVARLAAHLRDQGRDAEAEELLALAEEILRAALAGTPEEVTTTARAVGQAAWLAFVAGRS<br>VELALENLRKVKEANLEERRAFAEEGEAASVRWLADTYGAEAAAAHLAAYAAGVAAAVAILQAS | 15470 |
| 3 | GQEFADAVAEAGRIIIRHDRASPHPALQAHAVFSAAFGERNGDTIAAEIVAGKTDFRAGFAAWNATLE<br>RVAALHPEYAPLLEKVIALNEEMTAHLDAAPVTSEEEAAMRAFNAAIGVGNRLRAADPALDTYISTEEG<br>AEAALIGGVAMLLFRGKFAGPEYQAKAEEILAQLPPELQAAVHNHLAGLEEAFAHFKKVYLSL | 11460 |
| 4 | GQARADAVARAGEIIIRHDRASPHPALQVHAEYARAVADEHGDEIAARIVAGENDFVAGQQAWVDTLA<br>QVAALHPELAPLLDQVAALKREMTAHLAAAPITSDDEAAARVFVADIGVANRIRAADPRLAAYIATREG<br>AEAALIGGVAMLLFLARFAGPEFAARAEAILAQLPPELQAAVHQHLAGLEEAFAYFKEVYLSL | 11460 |
| 5 | GPPPPGPVSPEGQAARDGHLAGLEAGREVVREAVRAFLPEDPEAADLIADLAWALFQEYLARLRAQA<br>ETAEEALATIPLAFELAARGAVEIYNAVTGRLTPEEAAALVEVLREALRRGYAVLLAFDLSFSDEAYAEG<br>MEAMRNDFAANYDAIMAALRAELEAETDPIRRATIQAEIDGREAWNANLDHITKLTIEGRKIL | 18450 |
| 6 | GPPPPGPVSPEGQAAADGHREGLEAGREVVREKVAAFLPEDPEAAEKIADLAWAKYLMYLAKLRAQ<br>AKTAEEALATIPLAFELAAKEAVDIYNAVTKRLTPEEAEELKKVLYEALKEGYAVLLAYDLSFSDEAYAE<br>GMELLRRTFAANYDAIMAALKAELEAETDPIRKATIQAIIDGKAAWAANLDHITELTIEGRRIL | 22920 |
| 7 | GRLRRIVALSRKILEIHGAGLKQMPDSEFAALLADLALAYADYVQALVDGDDAAAAAAKARVDALTARI<br>NADPVGRAALETWAATLRKEVPAYAAALEEARAAEAAGDRAALAEALARALAYEAIAIVRNIENKDEVL<br>AARRAAYEELVPKAVGRPELEVLAEMYRWLAEIAAHPKHRAIDDKYAEELKKLAEELERLLA | 21430 |
| 8 | GAAAAGYYVREGIAVVRGELEAELTPEELAAFDALFARIEARVEPLITAGDPPLALVRAAFYAANNTLR<br>GLDPAILRLFGATDEEIAAAQAVARKLLVEFERALAERDRLTDEELTARLEAIAVEAWNAIAPSALRSPY<br>PVLRAAALAGARTTPALSAEDAALRAAALAAATPEELAAADAHDAAFAAVDARAQELYDAA | 12950 |
| 9 | GAAAAGYYVREGIAVNRGELERELTPEQLAHWEALFEKIVAEVEPLITAGDPPLALVDAAFYAANNTLR<br>GLDPGILRLFGATDAEIAAAQAVAERLLVEFREALARRDELTDEELTARLRAIAVEGWNAIAPAALRSPY<br>PVLRAAALAGARRTPALSAEDAALRAAALAAATPEELAAADAHAAAHDAVDARARELYEAA | 18450 |
| 10 | GLRELLGRHLAGAALLDGAIVDPALVPQAVAFLQQPNPALSPVQRQAVIDALRALETGLTPEQRAGIRA<br>YLEANVKERAADPARLEAAEALAIVLGQKSLGVTDIPADLVAKAEVARTAWDEETASYADVTDAIADKV<br>VEIGTTRLGIDADLARSTGRYGGLFVALGILKDPELAVHIAIVHAASLILIGYAALIEQQLA | 11460 |
| 11 | SALLERAKAKAKELGEIWKSMLGPVFKKVFDADPERFFKANLGGLLYYVMVAERGAEAADAELRKISP<br>EVAAGIEADRDILEGNPELRKEIIERGQEYIDRLKVEDPERLKKIEDLFEKILEIRREFLLIGGMSEEETDA<br>FLEASRGFELLYVFFLLYIDRLLQEDPEKGLALVKKFAPYFVAHFNRFIILAEKYEAEAK | 15930 |
| 12 | GGKIWGIAHAGIISLDIGRAETTDPRLLEFIDLARELLRAVARAALAMPEEEVEKMWEFFKQVVPEERLT<br>AAPPVPDVPLTEEEFEEYLNRAIELAEELARRFEGELLELLAEHGRRHASSLTPEELARVEKANKLILEE<br>LEKIRTEISKVGTPEQIATANRIIDFIRKLIETAPYSAIVWGTILGLLMLLWVLLWKAKQ | 30480 |
| 13 | GDAAVAGELREGAEKADGAIIGDYDIKEVAQELLDLAKRYGNPTLVKIAEIGLNNIDTEITPKGRENYRKI<br>IETAKVERAANPELKEKVEKFQYLFGRAVLAPVEEIPALMAEAVKHSQWWLDDPFGRAYLALMINVVIK<br>ANPEKGRAIADAFGFDDADLAAGVAAMGALGITEAIAAIHLTGNVLAALAEDFAARAAAA | 18450 |
| 14 | GAALAGAEAAAGAVLAARHAARPEAVAAEKEIEAAPDMAAAVEAAARYFRAGLDDYAAEIRAEADSP<br>AASPRASAFMRALADAMREGAALMAEDPIAGLYALILRVEAIIKADPEAAFEASDFHFSLPEYAAFRPV<br>SERKAKEIAARVPEIVAEAGLPADVQAAMVYVAEVAAEAGGGPLWPFLEIIEALERAVAAVKAK | 12950 |
| 15 | GGAEFGAMIAGTGAAVDGLRALPGGEAAAFAVDVLGQNEILAYMVELAAGDPDIAAEFGILVVAREVA<br>HIYGVLRDDPEAAERLGNAFLDVALGAKTVEELRAGVLAVYRALMEHLAKRDPSWLDRLALWEEVVD<br>KVEPESYWTNFLALVNGALARYSREEIRARALELAREAVATGDADAAMDARLVAAVERAEARKAA | 23950 |
| 16 | GGLELLGAYARYGLEIAGKEVAADKERYEELSQLLVEVGAALDEALAALRAGDYERAVEIILDTLEKHR<br>EKLTYLGRRHEKYLLENKDDPKLAKDVASYENLKKSLQAIVDGDLDTAADYYRKSLEAVRDRYVALHP<br>DGETVYQEFVDRITNTIRNANPETYAILKRVSEFLLEAGISGLLWALLEVLLYALSGVQEAAQ | 24870 |
| 17 | GGLYEFAGRFAAGAVERGKLWKTVTKATYHYGVDYVVGKALQLGREEPELAASYRALARGLIRERLY<br>LDLYHAGRRDEAIEIIQGHYGVSREAAERILDRRVAEEKEYLASPPPTDEEVTAALRGLAGYRKRLTQA<br>VLDRFGREDDVILVQLYLDLSSDPELLADFLSHSDAEIVASSRGIDDRLQPVYDARLAEIEAKL | 23380 |

| | | |
|---|---|---|
| 18 | GVAEAAAGTRAYYGARHGVGLLEGILAAFPDLPAEDRAYLEDALAYYRGLAEKYQPVVDANPEVSAAI LDFIAEQLLASLRARIESGENPVLELAAGVVRTLEDYARAVGKPEAIEVGKRVALLVAKTGAPTIWLSLG RADAELLPKYREEIIAGYQRLAELAVKYAREVTMEEIEAAIAKWEAAQPAIDAAFQAVVDSL | 25900 |
| 19 | GDGLARVEAAFGALADALAALAGEDPLAGVLAEWVRLIPPVLYRDDPALALALADLLAAAVADDPVAA ARAFLDANRYIDAVDDEVAAKDPAARKAILDAGWAYLRGGGDRQPLLDAARAHPEAFRRHCLASVEA SRRSVRRARADPARARAYTAVAVLMGVFGVVDAEALLEAGVGNQELVDEARGALDEARALLEELL | 16960 |
| 20 | GLRELLEELGAIERAGLANLVAHPERAAVIRGTDAANLARFRATGNEAFVELRKRFYLSIANAIKRAFG WSEEEMRTWLATAGADPALADRLPDDISDEQLAYFLSAELVVGCTYALGGDPADVRALGRRLFSDW LDFASDAPIVQELLSTFSPEEIELFIDGLVDHAVAGSVRFADEEAVRRALEVGERGVAWAKSLKA | 26470 |
| 21 | GMEKVEELSAESVRVTATELVRLAREDPAAARAHAEAGKVVGKKVYEACQELAARSTSERVRTLLRFI GENELIGAEMDEMALAGASVEELVAFAEERSAEIAAPLEGWPEGFLLTFNIAFNTGWLAGVTGDEELY NKALEYFEKVGIPREDAEKILDIRLEMAKADAADGSPVLAAAIARITEVTLKLIKAEEEALNSL | 15470 |
| 22 | SPLAALLREVAAAILGGDPERGLQLAKKLWEGLWWLGVDPARGEEILAEVKAATGLTAEIDTLAAGVR ELTRTVIEADERLRAEDPAAYAAIMAFAKLITTLKLYPEEVSPEEAKAAELFLKGEELIEVYLKGGPESYL TQGKKYVGEAAKILGLDPEVAWRGAVIHAEGVVKMTPEEAAALAAMDATLEEAIAAAEARL | 29450 |
| 23 | GMEEVERLSEELVRVTATELVRLAREDPAAARAHAEAGEVIGKAVHDACKELAAQSDSERVKTLLNFI GENELIGARMDRMALAGASVEELVDFAKKNNAKIAKPLEGWPKGFLLTFLIAFYSGWLAGVTGDEKLY NLALEYFEKVGIPREWAETILDIRLGMAKADAADGSPVLAAAIARITAVTLELIKACEKALNS | 21095 |
| 24 | SPAAALARELLDAILGGDPERGLALAKKLWEGLWWLGFDPARGDAILAEVKAATGRTAEIDTLAAVVR ELNRTQVAADEELKKADPKAYAAIKAFADLVTTARLYPEKVTPEEREAAELFLRGERKIREYLRGGPEE LLEQGRVDVARAAEILGLDPDVAWRGAVVHAQGSVAQTPEEAAALAELDATMAAAIAAAEAR | 26470 |
| 25 | GEGLARVEAAYGALADALAALASRDPRAGVLAEWIRLQPPVLYKMDPALALKLAELTARAVARDPEAA ARAFLDANRKIDAVDAEEAAKDPAATADILAAGWAYLRGGGDMQPLLEAARAHPEAFRRHCLASVEA SRESVRRARADPERALAYTSVAVLMGVNGVVDALALLLAGVGDQGLVDLARGALDEAEALLDELL | 16960 |
| 26 | SAAAALLQRLLDKILGGDPVEGLKLAKKLYEGLWWIGFDPARGNKILDEVRKATGLDAEINALAAVRRK LNATVIAADEALRAADPAAYAAIRAFADLVTTLKLYPEKVTPEQAEAANLFLAGEDLIAQYLRGGPESLK TKGVEYVARAAKILGLDPAHAIRGAFVHVQGLVDRTPEDRAALAAMDATASAAIAAAEARL | 18450 |
| 27 | DPAHTALEAAYRAVVGVLGEAVVDAFLKATALAEIAGFRLLLEGADLETALAAAREAVRAYLPTLVEAT GNRAVAEAIAEAAFAHAELGLRVAEVLRADPALFRAYLEDALKVNEAMVEDPRKGLRLFVDFSKKYGK LWLIAVGKLHGAKTDEEALKLGKKYSPLVDLRGALRIFEAYAAATGRGAELAAATRAEIERLE | 14440 |
| 28 | MKAVEEAEKRYEEIVSVLGEAVVDALWKATALAEIAGYEALLAGADLAEALALARQAVEDYLPTLVEAT GDRAVAEAIAEAAFAHAELGLRVAEVLRADPALFRAYLEDALKVNRAMVADPREGLRLFVEFSKKYGK EWLIAVGKLHGAKTDEEALKLGKIYSPLVDLVALYKVFEAYAKATGKGKELAEETKKKIEETL | 22920 |
| 29 | MEELEKYLAALAERKKAVDARVAERLAKIDPAEWARAAAPIGLRLADLVEELGFPQEVVDAMRADPAV AGDVLSRAVVEALAERVRQELAAQGLAELGEIVANLLITEADVLRQELALEFEDMYHAGATLDDLAAR ADELAAALRAAAPSELVGLILEAAVLGGLLHALMGIDIELGGKAWPYIERTVRRLYELYQVATA | 18450 |
| 30 | GEALRARALELGRELGEAFRRSISKLTPAEIQAHADAGAALAAVPVDPEAFLAAARAYMRVDAEVGER AGLCDSADELFARGLENVELALGGVKELGLESRLPELVAELVALTKAGAAKDKELADDTAAYADYVRD LVVGLVQAVVDASRTLPAEEAVFLAALVALWIAIAATLSDEVVEAGRRVIEALLDLARYLKAAS | 11460 |
| 31 | DLRELARAVIRALDAVHQALIATSDEGAQYVRDWYATLLAALDRLGLGDVRDEVLAAIERHGLAAILDV VLGTLTPEQIDKAVELGPVHAETGLALQEAVFGGASPEEVAAAKEKVAAADAEVMAQLRPLITEEQVE AVAEALGVSRRAALALILLTLVIYGGTVAEVALLVEGTPEQKARVAGLLAQLAAEYAALLARL | 11460 |
| 32 | GDGPAGLLAALAEMRAAGASPEELFFFACSVILEALQTDPVLGQAMLDALATLPEEVQEIALEVSRLHL EGTLVHQDELDAARAAVAAATAALLAALAAAGADAAQLAAARERLAQAVTGAWFRGAGYPHVAARY QPYLDWARAADPALAAALDALVAVGEKIAEFDVEAMKLVRKITDEELLRRATETLVAALRRVAAQ | 15470 |
| 33 | GEELRERALVLGRELGQAWRDSVLQLTPEEIRAHAEAGAALAAVPVDPAAFRAAARAFMRVDAEVGE RAGLCDSAEELFARGLENVDLALAGVDELGLAARLPELVAELQRLTRAGAATDFELADDTAAYAETVE RSVVSLVQSVVDASRELPAEEAVFLAALVALWIAYAATISDAVVEAGRRTIEALLELARYLKAAS | 15470 |
| 34 | PVDAAGVLAALDEMRAAGASPRDLFLFAAGVVLEFLLYDPAAGRAMLEALAGQPAEIQKLALQVSEM HLAGSLKYKEELAAALARVEAAAAALLARMRAEGASDAELAAARRLLADARRAAWYRGAGYDFVAER YQPALDWARAADPALGAALDEYEAVGKKIAKYDVEAQKLVDAITDEAALTAATEALRAALRRVAAA | 21430 |
| 35 | GADELAQRLGELIRGGLANLKAHPERAAVIRDTDAQNAARFRATGDQRFADLVDRFYLAIANAIIRTFG MTDEEAETWFRLGGGDPADLSKFEDTISRDHLLYFLSIELVVGCTHALGGDPKDVRALGRKLFKDFLR YASDAPIVQELLEGFTPEQMELFLDALVDHAVAGSVRFADEAQVREALEVGREGVAWAKRRAA | 15470 |
| 36 | SPELAEKLKKKAEEMRAWLEEHAAEVRRTTLAVERALARELLAEARAAGDARAAAVWEAYQAVLAWL SDPANYALWLRVAAANPRLVVSRAVDDLAVGVLAVICLDMTREEIEENKEELIAIYAVLHATQSIAVLET VIATPELADRWVGMLDLSPEERARALVDIWAEASNSEEGKKRITEAINKVLKEIAKELEAKKK | 37470 |
| 37 | GPGFTPEEQALVEELAKKGEELTKAIFEAASKHERAYYEAQGKPELYELILAHGERALKDGAARVAGL GDRLLTAEGQIELGRAYVKAAPMDFLHFKDAIETLGLEDKIFNGNLHEFLTALLKAVDYMLKNADVYGK RHAEQGVAATERMIEEGVDLTDYDAFRAAWERLNKEVFPEEYEEALTYFKKLEEVIRRLHAEL | 18910 |

| | | |
|---|---|---|
| 38 | GQESVDFGRRGLELLAEKKLYKSLAAYYAALHIELLLLPLALSVLPIPQEAVEAFLGIYRAAQEILAKNPE<br>AAKLVPDMIPYGLAHVAGWAGTTAINVQIAQEAQLTSPEKRIKYLKAWLNGFIAGFVSPEEAEKLIKEG<br>AELLSEIVPKELAERYYEASLVGAAADQQATDGILTPEERKATAECFEKAVEEVIAKIKA | 22920 |
| 39 | GVEEAARRLLDIYLEALQVTVEHADEFIKYLPIHVKGSEEGTKLYAAGDYVGATRTIISADREIAEIIGNE<br>EAVKLFDELLAEAERRAALRRELYAEAGLTDADVVAILQAGVDYDRSLLTYSPEEAAALLRKETQAEID<br>ALRARLYGDRPLTPEELRQRAAFVAAWLLGGGTAAAIFDPKLYRRMLDAALAFHAALKAS | 18910 |
| 40 | CPGDTPEEAALRRELAARGAELTRAVNDAAFKYERAYYEEKGEPERYERLKRFGERALAQGAARVA<br>ALAPELCTAEGRVAFGKDYVVASPVDVLNLKDAIETLGLEEEIENGSKEQFLRALLRAVDYMLEHADEY<br>GKKHAEAGVAATEEAVRRGVDLDDFDAFSAAWKELNKEVFPEEYKEALEFWAKIDEVIRRLAAER | 23045 |
| 41 | MKELLEKLKKRTEEMKEFLKKHEEEVRKKTFEVERKLAKELLAVAKAAGDARAAAVWEAYIKRLDWLA<br>DPENLALFERVAAAEPRLVVSRAVDDLAVGVLAVITLDMTKEEIEENKDEIIEIYAKIHAAKSVAVLDTVV<br>ATPELAERWVGNLDLTPEERAREYVAIWAEASNSPEGRRRIAERIGALLVEIAKKLAAKLL | 26470 |
| 42 | GEASVRFGRLALQNLGKAKLYKAMAATYAALLVERALLPFALRVLPIPEEAVAAYRGIYDAAEEILAKYP<br>EAAKLVPDMIPYGLAHVAGWAETSVLNVRIAREAKLTTVEKRIAYLRAWWNGYGAGFVDPETAAKMIR<br>EGAKLLSKIVPKELADEYEKAAIKGAGADQEAGAKILTPEERAATHACFAAAVDEVIAELKK | 29910 |
| 43 | SREEVARLAGRALELDVRLVEDLRAADPEKAEAWLEGARVPGPPVSNVTRLANAALGMLFEAAYVKL<br>KVGGISMEEALDRLYRLQLDWALAASVEELIALGREAGKADLAACYRADQILAADPSIIPDPARAEAIAA<br>RALKGEGIDEGVALLATLPADVLIELGFTHARVGLERNYGANEEELEKARELGTAMIRELTAP | 16960 |
| 44 | SPALAAALAALAALKKRVDARVAERLAKIDPAEWARAAAPIGRRLARLVAELGFPQEVVDAMLADPAR<br>AADVLGEAVIRALAEAVRRRLAALGLAELGEVVAAFLVEEARVLAQELALLFEDAYHAGASLDDLYARA<br>EELAAALRAAAPDELTGLIRRAAVLSGLLHALVGIDVEEGGKAWPYIERTVRELYRLYQIATA | 18450 |
| 45 | DDAAAQAVIDAILTEGGALWAKYPELWEAVLDAHALAGALLDRAAAKTGKPLLDITPEEETPELREAKA<br>ILSGTDIRVDRLIEKYLPGKAFLFRGPLYLAGSVKAIELGKLREWLGNILDAYVPELRALGLDDADLEALT<br>AELVRLFTLAAQDPRVLALSKEILEKEEAIQAEAAGLTDEEIRARLEEALARVEEILSKI | 22460 |
| 46 | PPLPPFGPPELEERVREELERAREVGIKVGNAVEGEMVARGGYTEEEVDKLVKEGADLIRNATSVEEI<br>GIGMGMIHAGATGYAYLLLPEEKRPGLLALTAFGIAATAAVGPEAAAALMRAAFPETVAAWEEVAPAV<br>LDKVLETIGVDADTAAVLREIDAATRGRGVDVLIEALARQALLRLHGVEVSLKEVVEALKRELL | 9970 |
| 47 | MKERVARLAALALERRHAMVARLAAADPAKAAAWAAGAAVPGPAVSATTRLANAALGNLFEAAYIKL<br>KVGGITMEEALDIHFDELLKWALAASLEELIAIGREAGRADREACARADRILAADPSIIPDPARAEAIAAR<br>ALRGEGIAEGVALLSTLPAEVLIELAFTHAKVGLERIYGANEEELKKARELGTEMIKTLTAP | 13980 |
| 48 | GAGAALGRRLLGAILGGDPREGLALAKRLYEGLWMLGVDPERGEAILAEVRAATGLDAEIDALAAVMR<br>ALNATIIAADEELRAADPAAYAAVLTFADLVLTSKLYPEKVSPEEKKAAELFLKGEELIAEYLRGGPESY<br>LTKGKEYVAEAAKILGLDPAQAWRGAVVHVAGSVARTPEQAAALAALDRTFEEALAAAEARL | 19940 |
| 49 | GLGELLGQLGAILRAGLANLVAHPERAAVIRDTDAANLARLRATGNQRFVDLVQRFYLSVFNAMLRAF<br>GWTREEGEVWLAVAGLDPALLDRLPDTISDENLAYFLSVELVVGCTHALGGDPEEVRALGRRLFRDW<br>LEFASDAPIVQELLATFTPEELELFLDGQVDHAVAGSVAFPDRAAVEAALRVGEAGVAYARSLAA | 20970 |
| 50 | GVGEAGRRLLGLYLKGLEVTIEHADEYIKYLPIHVEGAEKGTALYEAGDYVGATRTIIEADRKIAELIGNE<br>EAVALFDELLAAAAERAELRRALYAEAGLDDADVVAILRAGVAYDRGLLTLSPEEAAAALRADTQAEID<br>TLRARLEGERPLSPEELRRLAAFVAAWILGGGAATAIFDPELYREMLAAARAFYEALQRA | 18910 |
| 51 | GAGLAARLGARGARMGAWLEEHRDEVLAKTFEVERKLARELLAEAEAAGDARAAAVWRAYLEVLDW<br>LADPENLALWERVAAATPDLVVSRAVDALAVGVLAVICLDMSDEEIEERKDELIRIYAIIHAAQAVAVLET<br>VVATPELADRWVGMLSLSPEERAREIVRIWAEASNSPEGRRRIGERILAILKEIAEKLEAEKK | 35980 |
| 52 | GLGEAALGVIRGLDAGHRTLIETSEEGAQYVDDWYAVLLRALDRLGLGAVNDEVLAAIKRHGLYAITEV<br>VLGTLSPEQIQKAVDLGPLHAEAGLKLLEAVHGGASPEEVAAAQDAVADVDAEVMAQLRPLISEAQVE<br>AVAEALGVSREAALALILLTLAIYGGTVAEVRLLVEGTPEQRARVAGLLAQSAAALAALLAR | 11460 |
| 53 | GAGLAGGAAAAAGLAGALAAIYSTPEGKLVHRLWLTGAVLARDEVLAVADEAAAFARRAVDEKLDDE<br>TIAREAGELHARAAVVILRAVGLDAARLAAYRRVQAAMIAALLAAADAYAAAGLDFRALYLRAVVEAEL<br>AREPELDRIPVGDAPLLPITPEAEAIAFAADRYIDALIDYRVAARVDPALGEAARAALLALVDA | 14440 |
| 54 | GAGEGVRRRLGLLVVGARGYARLTPEEQRLIGVHGIAGMKIIEEKGLPDTEEEWIETGGTRELYEAWK<br>ATLARLLAALPPPDELDEAAVERYLAILSEFLREQAPLIVGGDEKAGYPHAELEAELAADPAAWRARVE<br>RLFARADEEFRLGTPEAFERAIKYFLTGLEAGLGVAGLAGPEAVAALCARAAERAQAWLDSLR | 29450 |
| 55 | AARAAGFDAIAATPEGRRRFFELLLRLPLVYDLLMAELLLARTEGASPEEQAELRAAADNLRAHFARST<br>SDPAAAEAAVLPLVDLLLEILAPYRDTPEFDEAVEAALTIARTAPAFDAATPAAVRALRVAAGDAFTAAA<br>RALNAGDLDTARVHVDAGLALAAQSWAANRAAGLGPQVDTLLAASKVISAFVRKAVEEYLA | 9970 |
| 56 | SAAALAAAAAGVAALGERRAAIVAAYRGGYAADMKIFGFTEAEVRELAERIRRPPFTGFTPEQLGQILL<br>GTAAAQDAWDKVVEAAGSYEAVARAYWARRVEEVGADEEEARQAGLIHAALSLVAAELYATLSPEEA<br>VLTLLSTRYYACLDALKEYGKKVFLLLSGDFLTAGAARLLVEDPSLNLDEATARTREIIESIRK | 22920 |
| 57 | ADLATGLERINATPEGRRRFFELLLRLPLVYDLLMAALLLRRTEGASPEERARLTAAADNLLAHFARST<br>SDPERARAAVQPLVDVLLAILKPYEDTPEMDKAIEAALTIADTAPAFDAATPASVRALRVAAGDAFTAA<br>AEALNAGDLDTAEVHVRAGLALAAQSWAANRAAGLQPQVDTLLAAVEVITAFVRKAVEDYLA | 9970 |

| | | |
|---|---|---|
| 58 | GPGKEIGEKLLELYEETVRLMRAEPGDPRSICLYDIVFHSLLIVLDPRLAPITEPLIALNQAALDAGTAEA AAAAHIETSKRLPEIYRAAVDDPATSEDALRFLIEFTYYWAKVGSSLPSARALVEALAPVIKKIDAGTAT REELRAALGAALPLIEARIAEGVAADEAATPEEVCLGFDKVFLAVLELRDVLREYIENL | 14565 |
| 59 | SEEARRRREEGIRRLEERREEIVASYLGGLKADMAIFGFTDEEVRALAEKIRRPPFTGFTPEQLGQILL SEADAFEEWKKVVEKYGSYEAVARAYWAKKVKEVGANEEELRQAGLVHAALSLVAAELFATLSPEEA VLTLLSTRYYACLDALRKYGKKVFLLLSDDFLRAGAARLLVEDPSLNLEEARARTQEIIDAIRK | 21430 |
| 60 | GEGKKIGEELLERYEEMVRLMREQPGHPASITMYDIVFNRLLTVLDPRLAPITEPLIAENEAALEAGTAE AAAAAHIANSRRLPDIYRAAVDDPATSERALEFLIRFTYYWAKVASALPAARALVEALAPVIAKIDAGTA TREELRAALAAALPLIEAVIAEGVAADEAATPAEVCAGFDRTFLAVLEERDVLREFIENL | 12950 |
| 61 | AAALAAEIGAAAAEFAQLSLDEQIAAAIAKQQAWLDFVNSHTYEEIKALAAELAKDPRAAALVDLLRLAE VLARAVAARVRDGEAGRERLRKVLETATQYVIENSKPIHDKLVAEESIFPNYADFQRYRLEKELAILDA PTAEAAVDAVLDAIRDSLKVYKGPFHRTGLEYLVAHEAEVRAFLRAVADAAAAATRARLPR | 14440 |
| 62 | ALAALLARAAGIAALGAARAAIVAAYRGGLAADMAIFGFTEAEVEALARRIRRPPFTGFTPEQLGQILLG TAAFQAAWQALVDAAGSYEAVARAYWERKVAEVGANEEELRQAALIHAALSLVAAELFATLTPEEAVL TLLSTRYYACLDALKEYGPKVFLLLSDDFLRAGAARLLVEDPSLNLEEATARTREIIEAIRK | 19940 |
| 63 | SPALEAALEAAGPRAAAAIERLRPLFDYLVTISKADIDPNLPAEERERIAKAIDDFFYGRPGFALAAQGV WFRHMAEITPEGPTREAYRAGAEGVLRILRGEDLEAGLEGLRRGLEEVVRVNAKEEALEPELRKELET VTGEEVVARAERLFAEGRIVEAGATLHTGAQHPKTAPVASKWLRDRNAALLERGFAELAGVF | 15470 |

**Table S7**:

Conversion ± standard deviation of substrates 2-cyclohexenone and 4-nitrobenzaldehyde after 8 hours with 2 mol% of enzyme or small molecule catalyst. Designs with identical backbone numbers have the same fold. Relative conversions denote conversions relative to the construct with maximum conversion, MBH48. AS model indicates the active site model the design was based on. Fragment picking indicates if an idealized helical backbone fragment was used as scaffold (no) or if fragments were picked from the PDB (yes). Final CM refinement indicates if the Rosetta Coupled Moves protocol was used to generate final active site sequences instead of LigandMPNN.

| MBH | backbone number | conversion [%] | relative conversion [%] | AS model | fragment picking | final CM refinement |
|---|---|---|---|---|---|---|
| 1 | 1 | 2.68 ± 0.57 | 16.77 | | | |
| 2 | 1 | 1.14 ± 0.2 | 7.12 | | | |
| 3 | 2 | 1.80 ± 0.4 | 11.25 | | | |
| 4 | 2 | 0.77 ± 0.18 | 4.84 | | | |
| 5 | 3 | 1.32 ± 0.24 | 8.26 | | | |
| 6 | 3 | 2.67 ± 0.69 | 16.71 | | | |
| 7 | 4 | 1.00 ± 0.17 | 6.28 | | | |
| 8 | 5 | 1.05 ± 0.14 | 6.57 | | | |
| 9 | 5 | 1.46 ± 0.22 | 9.15 | BH32.14 | no | no |
| 10 | 6 | 1.88 ± 0.23 | 11.76 | | | |
| 11 | 7 | 1.56 ± 0.11 | 9.73 | | | |
| 12 | 8 | 1.52 ± 0.2 | 9.52 | | | |
| 13 | 9 | 1.43 ± 0.15 | 8.95 | | | |
| 14 | 10 | 1.81 ± 0.3 | 11.33 | | | |
| 15 | 11 | 2.4 ± 0.39 | 14.98 | | | |
| 16 | 12 | n. d. | n. d. | | | |
| 17 | 13 | 1.32 ± 0.15 | 8.25 | | | |

| | | | | | | |
|---|---|---|---|---|---|---|
| 18 | 14 | 3.01 ± 0.48 | 18.83 | | | |
| 19 | 15 | 4.78 ± 0.65 | 29.91 | BH1.8 | yes | no |
| 20 | 16 | 1.51 ± 0.22 | 9.45 | | | |
| 21 | 17 | 1.12 ± 0.04 | 6.98 | | | |
| 22 | 18 | 2.17 ± 0.23 | 13.55 | | | |
| 23 | 17 | 0.53 ± 0.01 | 3.33 | | | |
| 24 | 18 | 2.1 ± 0.16 | 13.15 | | | |
| 25 | 15 | 0.61 ± 0.12 | 3.80 | | | |
| 26 | 18 | 5.31 ± 0.82 | 33.18 | | | |
| 27 | 19 | 1.7 ± 0.12 | 10.63 | | | |
| 28 | 19 | 1.43 ± 0.04 | 8.95 | | | |
| 29 | 20 | 4.4 ± 0.42 | 27.51 | | | |
| 30 | 21 | 0.92 ± 0.16 | 5.78 | | | |
| 31 | 22 | 2.82 ± 0.14 | 17.60 | | | |
| 32 | 23 | 1.6 ± 0.14 | 10.03 | | | |
| 33 | 21 | 1.26 ± 0.03 | 7.90 | | | |
| 34 | 23 | 1.58 ± 0.05 | 9.85 | | | |
| 35 | 16 | 1.67 ± 0.13 | 10.45 | | | |
| 36 | 24 | 4.11 ± 0.16 | 25.68 | | | |
| 37 | 25 | 3.41 ± 0.07 | 21.33 | | | |
| 38 | 26 | 1.05 ± 0.09 | 6.55 | | | |
| 39 | 27 | 1.62 ± 0.12 | 10.13 | | | |
| 40 | 25 | 1.27 ± 0.03 | 7.95 | | | |

| | | | | | | |
|---|---|---|---|---|---|---|
| 41 | 24 | 2.05 ± 0.1 | 12.83 | | | |
| 42 | 26 | n. d. | n. d. | | | |
| 43 | 28 | 1.9 ± 0.02 | 11.85 | | | |
| 44 | 20 | 3.51 ± 0.22 | 21.93 | | | |
| 45 | 29 | 4.07 ± 0.19 | 25.43 | | | |
| 46 | 30 | 7.06 ± 0.38 | 44.16 | | | |
| 47 | 28 | 9.46 ± 0.68 | 59.16 | | | |
| 48 | 18 | 16 ± 0.57 | 100.00 | | | |
| 49 | 16 | 2.34 ± 0.47 | 14.63 | | | |
| 50 | 27 | 1.78 ± 0.1 | 11.13 | | | |
| 51 | 24 | 1.47 ± 0.05 | 9.18 | | | yes |
| 52 | 22 | 0.34 ± 0.07 | 2.10 | | | |
| 53 | 31 | 1.19 ± 0.06 | 7.43 | | | |
| 54 | 32 | 3.18 ± 0.06 | 19.90 | | | |
| 55 | 33 | 3.22 ± 0.95 | 20.13 | | | |
| 56 | 34 | 10.94 ± 0.89 | 68.37 | | | |
| 57 | 33 | 5.83 ± 0.74 | 36.43 | | | |
| 58 | 35 | 0.55 ± 0.12 | 3.45 | | | |
| 59 | 34 | 1.8 ± 0.1 | 11.28 | | no | no |
| 60 | 35 | n. d. | n. d. | | | |
| 61 | 36 | 4.86 ± 0.69 | 30.41 | | | |
| 62 | 34 | 2.86 ± 0.14 | 17.90 | | | |
| 63 | 37 | n. d. | n. d. | | | |

| | | | | | | |
|---|---|---|---|---|---|---|
| lysozyme | n.a. | 0.26 ± 0.09 | 0.00 | n.a. | n.a. | n.a. |
| DMAP | n.a. | 0.18 ± 0.02 | 0.00 | n.a. | n.a. | n.a. |
| imidazole | n.a. | 0.14 ± 0.02 | 0.00 | n.a. | n.a. | n.a. |

**Table S8.**

Comparison of designed and evolved enzymes for the Morita-Baylis-Hillman reaction of 2-cyclohexenone and 4-nitrobenzaldehyde according to a random-order fit. ± indicates 95% confidence interval of the fit.

| | kcat [$min^{-1}$] | $K_{M, cyclohexenone}$ [mM] | $K_{M, 4\text{-}nitrobenzaldehyde}$ [mM] | Ref. |
|---|---|---|---|---|
| **Computationally designed MBH enzymes (this work)** | | | | |
| MBH18 | 7.70E-03 ± 1.9E-03 | 8.33 ± 2.97 | 8.55 ± 3.73 | this work |
| MBH48 | 2.49E-02 ± 5.9E-03 | 25.16 ± 10.6 | 1.42 ± 0.35 | this work |
| **Computationally designed MBH enzymes (from existing scaffolds; no laboratory evolution)** | | | | |
| BH32 | 2.23E-03 | 7.998 | 1.77 | [11, 12] |
| **Computationally designed MBH enzymes (from existing scaffolds; laboratory evolution)** | | | | |
| BH32.8 | 1.68E-02 | 4.173 | 2.381 | [12] |
| BH32.12 | 1.00E-01 | 9.719 | 0.893 | [12] |
| BH32.14 | 3.49E-01 | 2.556 | 1.137 | [12] |
| BH1.8 | 4.50E+00 | 12.02 | 0.323 | [13] |
| BH1.8 23H | 1.13E+00 | 12.19 | 0.293 | [13] |

**Table S9.**

Primers used for generation of RAD variants.

| RAD | variant | forward primer | reverse primer | Ta [°C] |
|---|---|---|---|---|
| 1 | K123A | TGGTGTAGGTgcgTATGAGGGTATG | AGGAAAAAAGCAAGTGAG | 58 |
| 2 | K122A | GGGAATCGGAgcgGCGAAAGGAG | AAGAAGAACGCCAAAC | 57 |
| 3 | K119A | TGGCGGAATTgcgGGTGTTGCTG | ACGGCGATCGTCAATG | 63 |
| 4 | K119A | ATTGGGGATTgcgGGAGTTGCTG | ACCGCGATAGCCAATG | 60 |
| 5 | K119A | TTCAGGAATTgcgGGCGTAGCTG | ACGGCAATGGTTAAAG | 57 |
| 6 | K119A | AGGAGCCGTAgcgGGCACGCGG | AAACCGATGGTCAGAGCTAATTC | 66 |
| 7 | K177A | CTCCATCATCgcgGGTTTGGCGTTTG | AAGCCTAACACCAATGC | 61 |
| 8 | K22A | CTTCGGCGCTgcgGGACTGCTTAATC | ATAGCCAGCAGGCGT | 64 |
| 9 | K65A | TGCTGCCGCTgcgGGGCTGGTTT | ATGATGGTCTCATCCGATAATTTCC | 66 |
| 10 | K65A | CGCAGCGGCAgcgGGTTTAACTTAC | ATAATAGCATCATCGCTAAGTTTC | 62 |
| 11 | K65A | CGCGGCAGCCgcgGGCTTGGCGT | ATAATTGCGTTGTAGCTTAACTTCCC | 68 |
| 12 | K65A | TGCCGCCGCAgcgGGGCTTGTAT | ATAATTGTTTCGCGCGAC | 62 |
| 13 | K65A | TGCCGCGGCTgcgGGTCTTGCTT | ATAATCGCGTCATCGGAAAG | 64 |
| 14 | K70A | TGCGCTTGTTgcgGGGATCGCTC | GCCAGCACCAGTGCT | 66 |
| 15 | K70A | TGCTTTGGTGgcgGGTATGGCCC | GCTAATACTAACGCGGC | 63 |
| 16 | K70A | AGTTCTGGTGgcgGCCATGGCATTATA | GCCAGGACAAGAGCG | 64 |
| 17 | K81A | GTCCCTGGTCgcgGCTACGGCCA | AGAAGTACAAGAATCTTTTTTACGACTTC | 64 |
| 18 | K81A | GGCCCTTATCgcgGCCACCGCAATTTAC | AAAAGGATTAAAATCTCCTCGATC | 61 |
| 19 | K81A | GGCGTTAGTAgcgGCTACAACTATCTAC | AGCAGTACCAGCAAG | 59 |
| 20 | K81A | AGCATACATCgcgGCAGCGACGATC | AACAAAATCAACATCTTCTTTG | 57 |
| 21 | K81A | AGCATACATTgcgGCAACCGCCATTAAG | AAAAGGATTAACATCTTGCG | 59 |
| 22 | K122A | AGTAATCGCGgcgGGGGCGTATTTTGATC | GTCAGGGCAGCGGTG | 65 |
| 23 | K122A | AGTGATCGCTgcgGGCTCTTACAAG | GTGATCGCATTAGTCAC | 58 |
| 24 | K122A | TGTTATCGCAgcgGGTGCCGTGT | GTTAAAGCGTTCGTTACC | 60 |
| 25 | K62A | CGCCACACTTgcgGGGGTGCTTG | GTGATGACGGTAAAGAATC | 59 |
| 26 | K76A | TGTCGGGGCGgcgGGATACGTCG | ATAACAGCGAATGCGGC | 65 |
| 27 | K62A | ACTTTCGTTGgcgGGGGTTCTTGCAG | GTGAACACTGTAAAGAACTCATC | 62 |
| 28 | K62A | ATACACCCTGgcgGGGGTGCTTG | GTAATTACAGTAAAGAACTCCC | 60 |
| 29 | N178A | GCGTAAGTTCgcgGAGGAAGCTGG | ATTTCCTCCACCTGTTC | 58 |
| 29 | K76M | TGTCTACGCTatgGGCTTAGCAC | ATGACAGCAAACGCGG | 64 |
| 29 | Y23A | TGCTCAGCCGgcgTATGATTTTGTCAAAAAG | TTCTCCTTCGCAGCTTC | 58 |
| 29 | Y124A | TCCGAAAAAAgcgTTTGGCGCAATCAC | TCCTTGATCAACTTCTTG | 56 |
| 29 | K76A | TGTCTACGCTgcgGGCTTAGCAC | ATGACAGCAAACGCG | 61 |
| 30 | K76A | TGTTGGAGCGgcgGGATTTGCAG | ATAACAGCGAAAGCC | 57 |
| 31 | K62A | CATGTTGTTGgcgGGTCTTTTCTATCAC | GCCAATACAAGCAAGTC | 57 |
| 32 | K62A | CATGCTGTTAgcgGGTATCTTTTATCACG | GCTAATACCTCCAAGTC | 57 |
| 32 | K62M | CATGCTGTTAatgGGTATCTTTTATCAC | GCTAATACCTCCAAGTC | 56 |
| 32 | Y120F | TGCACCGCAAtttTTAACCCTTATC | AATTCCTCCCAGGAAATAC | 60 |
| 32 | Y99F | GTCGCTTTTGtttGCGGAACTTC | ATAAGACGCTGTTCCAAG | 60 |

| | | | | |
|---|---|---|---|---|
| 32 | N178A | AGAACCGATCgcgAAAAAGATGGGAGAATAC | ACCTCTTTGGTCGCTTC | 61 |
| 34 | K69A | AATCGCAATCgcgGGCGCTGTCTTCTCC | GCCAACGCGATGCCA | 65 |
| 35 | K62A | CATGTTTGTCgcgGGGATCTTTTTTCATG | GCCAGAACGTCCATATC | 56 |
| 35 | K62M | CATGTTTGTCatgGGGATCTTTTTTCATG | GCCAGAACGTCCATATC | 58 |
| 35 | Y23F | TGTAAAGGAAtttGTGGAGCGTG | ATTTTTTCACGATTTTTGATAGC | 58 |
| 35 | Y120F | TTGGCCGTTAtttTTATCCCTTATC | AACTCTTCACGGCTG | 59 |
| 35 | N178A | TGCACCTATCgcgGAAGAAGCTGC | ACTTCTTCGGTGGCC | 58 |
| 36 | K69A | AATCGGCATGgcgGGCCCGATCGAG | GCTAAAGCAATTCCTAAGGCTTT | 65 |

**Table S10.**

**Crystallization conditions and diffraction parameters.** Values in the left column of the respective crystal structure denote overall parameters, values in the right column belong to the highest resolution shell.

| construct | RAD13 | | RAD17 | | RAD32 | | RAD36 | | MBH2 | | MBH48 | |
|---|---|---|---|---|---|---|---|---|---|---|---|---|
| **PDB ID** | 9GBT | | 9FW5 | | 9FW7 | | 9FWA | | 9QDP | | 9R7F | |
| **screen name** | Hampton Research JCSG Plus | | Hampton Research JCSG Plus | | Jena Bioscience JBScreen PACT++ | | Hampton Research JCSG Plus | | Hampton Research Index | | Molecular Dimensions SG1 | |
| **crystallization conditions** | 50% PEG400, 0.1 M sodium acetate, pH 4.5 | | 2.1 M D/L malic acid, pH 7.0 | | 25% PEG1500, 0.1 M MIB, pH 4.0 | | 20% PEG8000, 0.1 M CHES, pH 9.0 | | 240 mM $NH_4CH_3CO_2$, 25% PEG 3350, 100 mM BIS-TRIS pH 5.5 | | 0.2 M $(NH_4)_2SO_4$, 0.1M Bis-Tris pH 6.5, 25% PEG 3350 | |
| **Beamline** | ESRF ID30A-3 | | ESRF ID30B | | ESRF ID30A-3 | | ESRF ID30A-3 | | ESRF ID30A-3 | | ESRF ID23-2 | |
| **DOI** | 10.15151/ESRF-ES-1476096327 | | 10.15151/ESRF-ES-1473914182 | | 10.15151/ESRF-ES-1413581206 | | 10.15151/ESRF-ES-1413581206 | | 10.15151/ESRF-ES-2006582544 | | 10.15151/ESRF-ES-2015886544 | |
| **Wavelength** | 0.9677 | | 0.9677 | | 0.9677 | | 0.9677 | | 0.9677 | | 0.8731 | |
| **Resolution range** | 59.61-2.43 | 2.62-2.43 | 47.08-2.9 | 3.004-2.9 | 32.45-2.0 | 2.2-2.0 | 36.67-1.73 | 1.792-1.73 | 36.75-1.13 | 1.15-1.13 | 44.21-1.93 | 2.08-1.93 |
| **Space group** | P 1 21 1 | | I 41 | | P 31 | | P 21 21 21 | | C 1 2 1 | | P 63 | |
| **Unit cell** | 38.044 76.936 59.619 90 91.23 90 | | 107.108 107.108 52.415 90 90 90 | | 60.51 60.51 41.32 90 90 120 | | 46.6 55.932 73.336 90 90 90 | | 56.775 51.881 72.606 90 112.2790 | | 76.056 76.056 59.638 90 90 120 | |
| **Total reflections** | 107876 | 20065 | 69818 | 7184 | 164833 | 39859 | 273610 | 28523 | 494274 | 4122 | 308129 | 61995 |
| **Unique reflections** | 15039 | 2922 | 6710 | 645 | 15322 | 3822 | 20599 | 2028 | 157994 | 2124 | 14863 | 2948 |
| **Multiplicity** | 7.2 | 6.9 | 10.4 | 11 | 10.8 | 10.4 | 13.3 | 14.1 | 3.1 | 1.9 | 20.7 | 21 |
| **Completeness (%)** | 98.94 | 98.88 | 99.55 | 98.77 | 99.89 | 99.97 | 99.86 | 100 | 96.77 | 93.17 | 98.24 | 91.49 |
| **Mean I/sigma(I)** | 4.54 | 0.47 | 11.89 | 0.89 | 5.7 | 0.72 | 15.26 | 2.25 | 9.83 | 1.12 | 13 | 0.65 |
| **Wilson B-factor** | 57.89 | | 25.96 | | 32.06 | | 22.43 | | 11.64 | | 41.85 | |
| **R-merge** | 0.221 | 2.641 | 0.111 | 2.692 | 0.272 | 2.989 | 0.1071 | 1.316 | 0.05517 | 0.8954 | 0.145 | 2.226 |
| **R-meas** | 0.2384 | 2.859 | 0.1174 | 2.826 | 0.2854 | 3.141 | 0.1115 | 1.365 | 0.06582 | 1.227 | 0.1486 | 2.281 |
| **R-pim** | 0.08844 | 1.081 | 0.03735 | 0.8503 | 0.086 | 0.9575 | 0.03069 | 0.3609 | 0.03543 | 0.8316 | 0.03266 | 0.4964 |
| **CC1/2** | 0.996 | 0.46 | 0.997 | 0.525 | 0.996 | 0.566 | 0.998 | 0.796 | 0.996 | 0.407 | 1 | 0.756 |
| **CC*** | 0.999 | 0.794 | 0.999 | 0.83 | 0.999 | 0.85 | 0.999 | 0.941 | 0.999 | 0.761 | 1 | 0.928 |
| **Reflections used in refinement** | 12940 | 2566 | 6697 | 645 | 11427 | 2866 | 20606 | 2028 | 70627 | 2675 | 14602 | 2697 |
| **Reflections used for R-free** | 652 | 128 | 302 | 33 | 572 | 144 | 1020 | 84 | 3520 | 111 | 749 | 141 |
| **R-work** | 0.2236 | 0.3291 | 0.2349 | 0.3205 | 0.2162 | 0.2594 | 0.1903 | 0.2791 | 0.1389 | 0.3319 | 0.2011 | 0.385 |
| **R-free** | 0.267 | 0.3496 | 0.2702 | 0.5281 | 0.2717 | 0.3375 | 0.2341 | 0.3419 | 0.1675 | 0.3329 | 0.2514 | 0.4967 |
| **Number of non-hydrogen atoms** | 3206 | | 1499 | | 1542 | | 1710 | | 2036 | | 1649 | |
| **macromolecules** | 3134 | | 1477 | | 1503 | | 1528 | | 1601 | | 1497 | |
| **ligands** | 47 | | 22 | | 0 | | 2 | | 15 | | 64 | |

| | | | | | | |
|---|---|---|---|---|---|---|
| **solvent** | 25 | 0 | 39 | 180 | 420 | 88 |
| **Protein residues** | 400 | 197 | 199 | 202 | 206 | 202 |
| **RMS(bonds) (Å)** | 0.003 | 0.011 | 0.002 | 0.007 | 0.003 | 0.006 |
| **RMS(angles) (°)** | 0.53 | 1.41 | 0.53 | 0.73 | 0.7 | 0.78 |
| **Ramachandran favored (%)** | 99.49 | 98.97 | 97.97 | 99.5 | 99.51 | 99 |
| **Ramachandran allowed (%)** | 0.25 | 1.03 | 2.03 | 0.5 | 0.49 | 1 |
| **Ramachandran outliers (%)** | 0.25 | 0 | 0 | 0 | 0 | 0 |
| **Rotamer outliers (%)** | 0.63 | 0.72 | 0 | 0.68 | 0 | 0 |
| **Clashscore** | 5.15 | 7.76 | 3.35 | 0.33 | 0.93 | 2.85 |
| **Average B-factor** | 66.79 | 67.25 | 33.33 | 24.05 | 18.87 | 52.32 |
| **macromolecules B-factor** | 66.76 | 67.82 | 33.25 | 22.86 | 14.71 | 51.58 |
| **ligands B-factor** | 73.2 | 29.35 | | 28.51 | 37.57 | 68.58 |
| **solvent B-factor** | 58.42 | | 36.26 | 34.08 | 34.07 | 53.16 |
| **number of TLS groups** | | 5 | | | | |
